# Impact of industrial production system parameters on chicken microbiomes: mechanisms to improve performance and reduce *Campylobacter*

**DOI:** 10.1101/2020.05.10.084251

**Authors:** Aaron McKenna, Umer Zeeshan Ijaz, Carmel Kelly, Mark Linton, William T. Sloan, Brian D. Green, Ursula Lavery, Nick Dorrell, Brendan W. Wren, Anne Richmond, Nicolae Corcionivoschi, Ozan Gundogdu

**Affiliations:** Moy Park, 39 Seagoe Industrial Estate, Portadown, Craigavon, Co. Armagh, BT63 5QE, UK; School of Engineering, University of Glasgow, Glasgow, G12 8LT, UK; Agri-Food and Biosciences Institute, Food Microbiology, Newforge Lane, Belfast, BT9 5PX, UK; Institute for Global Food Security, School of Biological Sciences, Queen’s University Belfast, Biological Sciences Building, Belfast BT9 5DL, Northern Ireland; Faculty of Infectious & Tropical Diseases, London School of Hygiene and Tropical Medicine, Keppel Street, London, WC1E 7HT, UK

**Keywords:** Chicken, Microbiome, *Campylobacter*, Production Systems, Environmental Filtering, Phylogenetic Signal, Competitive Exclusion, Diversity

## Abstract

**Background:** The factors affecting host-pathogen ecology in terms of the microbiome remain poorly studied. Chickens are a key source of protein with gut health heavily dependent on the complex microbiome which has key roles in nutrient assimilation and vitamin and amino acid biosynthesis. The chicken gut microbiome may be influenced by extrinsic production system parameters such as *Placement Birds/m*^*2*^ (stocking density), feed type and additives. Such parameters, in addition to on-farm biosecurity may influence performance and also pathogenic bacterial numbers such as *Campylobacter*. In this study, three different production systems ‘Normal’ (N), ‘Higher Welfare’ (HW) and ‘Omega-3 Higher Welfare’ (O) were investigated “in a natural environment” at day 7 and day 30 with a range of extrinsic parameters assessing performance in correlation with microbial dynamics and *Campylobacter* presence.

**Results:** Our data identified production system N as significantly dissimilar from production systems HW and O when comparing the prevalence of genera. An increase in *Placement Birds/m*^*2*^ density led to a decrease in environmental pressure influencing the microbial community structure. Prevalence of genera such as *Eisenbergiella* within HW and O, and likewise *Alistipes* within N were representative. These genera have roles directly relating to energy metabolism, amino acid, nucleotide and short chain fatty acid (SCFA) utilisation. Thus, an association exists between consistent and differentiating parameters of the production systems, that affect feed utilisation, advance our knowledge of mechanistic underpinnings, leading to competitive exclusion of genera based on competition for nutrients and other factors. *Campylobacter* was identified within specific production system and presence was linked with the increased diversity and increased environmental pressure on microbial community structure. Addition of Omega-3 though did alter prevalence of specific genera, in our analysis did not differentiate itself from HW production system. However, Omega-3 was linked with a positive impact on weight gain.

**Conclusions:** Overall, our results show that microbial communities in different industrial production systems are deterministic in elucidating the underlying biological confounders, and these recommendations are transferable to farm practices and diet manipulation leading to improved performance and better intervention strategies against *Campylobacter* within the food chain.

**Graphical Abstract:** 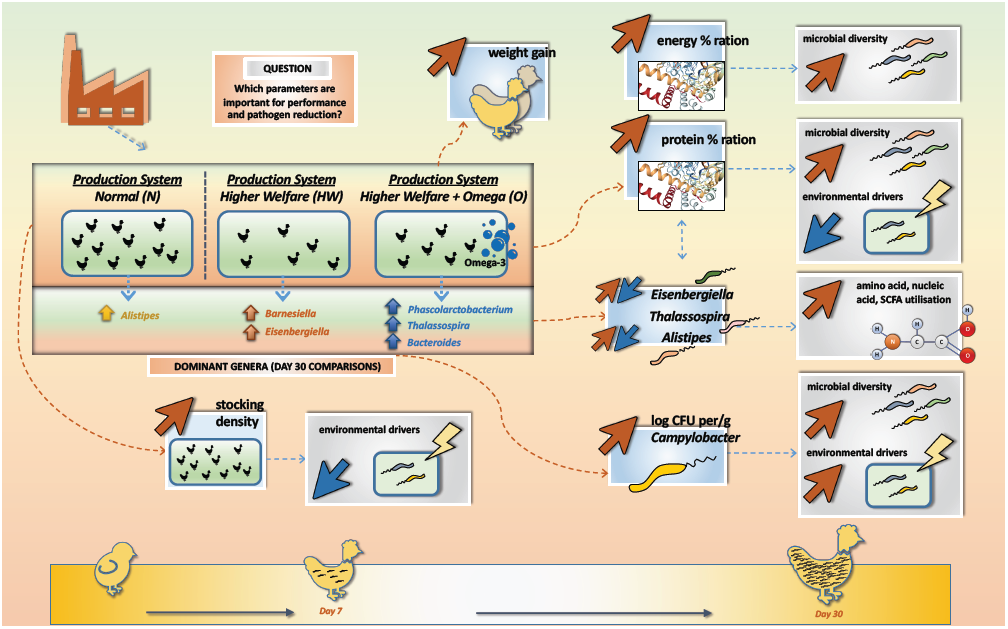

## Background

Chickens are a key source of protein for humans where poultry production is predicted to produce approximately 130 million tons of chicken meat in 2020 [1, 2]. Sustainable poultry practices are needed to help maintain an adequate supply of poultry products for the increasing human population without compromising the chicken or human health [3]. Selective breeding programmes has resulted in chickens that efficiently convert food into body mass, as defined by feed conversion ratios (FCR) [4]. The fundamental component needed to ensure an efficient poultry production is highly dependent in having an optimised nutrition and production set up [1].

Production systems vary immensely between countries, businesses and at farm level. Certain parameters such as *Placement Birds/m*^*2*^ (stocking density), *Protein_perc_ration* (protein percentage within ration) and *Energy_of_ration* (energy content) in relation to the feed are key determinants that if varied, may directly influence chicken microbial community structure. Thus, impacting performance and potentially reduction of pathogenic bacteria. Chicken diets are typically formulated to enhance production efficiency. However, some diets are formulated to enhance human health, such as diets containing Omega-3 polyunsaturated fatty acids (PUFA’s) [5]. There is anecdotal evidence at farm level to suggest that this enrichment has potential to improve chicken gut health and performance or reduce pathogen colonisation.

Many bacterial species found within the microbiome of farmed animals can potentially be considered pathogens and detrimental to human health. One of these pathogens is *Campylobacter* which is the leading cause of human foodborne bacterial gastroenteritis causing bloody diarrhoea, fever and abdominal pains in humans and can also cause post infectious sequelae such as Guillain-Barré syndrome which is a potentially fatal paralytic autoimmune illness [6]. *Campylobacter jejuni* (predominant species causing infection to humans) colonises chicken ceca with relatively high numbers (>10^9^ CFU per gram) and can be pathogenic to the chicken, with this dependent on the genetics of host and strain of infection [7-10]. Although there exists many intervention studies, in an industrial farm environment, we currently do not understand why we typically see *Campylobacter* at approximately two weeks into the chicken life cycle [11-13]. The natural growth and flux of the gut microbiome may have a role to play [14]. This is further convoluted by the fact that there are very few studies on how chicken diet impacts *Campylobacter* presence within the chicken gut microbiome, particularly from within an industrial farm environment.

Chicken performance and gut health is heavily dependent on the largely unexplored complex gut microbial community which plays a role in nutrient assimilation, vitamin and amino acid biosynthesis and prevention of pathogen colonization [15-18]. The microbiota is responsible for hydrolysing indigestible carbohydrates and polysaccharides allowing further fermentation by other members of the gut ecosystem that produce SCFA, in turn allowing utilisation by the host [1]. The relationship between gut microbiota, chicken health and performance represents a tripartite that has been under the scrutiny of the research community with the prospect of improving the efficiency of current microbiome manipulation strategies [19, 20]. As an example, a xylanase gene from chicken cecum has been isolated and overexpressed, potentially leading to development of new feed additives for industrial application [21]. Our understanding of diet and its impact on the intestinal microbiota is still nascent and requires further exploration.

In the present study, we aim to build on our previous research where we performed a comprehensive analysis of the chicken cecal microbiome from days 3 to 35 investigating the driving forces of bacterial dynamics over time, and how this relates to *Campylobacter* appearance within a natural habitat setting [14]. Microbial variation over time was heavily influenced by the diet, where significant shifts in bacterial composition were observed [14]. The factors affecting host-pathogen ecology in terms of microbial community structure remain poorly studied at an industrial farm level. Therefore, extending our previous work and in view of the above ambitions, in this study, three different industrial production systems, namely, ‘Normal’ (N), ‘Higher Welfare’ (HW) and ‘Omega-3 Higher Welfare’ (O) were investigated at day 7 and day 30, along with extrinsic parameters (which were not available in our previous study) to ascertain mechanisms on improving the overall performance of chickens, and also elucidating the role of microbial community dynamics on revealing *Campylobacter* pathogenesis.

## Materials and Methods

### Ethics Statement

Euthanasia of birds was carried out under veterinary supervision alongside routine veterinary diagnostic inspection and after consultation and approval from the ethical committee within Moy Park.

### Experimental design, broilers and sample collection

Chickens reared under three different industrial growing regimes, ‘Normal’ (N), ‘Higher Welfare’ (HW) and ‘Omega-3 Higher Welfare’ (O), were sourced from three different contract farms supplying chicken to Moy Park (39 Seagoe Industrial Estate, Portadown, Craigavon, Co. Armagh, BT63 5QE, UK).

Although the three production systems differ in receiving chicks from multiple flocks, our analysis suggested that they had no bearing on microbial community structure at finer level (Fig. 3 and discussion later) although some parameters were implicated as significant in PERMANOVA analysis. All chicks were Ross 308 as hatched (AH) and were supplied from the same Moy Park hatchery to all three farms on 11/10/2018. All birds were grown in typical industrial poultry houses and were raised on a four-stage diet made up of a starter, grower, finisher and withdrawal ration, however composition of these diets differed across the three growing regimes. Farm N – Birds were offered standard starter ration from days 0 to 11, standard grower ration from days 11 to 22 and standard finisher from day 22 to day 34 before moving to a standard withdrawal ration prior to slaughter. Farm HW - Birds were offered higher welfare starter ration from days 0 to 11, higher welfare grower ration from days 11 to 23 and higher welfare finisher from day 23 to day 31 before moving to a higher welfare withdrawal ration prior to slaughter. Farm O - Birds were offered higher welfare starter ration from days 0 to 11, higher welfare grower ration from days 11 to 20 and Omega-3 finisher ration from day 20 to day 30 before moving to an Omega-3 withdrawal ration prior to slaughter. Both Omega-3 finisher rations are identical to HW equivalent ration, but with addition of an Omega-3 premix produced by Devenish Nutrition Ltd (Belfast, UK) and added at the feed mill. The change in dietary ration has influenced microbial community structure, as can be seen later in the analysis. At day 7, 19 chickens and at day 30, 10 chickens were randomly removed from the same single poultry house on each of the three farms.

**Figure 1:**
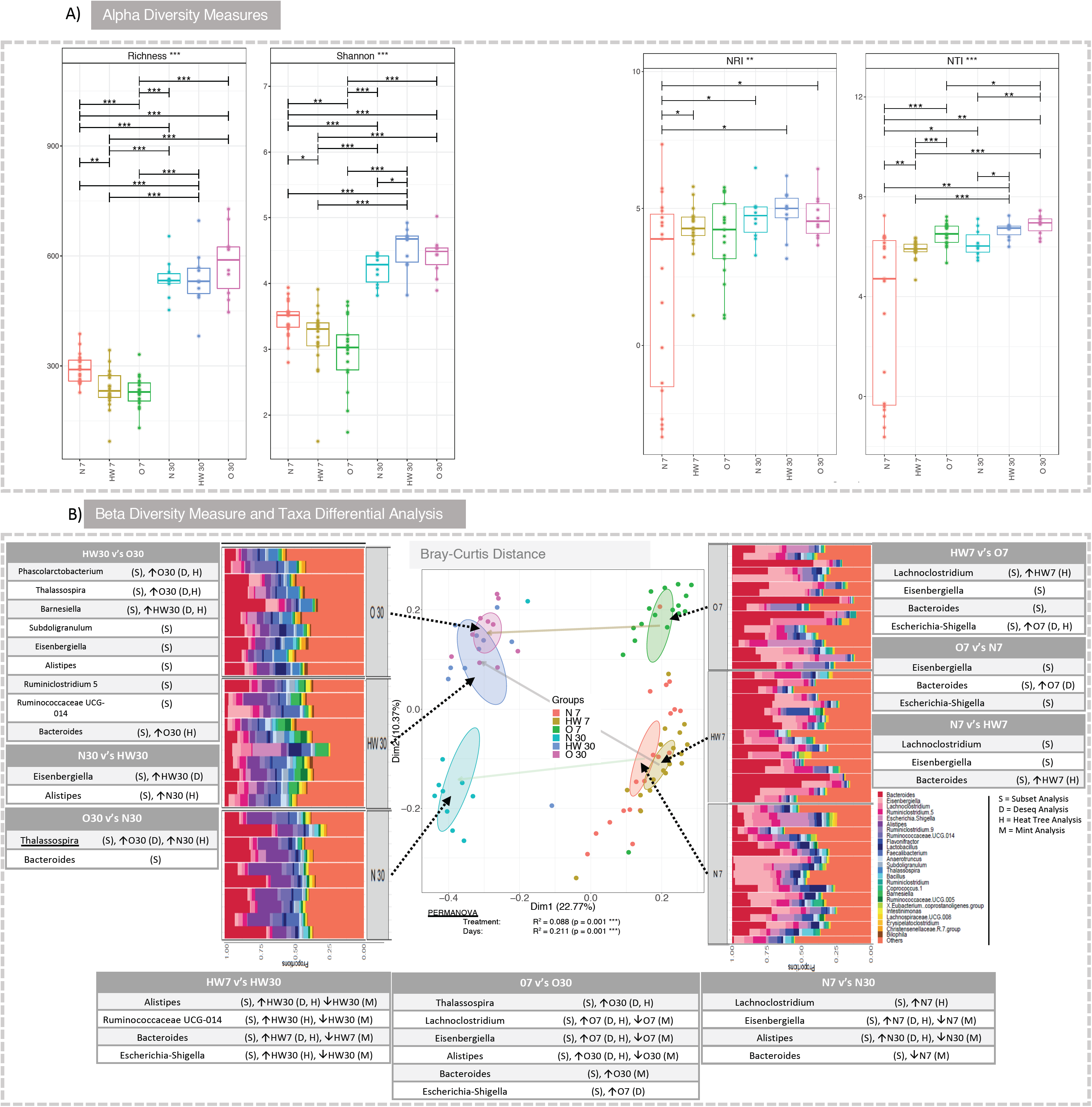
Microbial diversity and community structure. (a) represents alpha diversity (Richness and Shannon entropy) and environmental filtering (NRI/NTI) measures respectively. Lines for figure (**a)** connect two categories where the differences were significant (ANOVA) with * (P < 0.05), ** (P < 0.01), or *** (p < 0.001). (b) beta diversity using Bray-Curtis distance measure along with Top-25 genera observed in all samples grouped by categories. The tables represent taxa that were found to be significant based on Subset analysis (Supplementary S1) i.e., those genera selected in the subsets that explain roughly the same distance between samples as all the genera. Additionally, if the taxa were found to be differentially expressed based on other analyses, such as DESeq2 (Supplementary S11), MINT (Supplementary S2), Differential Heat Tree (Figure 2), the categories they up and down-regulated are represented with corresponding up and down arrows. For example, in HW30 vs O30 comparison, “(S), ↑O30 (D, H)” for *Phascolarctobacterium* should be read as selected in subset analysis: (S) and upregulated in O30 according to both DESeq2 and Differential Tree: ↑O30 (D, H).

**Figure 2:**
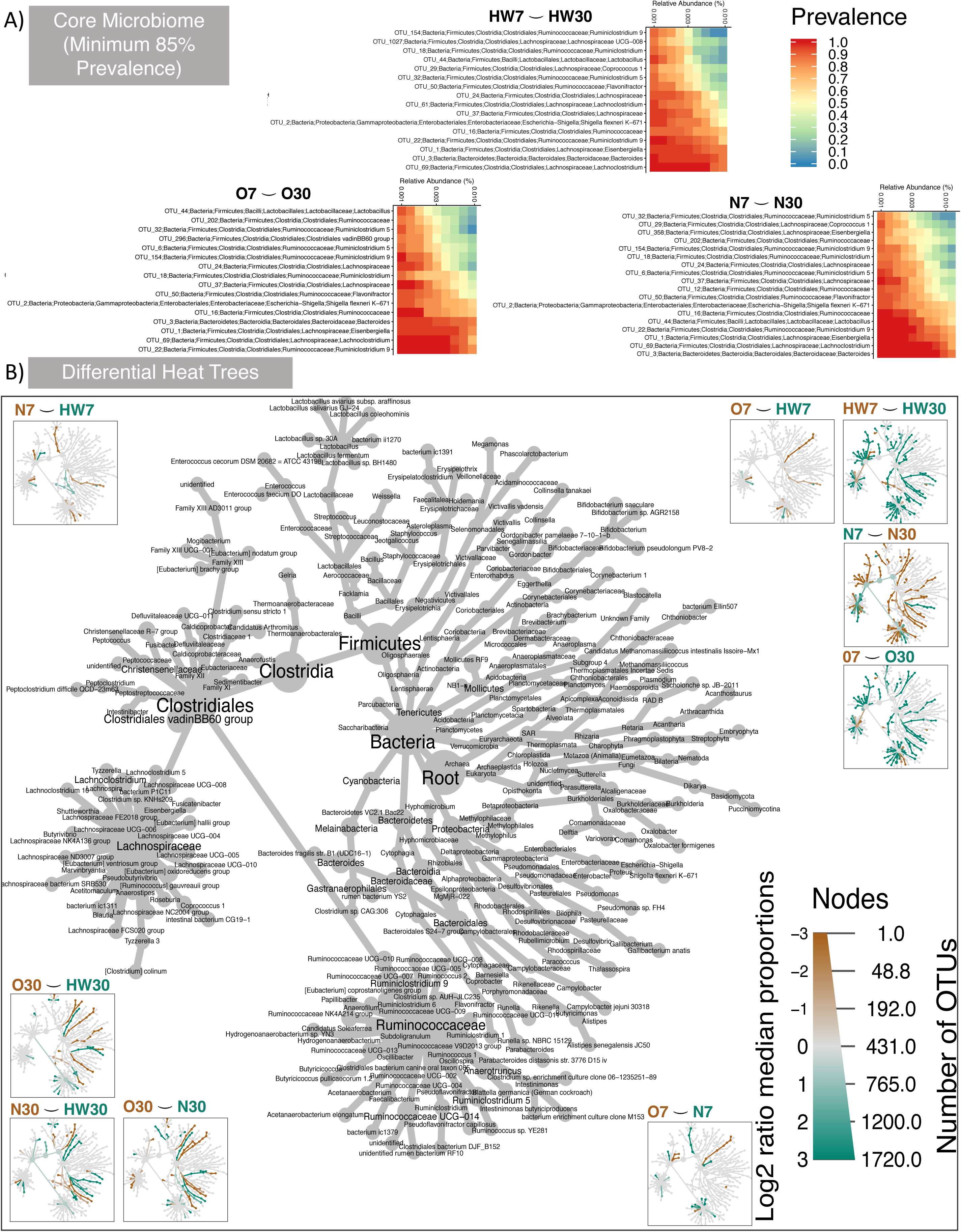
Taxa that persist and those that are differentially abundant. (a) Core microbiome analyses that persist in 85% of the samples for different production systems (day 7 and day 30). In the heat maps the OTUs are sorted by their abundances with those on the top being low abundant whereas those at the bottom are highly abundant. (b) Differential heat tree with taxonomy key given in the middle, and the branches where they are upregulated are colored according to their respective categories shown on top of each subpanel.

**Figure 3:**
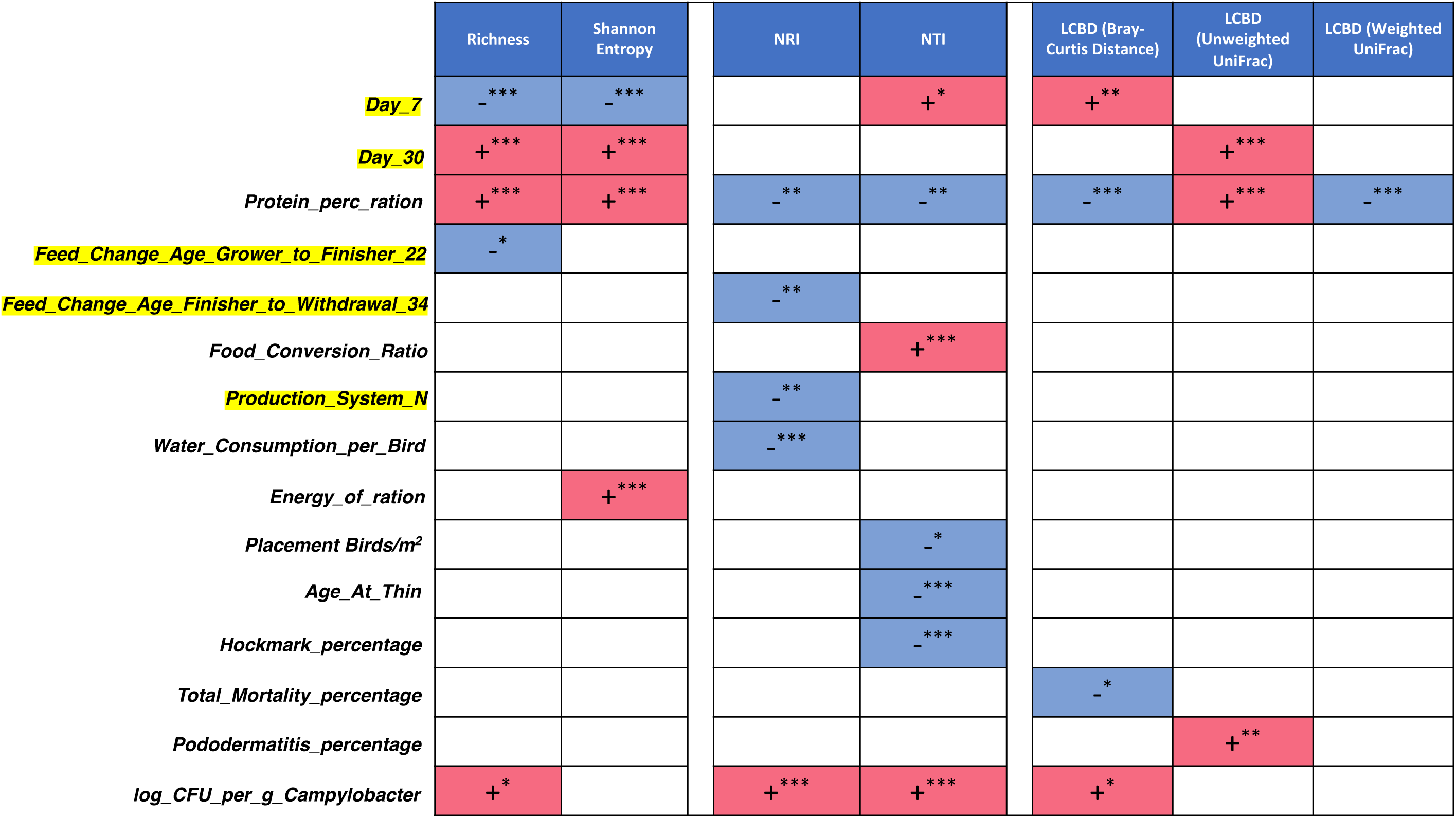
Heatmap of key extrinsic parameters that influence different attributes of microbiome. The figure is based on subset regressions (Supplementary S4 to S10), where red and blue represent the significant positive and negative beta coefficients that were consistently selected in different regression models. The categorical variables are represented with a yellow highlight (coded as 1 (present) or 0 (absent)) and if selected is interpreted as the samples belonging to those categories having positive/negative influence on the respective microbiome metrics.

### Campylobacter isolation and identification

The contents of a pair of ceca were transferred into a sterile stomacher bag (∼10 g ± 1 g), diluted with maximum recovery diluent (MRD) buffer to make a 1/10 dilution and stomached at 260 rpm for 1 minute. Further decimal dilutions were carried out in MRD to give a range of dilutions suitable to achieve countable plates. 100 µl of the original suspension (10^−1^ dilution) was inoculated onto duplicate plates of mCCDA (Oxoid, UK). The inoculum was spread uniformly over the surface of the agar plate with a sterile spreader until fully absorbed. This spread plating was repeated for all the other decimal dilutions. When low numbers of *Campylobacter* were expected, the limit of detection was increased by also spread plating 1 ml of the original suspension. The 1 ml inoculum was inoculated over three plates. The plates were incubated in a microaerophilic atmosphere (5% oxygen, 10% carbon dioxide and 85% nitrogen) at 41.5°C and examined after 44 ± 4 hours for typical colonies of *Campylobacter* spp. Plates containing less than 300 colonies were counted. Colonies were considered typical if they were greyish (often with a metallic sheen), flat and moist, with a tendency to spread. *Campylobacter* numbers were expressed as CFU/g cecal content. Genomic DNA (gDNA) was also extracted from chicken ceca for 16S metagenomics experiments.

### Poultry Growth and Performance Measurements

Performance parameters on per flock basis were recorded in line with typical industrial practices. *Weight_Gain_per_Day, Feed_Conversion_Ratio, Total_Mortality_percentage, Energy_of_Ration, Protein_perc_ration, Water_Consumption_per_Bird, Age_At_Thin, Age_at_Total_Depopulation, PMI_Rejects_percentage, Hockmark_percentage, Pododermatitis_percentage*, log_CFU_per_g_Campylobacter, EPEF were all captured. These variables were then correlated with the microbial community’s composition in various statistical analyses (Supplementary S13).

### DNA extraction, 16S rRNA amplification and sequencing

Cecal gDNA was extracted using QIAamp DNA Stool Mini Kit according to the manufacturer’s instructions and stored at −20°C. 16S metagenomic sequencing library construction was performed using Illumina guidelines (Illumina, U.S.A). The 16S ribosomal primers used were V3 (tcgtcggcagcgtcagatgtgtataagagacagcctacgggnggcwgcag) and V4 (gtctcgtgggctcggagatgtgtataagagacaggactachvgggtatctaatcc) [22, 23]. A second PCR step was performed to attach dual indices and Illumina sequencing adapters using the Nextera XT Index kit. Sequencing was performed on Illumina MiSeq at LSHTM using a v3 300 bp paired-end kit.

### Bioinformatics and Statistical Analysis

Details are described in the supplementary methods (Supplementary S12).

## Results

### Diversity patterns representative of the production systems

At alpha-diversity level, to investigate how diversity within the samples were influenced by the production system, both richness and Shannon entropy were calculated for day 7 and day 30 samples from production systems N, HW and O. In the host microbiome, time has a significant effect on microbial richness with all production systems significantly increasing from day 7 to day 30 (Fig. 1a). For day 7, production system N displayed the highest microbial richness when compared to HW and O. At day 30, no statistical difference in terms of richness was identified between the three production systems. Beta diversity using *Bray-Curtis* was measured, along with these OTUs collated together at genera level with only top-25 most abundant genera shown next to the PCoA diagram (Fig. 1b). At day 7, in terms of abundance counts, samples from production system O are far off from the clusters that contain samples from N and HW, respectively. On the other hand, at day 30, N seems to have a different community structure as compared to N and HW production systems, respectively. The breakdowns of taxa going from finer (OTUs), up coarser levels (family, class, phylum etc), are shown in Supplementary S3. Note, we are only considering the top-25 most abundant taxa identified at different taxonomic levels. Of interest, clear differences were observed using visual cues when comparing production system N at day 30 to HW and O, where genera *Bacteroides* and *Alistipes* were present in production system N at higher abundances.

Ecological drivers of microbial community were explored observing the clustering in the phylogenetic tree of the OTUs and utilising phylogenetic alpha diversity measures such as NRI/NTI. Positive NRI/NTI values indicate strong phylogenetic clustering (driven by environmental filtering), whereas reduced values represent phylogenetic overdispersion (environment has little or no role to play). Here, using NRI and NTI we can observe an increase in N, HW and O samples respectively from day 7 to day 30. Since chicken ceca are already a constrained environment to begin with (as opposed to real environmental datasets), the values <0^a^ (traditionally this implies stochasticity) may not be feasible to ascertain randomness/stochasticity/competitive exclusion principle, and hence values should be taken relatively with an increasing value implying increasing host environmental pressure (in the immediate case it is the environment within a chicken, whilst extrinsic environmental factors such as surroundings where chicken are confined and their diet may also have a role to play). Fig. 1a corroborates our findings from our previous longitudinal study [14].

### Key drivers of microbial community structure variation in terms of beta diversity

Next, we wanted to explore what drives the beta diversity amongst different categories (production system) and time. In this regard, we employed the “BVSTEP” routine [24] which was used to search for the highest correlation, in a Mantel test, by imploding the abundance table at genera level to the absolute minimal set of genera that preserve the beta diversity between samples in reduced feature space as compared to the complete feature space. This subset analysis was performed by considering all samples and then correlating the subset of genera for these samples that explain approximately the same beta diversity distances through permutation analysis (Supplementary S1). At the same time, after imploding to the subset of genera, we wanted to see if the resulting subset still has discriminatory power (in terms of grouping) through PERMANOVA analysis. In addition to PERMANOVA analysis, we employed several discriminatory algorithms such as i) DESeq2 with adjusted p-value significance cut-off of 0.05 and log2 fold change cut-off of 2 to find upregulated genera and vice versa (Supplementary S11), ii) Heat Tree analysis to find clades that were differentially expressed (Fig. 2), and iii) MINT analysis to consolidate the genera that have simultaneous discriminatory power at both spatial (production system N, HW and O) and temporal scales (day 7 and day 30)(Supplementary S2). To aid interpretation, all up/downregulated taxa are annotated with up and down arrows in Figure 1 according to which analysis they were selected in and with taxa from subset analysis as the seeding point. Redundancy in these analyses was to ensure there was no biases associated with the mathematical algorithms to find discriminating terms. In some cases, direction of the discrimination was contradictory, for example, there were a few cases where DESeq2 and Heat Tree agreed, whilst MINT analysis showed a reciprocal trend. This latter disagreement may possibly be because MINT consolidates all possible sources of variation. Nonetheless, we wanted to be comprehensive and therefore discuss only those taxa which achieve a majority consensus on multiple analyses albeit MINT underperforming in most cases.

For production system HW, *Alistipes, Ruminococcaceae UCG-014* and *Escherichia-Shigella* were significant genera increasing from day 7 to day 30. *Bacteroides* displayed significant increase at day 7 (decreasing at day 30) using DESeq2 and Heat Tree. For production system O, *Thalassospira, Alistipes* and *Bacteroides* increased at day 30 (*Bacteroides* directionality was only observed with MINT analysis). *Lachnoclostridium, Eisenbergiella, Escherichia-Shigella* all increased at day 7 (decreasing at day 30). For production system N, *Lachnoclostridium, Eisenbergiella* increased at day 7 (decreasing at day 30). *Bacteroides* was identified as decreasing at day 7 (increasing at day 30; directionality was only observed with MINT analysis). In general, *Alistipes* was observed to increase consistently at day 30 for all production systems, whilst all other genera showed mixed trends. Although it should be noted that *Alistipes* was present at higher abundance at production system N as compared to others.

Key genera were identified when comparing production systems at day 7. For HW vs O comparison, *Lachnoclostridium* was identified as increased for HW. *Escherichia-Shigella* was identified increased for production system O. For O vs N comparison, *Bacteroides* was increased for O. This was also replicated for HW when comparing to N. Key genera were identified when comparing production systems at day 30. For HW vs O comparison, *Phascolarctobacterium, Thalassospira* and *Bacteroides* were identified as increased for O, whereas *Barnesiella* was increased for HW when comparing to O. *Subdoligranulum, Eisenbergiella, Alistipes, Ruminiclostridium* 5 and *Ruminococcaceae UCG-014* were all identified as part of the subsets that explain beta diversity, although they were not implicated as discriminating in differential analyses. For N vs HW comparison at day 30, *Eisenbergiella* was increased for HW, and *Alistipes* was increased for N. For O vs N comparison at day 30, *Bacteroides* was observed with subset analysis alone. *Thalassospira* was observed, yet its discriminatory power was inconclusive.

Core microbiome where genera persist in 85% of the samples (something that is traditionally used to define the prevalence of core microbiome) for different production systems (day 7 and day 30) was assessed (Fig. 2). Genera identified include *Bacteroides, Lachnoclostridium, Eisenbergiella, Ruminoclostridium 9, Lactobacillus, Ruminococcaceae, Shigella flexneri K-671, Flavonifractor, Ruminococcaceae, Lachnospiraceae, Ruminiclostridium 5, Ruminiclostridium, Coprococcus 1, Ruminiclostridium 5* at varying level of abundance. In Fig. 2, OTUs are sorted by their abundances with those on the top being low abundant prevalent OTUs, whereas those at the bottom are highly abundant prevalent OTUs.

### Parameters deriving microbial community structure

Analysis of parameters that had a significant effect on microbial diversity were assessed using PERMANOVA against performance parameters when using different dissimilarity measures on microbiome data (Table 1). Using R^2^, if significant, to represent the variability explained in the community structure for Bray-Curtis distance, the parameter with the greatest impact was days, explaining 21.2% of the variation. Next, key parameters of interest were: *log_CFU_per_g_Campylobacter* (4.1% variability), *Energy_of_Ration* (4.0% variability) and *Protein_perc_ration* (1.2% variability). Using both unweighted and weighted *UniFrac* as a beta diversity measure, the pattern was more or less similar with the same trends with days having the greatest impact on microbial diversity (32.7% and 43.0% respectively). All other parameters had R^2^ values at 1-5%.

**Table 1:** Statistics for PERMANOVA against performance parameters when using different dissimilarity measures on microbiome data. R^2^ represents the proportion of variability explained, for example, using “Days” and “Bray-Curtis” dissimilarity, the days explain 21.2% variability in microbial community structure.

| Bray-Curtis Distance |  |  |  |  |  |  |
| --- | --- | --- | --- | --- | --- | --- |
| *Predictors* | *Df* | *SumsOfSqs* | *MeanSqs* | *F.Model* | *R<sup>2</sup>* | *p* |
| Days | 1 | 4.9513 | 4.9513 | 26.8712 | 0.21163 *** | <0.001 |
| No. of Parent Flocks Used | 1 | 1.065 | 1.065 | 5.7796 | 0.04552 *** | <0.001 |
| Birds Placed | 1 | 1.0209 | 1.0209 | 5.5407 | 0.04364 *** | <0.001 |
| log CFU/g Campylobacter | 1 | 0.9606 | 0.9606 | 5.2131 | 0.04106 *** | <0.001 |
| Protein (% of Ration) | 1 | 0.2729 | 0.2729 | 1.4813 | 0.01167 . | <0.1 |
| Energy of Ration (Kcal/lb AME) | 1 | 0.9375 | 0.9375 | 5.088 | 0.04007 *** | <0.001 |
| Residuals | 77 | 14.1881 | 0.1843 | 0.60642 |  |  |
| Total | 83 | 23.3964 | 1 |  |  |  |
| Unweighted UniFrac Distance |  |  |  |  |  |  |
| *Predictors* | *Df* | *SumsOfSqs* | *MeanSqs* | *F.Model* | *R<sup>2</sup>* | *p* |
| Days | 1 | 4.2369 | 4.2369 | 46.886 | 0.32727 *** | <0.001 |
| No. of Parent Flocks Used | 1 | 0.3242 | 0.3242 | 3.588 | 0.02504 ** | <0.01 |
| Birds Placed | 1 | 0.6806 | 0.6806 | 7.532 | 0.05258 *** | <0.001 |
| log CFU/g Campylobacter | 1 | 0.386 | 0.386 | 4.272 | 0.02982 *** | <0.001 |
| Protein (% of Ration) | 1 | 0.1251 | 0.1251 | 1.385 | 0.00966 | 0.1744 |
| Energy of Ration (Kcal/lb AME) | 1 | 0.235 | 0.235 | 2.601 | 0.01816 * | 0.0165 |
| Residuals | 77 | 6.9582 | 0.0904 | 0.53747 |  |  |
| Total | 83 | 12.9461 | 1 |  |  |  |
| Weighted UniFrac Distance |  |  |  |  |  |  |
| *Predictors* | *Df* | *SumsOfSqs* | *MeanSqs* | *F.Model* | *R<sup>2</sup>* | *p* |
| Days | 1 | 0.055783 | 0.055783 | 76.574 | 0.43079 *** | <0.001 |
| No. of Parent Flocks Used | 1 | 0.001894 | 0.001894 | 2.6 | 0.01463 . | 0.0509 |
| Birds Placed | 1 | 0.005532 | 0.005532 | 7.594 | 0.04272 *** | **<0.001** |
| log CFU/g Campylobacter | 1 | 0.00484 | 0.00484 | 6.644 | 0.03738 *** | **<0.001** |
| Protein (% of Ration) | 1 | 0.00092 | 0.00092 | 1.262 | 0.0071 | 0.2534 |
| Energy of Ration (Kcal/lb AME) | 1 | 0.004429 | 0.004429 | 6.08 | 0.0342 ** | **<0.01** |
| Residuals | 77 | 0.056093 | 0.000728 | 0.43318 |  |  |
| Total | 83 | 0.129491 | 1 |  |  |  |

. $p < 0.1$ \* $p < 0.05$ \*\* $p < 0.01$ \*\*\* $p < 0.001$

### Direction of influence for extrinsic parameters influencing key metrics for the microbiome

Whilst PERMANOVA analyses show the extent of influence on microbiome structure in terms of variability, to obtain directions as to whether an increase or decrease in these parameters cause an increase or decrease in the properties of microbiome, we resorted to performing subset regressions on one-dimensional realisation of microbiome (alpha, beta diversity measures etc). These subset regressions permuted through all possible subsets of explanatory variables (extrinsic parameters considered in this study) by ranking them in terms of quantitative fit after performing cross-validation (Supplementary S4 to S10 and summarised in Fig. 3). Note that red and blue backgrounds represent whether predictors have a positive or a negative influence along with the significances, respectively in the regression model. In addition, all categorical variables were “dummyfied” (a standard procedure) to represent as present/absent as a tag and were used in the regression model to see whether their inclusion/exclusion has an effect on the final model. As expected, measuring alpha diversity, inclusion of day 30 samples increases richness and Shannon entropy. Although inclusion of day 7 samples led to an increased environmental pressure, it was marginally significant and therefore deterministic nature of microbial communities at local and terminal clade level should be taken with caution. In terms of how samples differ from each other, LCBD was also considered in subset regression analysis with positive/red explanatory variables causing community structure to become different from the average community structure as a mean to identify groups that were markedly different. This was only observed for *Bray-Curtis* distance metric (that considers abundances of taxa without their phylogenetic distances) at day 7, and unweighted *UniFrac* (phylogenetic distance considering only presence/absence of taxa without considering their abundance) for day 30.

An increase in *Protein_perc_ration* led to a positive effect on microbial diversity, whilst simultaneously reducing the effect of environmental pressure on microbial community structure. In terms of beta diversity measure (LCBD with different distances), overall there was a reduction in beta diversity when considering *Protein_perc_ration*, although the trend was opposite for unweighted UniFrac. *Energy_of_ration* also causes the microbial diversity to be more even as it had a positive and significant influence on Shannon entropy. The *Feed_conversion_ratio* on the other hand only caused shift in environmental pressure affecting the terminal clades by making them more clustered though the NTI measure.

In terms of production systems, it was observed that samples belonging to production system N had less influence by the environment and were possibly driven by competitive exclusion principles. Of note, production system N also had the lowest (best) *Feed_conversion_ratio*. A decrease in *Feed_conversion_ratio* was noted to lead to reduced environmental influence, which aligns with production system N having a reduced environmental influence, and also that a higher *Placement_birds/m*^*2*^ resulting in a reduced environmental influence. The same phenomena were also observed when considering *Age_at_thin, Hockmark_ percentage* and *Water_consumption_per _bird* as explanatory variables. Interestingly, recorded *log_CFU_per_g_Campylobacter* led to an increase in microbial diversity, with an increase in environmental pressure (at global scale: NRI, and also at local terminal clustering: NTI) as well as causing a marked shift in terms of beta diversity. Of all production systems, *Campylobacter* was only identified in production system N at day 30 based on 16S rRNA abundance count (Supplementary S2), corroborated with independent log CFU/g of *Campylobacter* measure.

## Discussion

Our data clearly show that there is a difference in microbial community structure between production systems with varying influence by extrinsic parameters considered within this study. We observed diversity increase significantly for day 30 when compared to day 7. This is in line with previous reports whereby the gastrointestinal (GI) tract of poultry comes into contact with exogenous microorganisms immediately after hatch and as the host grows, this microbiome becomes highly diverse until it reaches a relatively stable yet dynamic state [25]. An important finding in this study is that amongst the different production systems, only N is where microbial community assemblage seems to be random (less environmental pressure). Stocking density is clearly a parameter which if varied, can alter microbial community structure significantly. This demonstrates that production systems can be modified to alter the microbiome profile influencing performance at farm level.

Dietary nutrient components are implicated in improving the performance of broiler chickens. An increasing *Energy_of_ration* and decreasing *Protein_perc_ration* in feed over time, pertain to all production systems within this study. These variables have a direct influence on gut microbial composition, and in conjunction with differentiating parameters between production systems (i.e. stocking density), lead to differences in microbial composition between production systems. The role of diet is sufficiently important for microbial community structure assemblage as previously described whereby digestion of non-starch polysaccharides (NSPs; found in the grain of chicken feed) lead to production of SCFA, which are absorbed across mucosa and catabolised by the host [15, 26]. SCFAs contribute to chicken nutrition and also lower pH which can inhibit acid-sensitive pathogens and improve mineral absorption [17, 18]. Thus, an association exists between the consistent and differentiating parmeters of production systems, that affect feed utilisation, leading to competitive exclusion of genera based on competition for nutrients and other factors.

Genera that were differentially expressed between different production systems and days were identified (some also part of the core microbiome). For day 30, HW vs O comparison, *Phascolarctobacterium, Thalassospira* and *Bacteroides* were identified as increased for O. *Phascolarctobacterium* is involved in SCFA production, including acetate and propionate and described as an option for reduction of *Campylobacter* via competitive exclusion [27-29]. Here, Omega-3 fed poultry systems harboured *Phascolarctobacterium* at a higher prevalence, although we cannot state if this was directly associated with a reduction or absence of *Campylobacters* from the cecal community. *Eisenbergiella* was increased for HW vs N comparison at day 30, and *Alistipes* was increased for N when compared to HW at day 30. A reduction in *Eisenbergiella* has been associated with gastrointestinal disorders linked to metabolic and microbiota changes (functional dyspepsia) resulting in defective energy metabolism, amino acids, nucleotides and SCFA [30]. The HW system may improve the metabolism-microbiome interaction and could result in a competitive exclusion of bacterial pathogens via a fortified immune system. A dysfunctional microbiota can induce metabolic, autoimmune and inflammatory diseases, and can seriously undermine gut function [31-33]. *Barnesiella* was increased for HW when comparing to O. Presence of *Barnesiella* has been associated with prevention and spread of highly antibiotic-resistant bacteria [34] where the gut bacterial community may react to antibiotic inclusion in the diet and prophylactically encourage the presence of *Barnesiella*, an effect demonstrated in mice where ampicillin treatment increased their presence [35]. The competitive exclusion hypothesis is supported by the increase of *Alistipes* bacteria in N group at day 30, and using the top-25 most abundant taxa identified at OTU level where clear differences for N against HW and O were observed (also for genera *Bacteroides*). The presence of *Subdoligranulum* bacteria in HW and O production systems, although not discriminatory, still contributing to beta diversity, represents a sign of improved gut health as these bacteria are known to be involved in production of SCFAs (e.g. butyrate) with an important role in gut physiology [36]. *Ruminiclostridium 5*, identified within HW and O production systems, also within core microbiome, has been noted to impact SCFA concentration within the gut [37].

The European Union (EU) ban on antimicrobial growth promoters in 2006 has created an increased need to devise alternative methods to improve performance and potentially reduce numbers of pathogenic bacteria. Examples include use of natural plant derived products such as carvacrol [38], addition of dietary prebiotics [39] and administration of live probiotic bacteria [40]. More recently, prebiotic galacto-oligosaccharides (GOS) have been added to broiler feed and enhanced the growth rate and feed conversion of chickens relative to those obtained with a calorie-matched control diet [41]. Recent developments has observed chicken diets enriched with Omega-3 PUFA’s for benefits to human health [5]. Omega-3 fatty acids α-linolenic acid (ALA) (often found in plants), eicosapentaenoic acid (EPA) and docosahexaenoic acid (DHA) (found typically in marine oils), all play a role in forming structural components of cell membranes, serve as precursors to bioactive lipid mediators, and provide a source of energy [42]. PUFAs are also important as substrates for inflammatory and anti-inflammatory acids, with EPA is believed to have anti-inflammatory properties [43]. One of the proposed mechanisms as to how dietary antibiotics exert their growth promoting benefits is via anti-inflammatory effects towards the intestinal epithelium, by inhibition of production and excretion of catabolic mediators [44]. Increased levels of EPA following antibiotic supplementation align with this non-antibiotic, anti-inflammatory theory of antibiotic growth promotion [45]. In our study, we do not know if PUFAs were fully absorbed, and to what effect the interaction is with host microbiome, however we do observe an increase in weight for production system O which has Omega-3. Of note, production system N had the lowest (best) FCR.

Remarkably, *log_CFU_per_g_Campylobacter* led to an increase in microbial diversity, an increase in environmental assemblage metrics NRI and NTI, and also increased divergence in community structure from other samples. *Campylobacter* was identified in production system N at day 30 (Supplementary S2) along with measuring the log CFU/g of *Campylobacter* corroborating the 16S data. It has been previously reported that *Campylobacter* typically appears within week two of the life cycle [11-14, 46]. The lack of identification of *Campylobacter* at any of the day 7 samples is anticipated. It is interesting that detection of *Campylobacter* was only observed at day 30 for production system N. This may have been due to limitation of sampling points. Although we recognise the diversity will increase naturally from day 7 to day 30, based on our data, *Campylobacter* is associated with an increasing diversity. Broiler genetics is known to impact microbial diversity [47, 48] and this may explain why in our study the environmental pressure was significantly impacted positively by the presence of *Campylobacter*. Here the environmental influence may be the chicken host genetics influencing microbial community structure.

## Conclusion

Our findings demonstrate a relative role of different production system parameters in shaping the bacterial communities’ impact on the chicken microbiome, with stocking density playing a major role influencing microbial dynamics. Specific genera with higher prevalence were identified within production systems that have key roles in energy metabolism, amino acid, nucleotide and SCFA utilisation. It is clear that parameters between production system (whether constant or variable) have an impact on microbial diversity which subsequently influences feed breakdown and hence instigates competitive exclusion of certain genera. Omega-3 had a positive impact on weight gain and *Campylobacter* presence was linked with environmental pressure, which may be the external environment or the host itself. Future studies that will direct the optimisation of extrinsic parameters and optimisation of diets targeting microbes with the underlying benefits of improving performance will aid in reducing pathogens such as *Campylobacter*. Our study is the first to investigate the relative importance of production system parameters in an industrial farm environment without any intervention strategy (studying cecal microbiome at its natural environment), to reveal the factors that link microbial community structure to improved broiler performance, and reduced pathogenic bacteria such as *Campylobacter*.

## Supporting information

Supplementary S1-Subset Analysis

Supplementary S2-MINT Analysis

Supplementary S3-Top 25 Abundant Taxa

Supplementary S4-Richness Subset Regression

Supplementary S5-Shannon Entropy Subset Regression

Supplementary S6-NRI Subset Regression

Supplementary S7-NTI Subset Regression

Supplementary S8-LCBD Bray-Curtis Subset Regression

Supplementary S9-LCBD Unweighted UniFrac Subset Regression

Supplementary S10-LCBD Weighted UniFrac Subset Regression

Supplementary S11-Pairwise Differential Analysis

Supplementary S12-Bioinformatics_and_StatisticalAnalysis

Supplementary S13-Parameters Measured

## Availability of supporting data

The raw sequence files supporting the results of this article are available in the European Nucleotide Archive under the project accession number PRJEB34742.

## Competing interests

AM, AR and UL are employed by company Moy Park. AM is also a PhD student between Moy Park and AFBI. All other authors declare no competing interests.

## Author contributions

AM, AR, UL, UZI, NC and OG contributed to the study design. NC and OG managed the study. CK, AM and ML performed the sample collection and DNA extraction. AM and OG performed the library preparation and Illumina MiSeq sequencing at the LSHTM. UZI wrote the analysis scripts to generate the figures and tables in this paper. UZI, AM and OG performed the bioinformatics and statistical analysis. UZI, AM, NC and OG drafted the initial version of the manuscript with all authors including ND, BWW and WTS contributing to redrafting.

## Funding

OG and NC acknowledge research funding from Moy Park. UZI is funded by NERC Independent Research Fellowship (NE/L011956/1).

## List of Supplementary Material

**Supplementary S1-Subset Analysis.docx**: Subset analysis from BVSTEP routine listing top subsets with highest correlation with the full genera table considering Bray-Curtis distance and considering samples provided in the first column. For each subset, PERMANOVA was performed against Groups considered in the first column.

**Supplementary S2-MINT Analysis.pdf**: **MINT study-wise discriminant analysis between treatments (N, HW, and O)**. The algorithm is a two-step process where **(a)** two components were found that reduce the classification error rates (using mahalanobis.dist in the function) in the algorithm, showing the ordination of samples using all the genera in the first two components (MINT PLS-DA) with ellipse representing 95% confidence interval and percentage variations explained by these components in axes labels. In step two, **(b)** the number of discriminating genera were found for each component, highlighted as diamonds, **(c)** is similar to **(a)** however the ordination was considered using the discriminants from two components (MINT sPLS-DA); **(d)** shows the heatmap of these discriminant genera, with both rows and columns ordered using hierarchical (average linkage) clustering to identify blocks of genera of interest. Heatmap depicts TSS+CLR normalised abundances: high abundance (pink) and low abundance (blue), along with metadata drawn on top. Table on the right side shows the summary statistics for *Campylobacter* reads obtained for each category.

**Supplementary S3-Top 25 Abundant Taxa.pdf**: Community structure based on relative abundance of the top-25 most abundant taxa identified at different taxonomic groups, where ‘others’ refers to all taxa not included in the ‘top-25’.

**Supplementary S4-Richness Subset Regression.docx**: Subset regression analysis using Richness as the dependent variable.

**Supplementary S5-Shannon Entropy Subset Regression.docx**: Subset regression analysis using Shannon Entropy as the dependent variable.

**Supplementary S6-NRI Subset Regression.docx**: Subset regression analysis using phylogenetic alpha diversity (NRI) as the dependent variable.

**Supplementary S7-NTI Subset Regression.docx:** Subset regression analysis using phylogenetic alpha diversity (NTI) as the dependent variable.

**Supplementary S8-LCBD Bray-Curtis Subset Regression.docx:** Subset regression analysis using beta diversity measure (LCBD with Bray-Curtis distance) as the dependent variable.

**Supplementary S9-LCBD Unweighted UniFrac Subset Regression.docx:** Subset regression analysis using beta diversity measure (LCBD with Unweighted UniFrac distance) as the dependent variable.

**Supplementary S10-LCBD Weighted UniFrac Subset Regression.docx:** Subset regression analysis using beta diversity measure (LCBD with Weighted UniFrac distance) as the dependent variable.

**Supplementary S11-Pairwise Differential Analysis.xlsx:** Differential analysis of genera that are up/down-regulated between different groups (Adjusted P values ≤ 0.05) with at least log2 fold change from the base mean abundances for the samples considered in the first column.

**Supplementary S12-Bioinformatics_and_StatisticalAnalysis.docx:** Bioinformatics and statistical analysis methods used within the study.

**Supplementary S13-Parameters Measured.docx:** A list of all of the parameters measured within the study and their explanations.

## Footnotes

a It should be noted that NRI reflects phylogenetic clustering in a broad sense (whole phylogenetic tree) with lower values representing evenly spread community. On the other hand, NTI focuses more on the tips of the tree with positive values of NTI indicating that species co-occur with more closely related species than expected, and lower values indicating that closely related species do not co-occur by chance.

## Notes

### Competing Interest Statement

AM, AR and UL are employed by company Moy Park. AM is enrolled on a PhD programme at QUB, and undertook research work at AFBI and Moy Park. All other authors declare no competing interests.

### Summary of Updates

Competing interest statement update and minor updates throughout the text.

## References

1. Borda-Molina D, Seifert J, Camarinha-Silva A: Current perspectives of the chicken gastrointestinal tract and its microbiome. Computational and structural biotechnology journal 2018, 16:131–139.

2. FAO: Food outlook: biannual report on global food markets - November 2018. pp. 104: Rome; 2018:104.

3. Sood U, Gupta V, Kumar R, Lal S, Fawcett D, Rattan S, Poinern GEJ, Lal R: Chicken gut microbiome and human health: past scenarios, current perspectives, and futuristic applications. Indian Journal of Microbiology 2019:1–10.

4. Stanley D, Hughes RJ, Moore RJ: Microbiota of the chicken gastrointestinal tract: influence on health, productivity and disease. Applied microbiology and biotechnology 2014, 98:4301–4310.

5. Stanton AV, Shortall K, El-Sayed T, O’Donovan F, James K, Kennedy J, Hayes H, Fahey A, Dolan E, Williams D: Eating Omega-3 Polyunsaturated Fatty Acid Enriched Chicken-Meat and Eggs Results in Increased Plasma Docosahexaenoic and Eicosapentaenoic Acid Levels and an Improved Omega-3-Index. Circulation 2017, 136:A19913–A19913.

6. Amour C, Gratz J, Mduma E, Svensen E, Rogawski ET, McGrath M, Seidman JC, McCormick BJ, Shrestha S, Samie A, et al: Epidemiology and Impact of Campylobacter Infection in Children in 8 Low-Resource Settings: Results From the MAL-ED Study. Clin Infect Dis 2016, 63:1171–1179.

7. Hermans D, Pasmans F, Heyndrickx M, Van Immerseel F, Martel A, Van Deun K, Haesebrouck F: A tolerogenic mucosal immune response leads to persistent Campylobacter jejuni colonization in the chicken gut. Crit Rev Microbiol 2012, 38:17–29.

8. Humphrey S, Chaloner G, Kemmett K, Davidson N, Williams N, Kipar A, Humphrey T, Wigley P: Campylobacter jejuni is not merely a commensal in commercial broiler chickens and affects bird welfare. MBio 2014, 5:e01364–01314.

9. Van Deun K, Pasmans F, Ducatelle R, Flahou B, Vissenberg K, Martel A, Van den Broeck W, Van Immerseel F, Haesebrouck F: Colonization strategy of Campylobacter jejuni results in persistent infection of the chicken gut. Vet Microbiol 2008, 130:285–297.

10. Wigley P: Blurred Lines: Pathogens, Commensals, and the Healthy Gut. Front Vet Sci 2015, 2:40.

11. Kalupahana RS, Kottawatta KS, Kanankege KS, van Bergen MA, Abeynayake P, Wagenaar JA: Colonization of Campylobacter spp. in broiler chickens and laying hens reared in tropical climates with low-biosecurity housing. Appl Environ Microbiol 2013, 79:393–395.

12. Neill SD, Campbell JN, Greene JA: Campylobacter species in broiler chickens. Avian Pathology 1984, 13:777–785.

13. Thibodeau A, Fravalo P, Yergeau E, Arsenault J, Lahaye L, Letellier A: Chicken Caecal Microbiome Modifications Induced by Campylobacter jejuni Colonization and by a Non-Antibiotic Feed Additive. PLoS One 2015, 10:e0131978.

14. Ijaz UZ, Sivaloganathan L, McKenna A, Richmond A, Kelly C, Linton M, Stratakos AC, Lavery U, Elmi A, Wren BW, et al: Comprehensive Longitudinal Microbiome Analysis of the Chicken Cecum Reveals a Shift From Competitive to Environmental Drivers and a Window of Opportunity for Campylobacter. Front Microbiol 2018, 9:2452.

15. Jozefiak D, Rutkowski A, Martin S: Carbohydrate fermentation in the avian ceca: a review. Animal Feed Science and Technology 2004, 113:1–15.

16. McNab JM: The Avian Caeca: A Review. World’s Poultry Science Journal 2007, 29:251–263.

17. Sergeant MJ, Constantinidou C, Cogan TA, Bedford MR, Penn CW, Pallen MJ: Extensive microbial and functional diversity within the chicken cecal microbiome. PLoS One 2014, 9:e91941.

18. Apajalahti J: Comparative Gut Microflora, Metabolic Challenges, and Potential Opportunities. The Journal of Applied Poultry Research 2005, 14:444–453.

19. Connerton PL, Richards PJ, Lafontaine GM, O’Kane PM, Ghaffar N, Cummings NJ, Smith DL, Fish NM, Connerton IF: The effect of the timing of exposure to Campylobacter jejuni on the gut microbiome and inflammatory responses of broiler chickens. Microbiome 2018, 6:88.

20. Huyghebaert G, Ducatelle R, Van Immerseel F: An update on alternatives to antimicrobial growth promoters for broilers. The Veterinary Journal 2011, 187:182–188.

21. AL-Darkazali H, Meevootisom V, Isarangkul D, Wiyakrutta S: Gene expression and molecular characterization of a xylanase from chicken cecum metagenome. International journal of microbiology 2017, 2017.

22. D’Amore R, Ijaz UZ, Schirmer M, Kenny JG, Gregory R, Darby AC, Shakya M, Podar M, Quince C, Hall N: A comprehensive benchmarking study of protocols and sequencing platforms for 16S rRNA community profiling. BMC Genomics 2016, 17:55.

23. Klindworth A, Pruesse E, Schweer T, Peplies J, Quast C, Horn M, Glockner FO: Evaluation of general 16S ribosomal RNA gene PCR primers for classical and next-generation sequencing-based diversity studies. Nucleic Acids Res 2013, 41:e1.

24. Clarke KR, Ainsworth M: A Method of Linking Multivariate Community Structure to Environmental Variables. Marine Ecology Progress Series 1993, 92:205–219.

25. Pan D, Yu Z: Intestinal microbiome of poultry and its interaction with host and diet. Gut Microbes 2014, 5:108–119.

26. McWhorter TJ, Caviedes-Vidal E, Karasov WH: The integration of digestion and osmoregulation in the avian gut. Biol Rev Camb Philos Soc 2009, 84:533–565.

27. Peralta-Sánchez JM, Martín-Platero AM, Ariza-Romero JJ, Zurita-González MJ, Baños A, Rodriguez-Ruano SM, Maqueda M, Valdivia E, Martínez-Bueno M, Rabelo-Ruiz M: Egg production in poultry farming is improved by probiotic bacteria. Frontiers in microbiology 2019, 10:1042.

28. Wu F, Guo X, Zhang J, Zhang M, Ou Z, Peng Y: Phascolarctobacteriumáfaecium abundant colonization in human gastrointestinal tract. Experimental and therapeutic medicine 2017, 14:3122–3126.

29. Zheng M, Mao P, Tian X, Meng L: Growth performance, carcass characteristics, meat and egg quality, and intestinal microbiota in Beijing-you chicken on diets with inclusion of fresh chicory forage. Italian Journal of Animal Science 2019, 18:1310–1320.

30. Luo L, Hu M, Li Y, Chen Y, Zhang S, Chen J, Wang Y, Lu B, Xie Z, Liao Q: Association between metabolic profile and microbiomic changes in rats with functional dyspepsia. RSC advances 2018, 8:20166–20181.

31. Ferreira CM, Vieira AT, Vinolo MAR, Oliveira FA, Curi R, Martins FdS: The central role of the gut microbiota in chronic inflammatory diseases. Journal of immunology research 2014, 2014.

32. Joyce SA, Gahan CG: The gut microbiota and the metabolic health of the host. Current opinion in gastroenterology 2014, 30:120–127.

33. Silva MJB, Carneiro MBH, dos Anjos Pultz B, Pereira Silva D, Lopes MEdM, dos Santos LM: The multifaceted role of commensal microbiota in homeostasis and gastrointestinal diseases. Journal of immunology research 2015, 2015.

34. Ubeda C, Bucci V, Caballero S, Djukovic A, Toussaint NC, Equinda M, Lipuma L, Ling L, Gobourne A, No D, et al: Intestinal microbiota containing Barnesiella species cures vancomycin-resistant Enterococcus faecium colonization. Infect Immun 2013, 81:965–973.

35. O’Loughlin JL, Samuelson DR, Braundmeier-Fleming AG, White BA, Haldorson GJ, Stone JB, Lessmann JJ, Eucker TP, Konkel ME: The Intestinal Microbiota Influences Campylobacter jejuni Colonization and Extraintestinal Dissemination in Mice. Appl Environ Microbiol 2015, 81:4642–4650.

36. Fleming SE, Fitch MD, DeVries S, Liu ML, Kight C: Nutrient utilization by cells isolated from rat jejunum, cecum and colon. J Nutr 1991, 121:869–878.

37. Song Y, Malmuthuge N, Steele MA, Guan LL: Shift of hindgut microbiota and microbial short chain fatty acids profiles in dairy calves from birth to pre-weaning. FEMS microbiology ecology 2018, 94:fix179.

38. Kelly C, Gundogdu O, Pircalabioru G, Cean A, Scates P, Linton M, Pinkerton L, Magowan E, Stef L, Simiz E, et al: The In Vitro and In Vivo Effect of Carvacrol in Preventing Campylobacter Infection, Colonization and in Improving Productivity of Chicken Broilers. Foodborne Pathog Dis 2017, 14:341–349.

39. Sethiya NK: Review on natural growth promoters available for improving gut health of poultry: an alternative to antibiotic growth promoters. Asian Journal of Poultry Science 2016, 10:1–29.

40. Gadde U, Kim W, Oh S, Lillehoj HS: Alternatives to antibiotics for maximizing growth performance and feed efficiency in poultry: a review. Animal health research reviews 2017, 18:26–45.

41. Richards PJ, Lafontaine GMF, Connerton PL, Liang L, Asiani K, Fish NM, Connerton IF: Galacto-oligosaccharides modulate the juvenile gut microbiome and innate immunity to improve broiler chicken performance. mSystems 2020, 5.

42. Jump DB, Tripathy S, Depner CM: fatty acid–regulated transcription factors in the liver. Annual review of nutrition 2013, 33:249–269.

43. Calder PC: Omega-3 fatty acids and inflammatory processes. Nutrients 2010, 2:355–374.

44. Niewold TA: The nonantibiotic anti-inflammatory effect of antimicrobial growth promoters, the real mode of action? A hypothesis. Poultry science 2007, 86:605–609.

45. Gadde UD, Oh S, Lillehoj HS, Lillehoj EP: Antibiotic growth promoters virginiamycin and bacitracin methylene disalicylate alter the chicken intestinal metabolome. Sci Rep 2018, 8:3592.

46. Hermans D, Van Deun K, Martel A, Van Immerseel F, Messens W, Heyndrickx M, Haesebrouck F, Pasmans F: Colonization factors of Campylobacter jejuni in the chicken gut. Vet Res 2011, 42:82.

47. Psifidi A, Fife M, Howell J, Matika O, Van Diemen PM, Kuo R, Smith J, Hocking PM, Salmon N, Jones MA: The genomic architecture of resistance to Campylobacter jejuni intestinal colonisation in chickens. BMC genomics 2016, 17:293.

48. Psifidi A, Kranis A, Rothwell L, Bremmer A, Russell K, Robledo D, Bush S, Fife M, Hocking P, Banos G: Genetic control of Campylobacter colonisation in broiler chickens: genomic and transcriptomic characterisation. bioRxiv 2020.

