## Supplementary S1-Subset Analysis for "Impact of industrial production system parameters on chicken microbiomes: mechanisms to improve performance and reduce *Campylobacter*"

| **Group Comparison** | **Subset No** | **Subset** | **Correlation of Subset with Full Table (R)** | **PERMANOVA Subsets (Groups)** |
| --- | --- | --- | --- | --- |
| HW 7, HW 30 | S1 | Alistipes + Ruminococcaceae UCG-014 + __Unknowns__ + Bacteroides + Escherichia-Shigella | 0.95186 | R^2^ = 0.39873 ( p = 0.001 *** ) |
|  | S2 | Alistipes + Ruminococcaceae UCG-014 + __Unknowns__ + Bacteroides | 0.94427 | R^2^ = 0.37498 ( p = 0.001 *** ) |
|  | S3 | Alistipes + __Unknowns__ + Bacteroides | 0.93051 | R^2^ = 0.37166 ( p = 0.001 *** ) |
|  | S4 | __Unknowns__ + Bacteroides | 0.88781 | R^2^ = 0.29359 ( p = 0.001 *** ) |
| N 7, N 30 | S1 | Lachnoclostridium + Eisenbergiella + Alistipes + __Unknowns__ + Bacteroides | 0.95873 | R^2^ = 0.23063 ( p = 0.001 *** ) |
|  | S2 | Eisenbergiella + Alistipes + __Unknowns__ + Bacteroides | 0.93960 | R^2^ = 0.21998 ( p = 0.001 *** ) |
|  | S3 | Lachnoclostridium + Alistipes + __Unknowns__ + Bacteroides | 0.92765 | R^2^ = 0.19553 ( p = 0.001 *** ) |
|  | S4 | Lachnoclostridium + Alistipes + __Unknowns__ | 0.91398 | R^2^ = 0.24042 ( p = 0.001 *** ) |
|  | S5 | Alistipes + __Unknowns__ | 0.89150 | R^2^ = 0.22381 ( p = 0.001 *** ) |
| O 7, O 30 | S1 | Thalassospira + Lachnoclostridium + Eisenbergiella + Alistipes + __Unknowns__ + Bacteroides + Escherichia-Shigella | 0.95778 | R^2^ = 0.22552 ( p = 0.001 *** ) |
|  | S2 | Thalassospira + Lachnoclostridium + Eisenbergiella + __Unknowns__ + Bacteroides + Escherichia-Shigella | 0.94049 | R^2^ = 0.19443 ( p = 0.001 *** ) |
|  | S3 | Thalassospira + Eisenbergiella + __Unknowns__ + Bacteroides + Escherichia-Shigella | 0.91998 | R^2^ = 0.19298 ( p = 0.001 *** ) |
|  | S4 | Thalassospira + Eisenbergiella + __Unknowns__ + Bacteroides | 0.88900 | R^2^ = 0.19891 ( p = 0.001 *** ) |
|  | S5 | Eisenbergiella + __Unknowns__ + Bacteroides | 0.85410 | R^2^ = 0.15187 ( p = 0.006 ** ) |
|  | S6 | __Unknowns__ + Bacteroides | 0.75341 | R^2^ = 0.08786 ( p = 0.079 . ) |
| HW 7, O 7 | S1 | Lachnoclostridium + Eisenbergiella + __Unknowns__ + Bacteroides + Escherichia-Shigella | 0.96934 | R^2^ = 0.23547 ( p = 0.001 ***) |
|  | S2 | Eisenbergiella + __Unknowns__ + Bacteroides + Escherichia-Shigella | 0.94559 | R^2^ = 0.25088 ( p = 0.001 ***) |
|  | S3 | Eisenbergiella + __Unknowns__ + Bacteroides | 0.92002 | R^2^ = 0.23325 ( p = 0.001 ***) |
|  | S4 | __Unknowns__ + Bacteroides | 0.86272 | R^2^ = 0.21508 ( p = 0.001 ***) |
| HW 30, O 30 | S1 | Phascolarctobacterium + Thalassospira + Barnesiella + Subdoligranulum + Eisenbergiella + Alistipes + Ruminiclostridium 5 + Ruminococcaceae UCG-014 + __Unknowns__ + Bacteroides | 0.95426 | R^2^ = 0.19066 ( p = 0.001 ***) |
|  | S2 | Phascolarctobacterium + Thalassospira + Barnesiella + Subdoligranulum + Eisenbergiella + Ruminiclostridium 5 + Ruminococcaceae UCG-014 + __Unknowns__ + Bacteroides | 0.94626 | R^2^ = 0.19812 ( p = 0.001 ***) |
|  | S3 | Phascolarctobacterium + Thalassospira + Barnesiella + Subdoligranulum + Eisenbergiella + Ruminococcaceae UCG-014 + __Unknowns__ + Bacteroides | 0.19392 | R^2^ = 0.19392 ( p = 0.001 ***) |
|  | S4 | Thalassospira + Barnesiella + Subdoligranulum + Eisenbergiella + Ruminococcaceae UCG-014 + __Unknowns__ + Bacteroides | 0.92366 | R^2^ = 0.17354 ( p = 0.001 ***) |
|  | S5 | Thalassospira + Barnesiella + Eisenbergiella + Ruminococcaceae UCG-014 + __Unknowns__ + Bacteroides | 0.90784 | R^2^ = 0.17534 ( p = 0.002 **) |
|  | S6 | Barnesiella + Eisenbergiella + Ruminococcaceae UCG-014 + __Unknowns__ + Bacteroides | 0.88736 | R^2^ = 0.129 ( p = 0.01 **) |
|  | S7 | Barnesiella + Eisenbergiella + __Unknowns__ + Bacteroides | 0.86871 | R^2^ = 0.13841 (p = 0.008 **) |
|  | S8 | Barnesiella + __Unknowns__ + Bacteroides | 0.84291 | R^2^ = 0.15315 ( p = 0.001 ***) |
|  | S9 | __Unknowns_+ Bacteroides | 0.78765 | *n.s.* |
| HW 7, N7 | S1 | Lachnoclostridium + Eisenbergiella + __Unknowns__ + Bacteroides | 0.95571 | R^2^ = 0.08014 ( p = 0.023 *) |
|  | S2 | Eisenbergiella + __Unknowns__ + Bacteroides | 0.93717 | R^2^ = 0.08092 ( p = 0.026 *) |
|  | S3 | __Unknowns__ + Bacteroides | 0.90413 | R^2^ = 0.1049 ( p = 0.012 *) |
| HW 30, N 30 | S1 | Eisenbergiella + Alistipes + __Unknowns__ | 0.95583 | R^2^ = 0.28872 ( p = 0.005 **) |
|  | S2 | Alistipes + __Unknowns__ | 0.94816 | R^2^ = 0.28872 ( p = 0.005 **) |
| O 7, N 7 | S1 | Eisenbergiella + __Unknowns__ + Bacteroides + Escherichia-Shigella | 0.95594 | R^2^ = 0.25699 ( p = 0.001 ***) |
|  | S2 | Eisenbergiella + __Unknowns__ + Bacteroides | 0.92974 | R^2^ = 0.2726 ( p = 0.001 ***) |
|  | S3 | __Unknowns__ + Bacteroides | 0.87716 | R^2^ = 0.28225 ( p = 0.001 ***) |
| O 30, N 30 | S1 | Thalassospira + __Unknowns__ + Bacteroides | 0.95880 | R^2^ = 0.4124( p = 0.001 ***) |
|  | S2 | Thalassospira + __Unknowns__ | 0.93878 | R^2^ = 0.50904 ( p = 0.001 ***) |
| N 7, HW 7, O 7, N 30, HW 30, O 30 | S1 | Lachnoclostridium + Eisenbergiella + Alistipes + Ruminococcaceae UCG-014 + __Unknowns__ + Bacteroides + Escherichia-Shigella | 0.95782 | R^2^ = 0.37638 ( p = 0.001 ***) |
|  | S2 | Eisenbergiella + Alistipes + Ruminococcaceae UCG-014 + __Unknowns__ + Bacteroides | 0.93456 | R^2^ = 0.3761 ( p = 0.001 ***) |
|  | S3 | Eisenbergiella + Alistipes + __Unknowns__ + Bacteroides | 0.92356 | R^2^ = 0.36959 ( p = 0.001 ***) |
|  | S4 | Alistipes + __Unknowns__ + Bacteroides | 0.88901 | R^2^ = 0.36892 ( p = 0.001 ***) |
|  | S5 | __Unknowns__ + Bacteroides | 0.84599 | R^2^ = 0.31154 ( p = 0.001 ***) |
| HW 7, O 7, N 7 | S1 | Eisenbergiella + __Unknowns__ + Bacteroides + Escherichia-Shigella | 0.95067 | R^2^ = 0.26252 ( p = 0.001 ***) |
|  | S2 | Eisenbergiella + __Unknowns__ + Bacteroides | 0.93018 | R^2^ = 0.25709 ( p = 0.001 ***) |
|  | S3 | __Unknowns__ + Bacteroides | 0.88047 | R^2^ = 0.2628 ( p = 0.001 ***) |
| HW 30, O 30, N 30 | S1 | Barnesiella + Alistipes + __Unknowns__ + Bacteroides | 0.95180 | R^2^ = 0.33785 ( p = 0.001 ***) |
|  | S2 | Barnesiella + __Unknowns__ + Bacteroides | 0.93998 | R^2^ = 0.3732 ( p = 0.001 ***) |
|  | S3 | __Unknowns__ + Bacteroides | 0.92161 | R^2^ = 0.32944 ( p = 0.001 ***) |
