## Supplementary figures and images for "Impact of industrial production system parameters on chicken microbiomes: mechanisms to improve performance and reduce *Campylobacter*"

### Supplementary S2-MINT Analysis

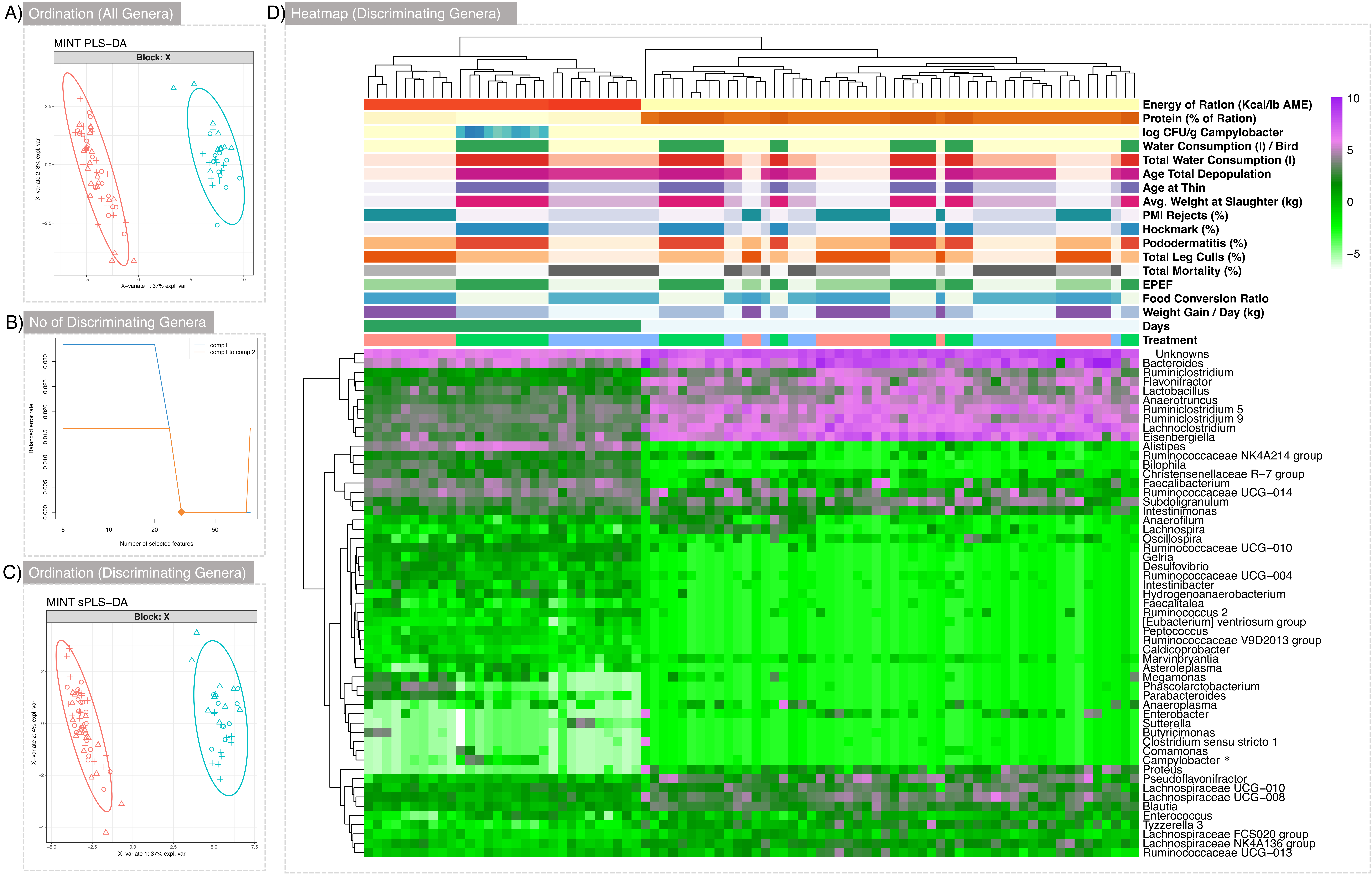


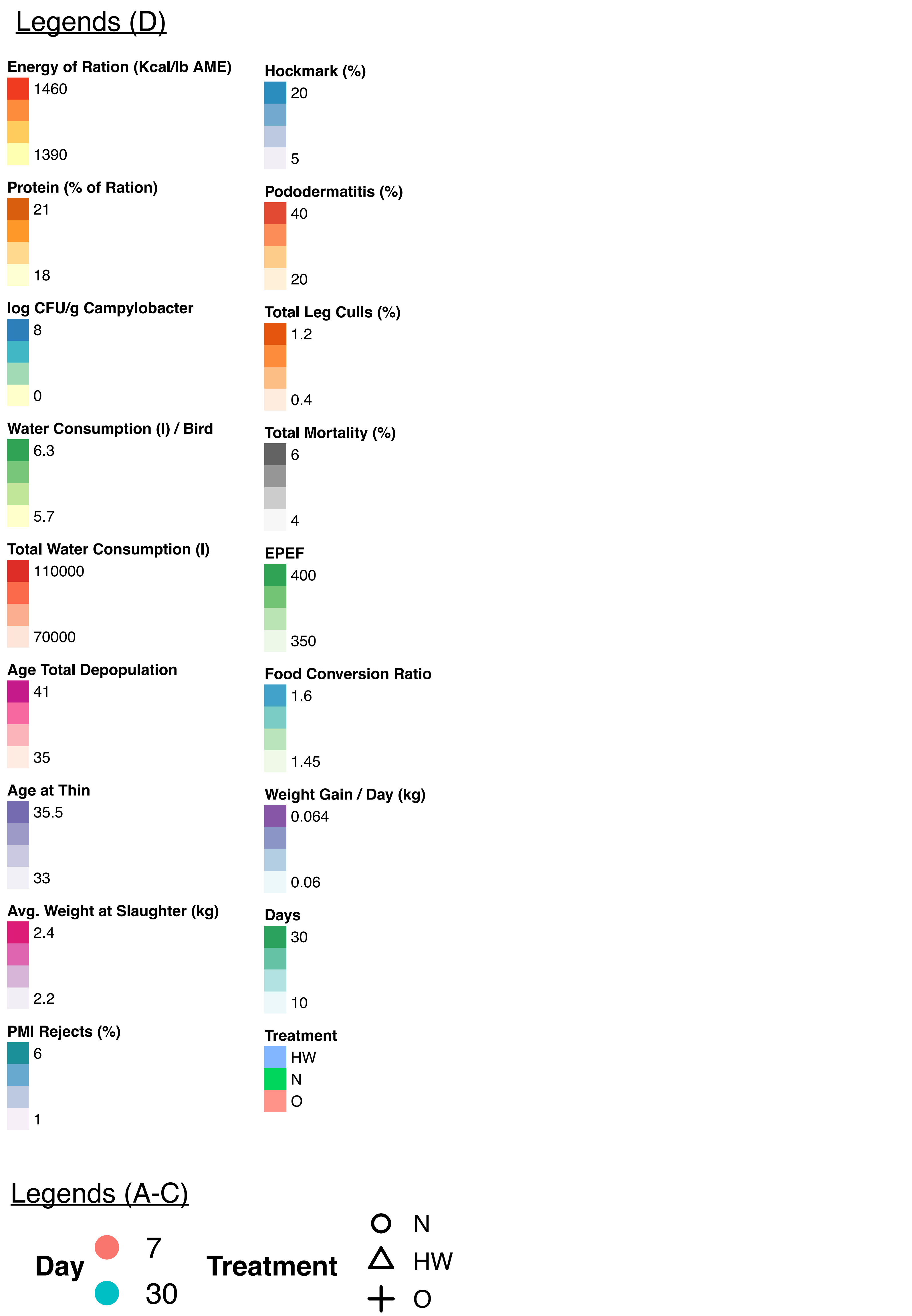


**Campylobacter* Reads in 16S rRNA Samples

| | Min | 1st. Quar. | Median | Mean | 3rd. Quar. | Max |
|---|---|---|---|---|---|---|
| HW 7 | 0 | 0 | 0 | 0.28 | 0 | 2 |
| N 7 | 0 | 0 | 0 | 0 | 0 | 0 |
| O 7 | 0 | 0 | 0 | 0 | 0 | 0 |
| HW 30 | 0 | 0 | 0 | 0 | 0 | 0 |
| N 30 | 0 | 13 | 35 | 472.50 | 100.5 | 2426 |
| O 30 | 0 | 0 | 0 | 0.5 | 0.75 | 3.0 |

### Supplementary S3-Top 25 Abundant Taxa

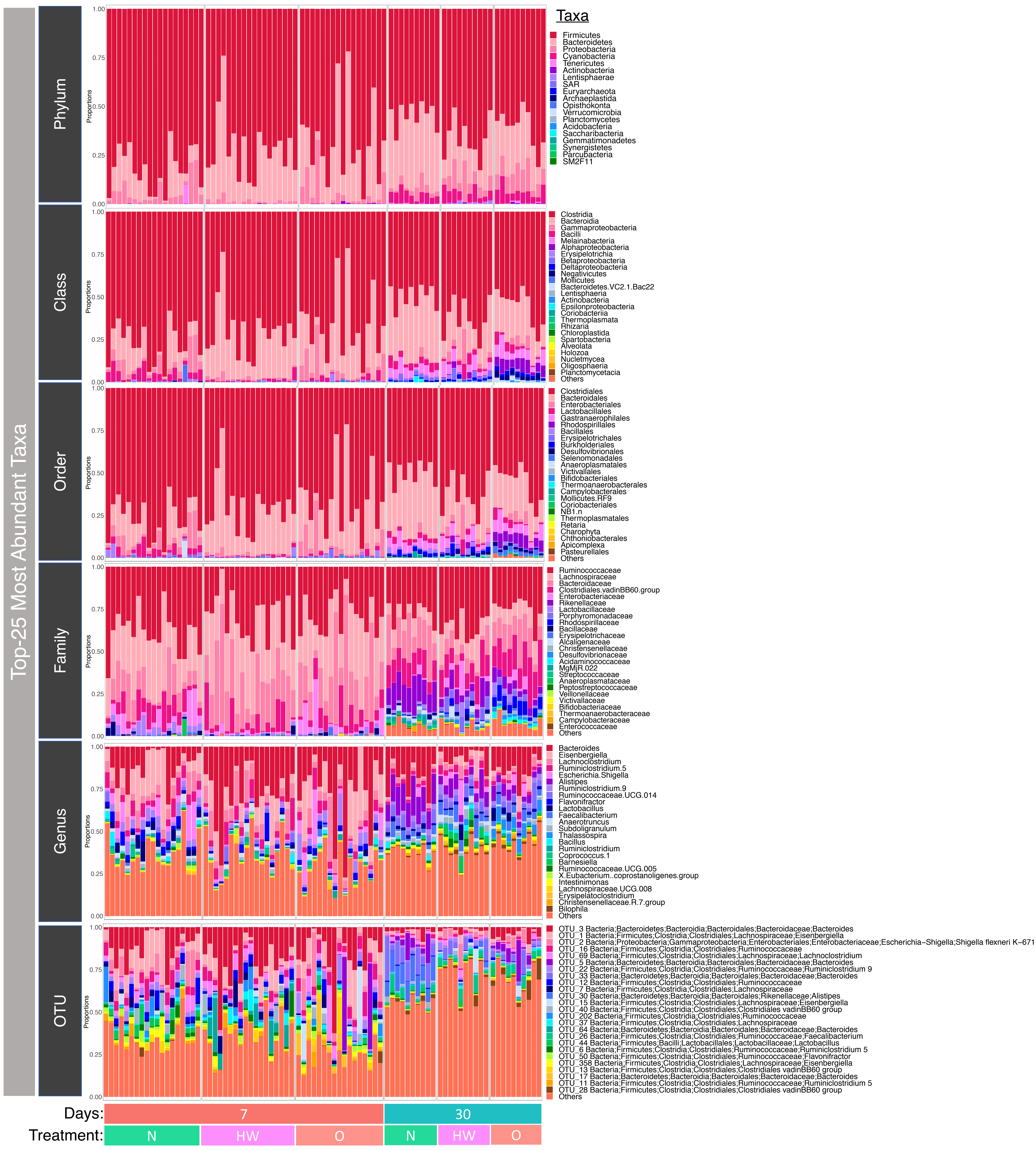
