## Supplementary S4-Richness Subset Regression for "Impact of industrial production system parameters on chicken microbiomes: mechanisms to improve performance and reduce *Campylobacter*"

**Top models (Model parameters given afterwards with significant positive influencers highlighted in orange and negative in blue):**

|  | Model | Cross-validation Errors (5-Folds Average RMSE) |
| --- | --- | --- |
| M2 | Richness ~ Day_7 + Protein_perc_ration | 60.37668 |
| M4 | Richness ~ Day_30 + Feed_Change_Age_Grower_to_Finisher_22 + log_CFU_per_g_Campylobacter + Protein_perc_ration | 61.31269 |
| M3 | Richness ~ Day_7 + Protein_perc_ration + Energy_of_Ration | 63.18702 |
| M5 | Richness ~ Feed_N + Day_30 + log_CFU_per_g_Campylobacter + Protein_perc_ration + Energy_of_Ration | 63.37658 |
| M6 | Richness ~ Feed_N + Day_7 + Total_Water_Consumption + log_CFU_per_g_Campylobacter + Protein_perc_ration + Energy_of_Ration | 63.79998 |
| M1 | Richness ~ Day_30 | 64.33040 |

|  | **Richness - M2** | | | | | | |
| --- | --- | --- | --- | --- | --- | --- | --- |
| *Predictors* | *Estimates* | *std. Error* | *std. Beta* | *CI* | *standardized CI* | *Statistic* | *p* |
| (Intercept) | -2003.17320 ^**^ | 675.00288 |  | -3346.21720 – -660.12921 |  | -2.96765 | **0.004** |
| Day_7 | -787.86843 ^***^ | 129.89801 | -2.41431 | -1046.32473 – -529.41213 | -3.19448 – -1.63414 | -6.06528 | **<0.001** |
| Protein_perc_ration | 140.68107 ^***^ | 37.15136 | 1.50731 | 66.76151 – 214.60063 | 0.72714 – 2.28748 | 3.78670 | **<0.001** |
| Observations | 84 | | | | | | |
| R^2^ / R^2^ adjusted | 0.862 / 0.859 | | | | | | |
| ** p<0.05   ** p<0.01   *** p<0.001* | | | | | | | |

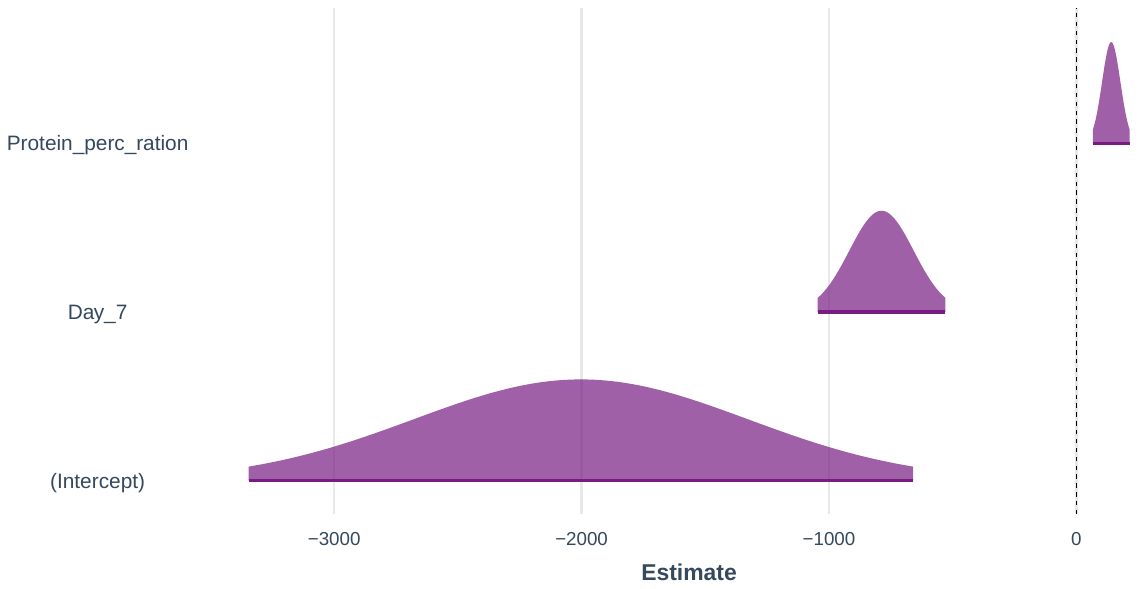

|  | **Richness – M4** | | | | | | |
| --- | --- | --- | --- | --- | --- | --- | --- |
| *Predictors* | *Estimates* | *std. Error* | *std. Beta* | *CI* | *standardized CI* | *Statistic* | *p* |
| (Intercept) | -7999.78990 ^**^ | 2370.77969 |  | -12718.70883 – -3280.87097 |  | -3.37433 | **0.001** |
| Day_30 | 1571.40876 ^***^ | 361.48778 | 4.81536 | 851.88532 – 2290.93219 | 2.64425 – 6.98647 | 4.34706 | **<0.001** |
| Feed_Change_Age_Grower_to_Finisher_22 | -97.33892 ^*^ | 45.12128 | -0.29597 | -187.15059 – -7.52726 | -0.56487 – -0.02707 | -2.15727 | **0.034** |
| log_CFU_per_g_Campylobacter | 27.70502 ^*^ | 11.20065 | 0.36600 | 5.41069 – 49.99936 | 0.07599 – 0.65601 | 2.47352 | **0.016** |
| Protein_perc_ration | 382.91607 ^***^ | 110.21832 | 4.10271 | 163.53199 – 602.30015 | 1.78815 – 6.41728 | 3.47416 | **0.001** |
| Observations | 84 | | | | | | |
| R^2^ / R^2^ adjusted | 0.872 / 0.866 | | | | | | |
| ** p<0.05   ** p<0.01   *** p<0.001* | | | | | | | |

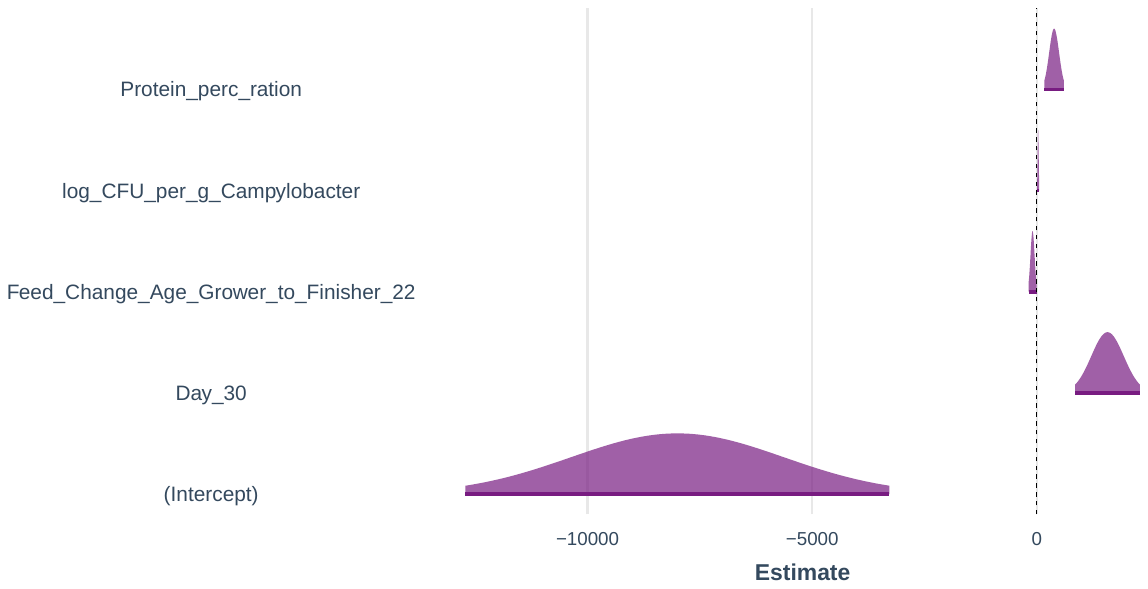

|  | **Richness – M3** | | | | | | |
| --- | --- | --- | --- | --- | --- | --- | --- |
| *Predictors* | *Estimates* | *std. Error* | *std. Beta* | *CI* | *standardized CI* | *Statistic* | *p* |
| (Intercept) | 7411.84056 | 6635.69881 |  | -5793.62091 – 20617.30203 |  | 1.11696 | 0.267 |
| Day_7 | -1269.79465 ^***^ | 361.73469 | -3.89111 | -1989.66962 – -549.91968 | -6.06370 – -1.71852 | -3.51029 | **0.001** |
| Protein_perc_ration | 144.96260 ^***^ | 37.03838 | 1.55319 | 71.25388 – 218.67132 | 0.77539 – 2.33098 | 3.91385 | **<0.001** |
| Energy_of_Ration | -6.51679 | 4.56951 | -1.43237 | -15.61041 – 2.57682 | -3.40089 – 0.53615 | -1.42615 | 0.158 |
| Observations | 84 | | | | | | |
| R^2^ / R^2^ adjusted | 0.865 / 0.860 | | | | | | |
| ** p<0.05   ** p<0.01   *** p<0.001* | | | | | | | |

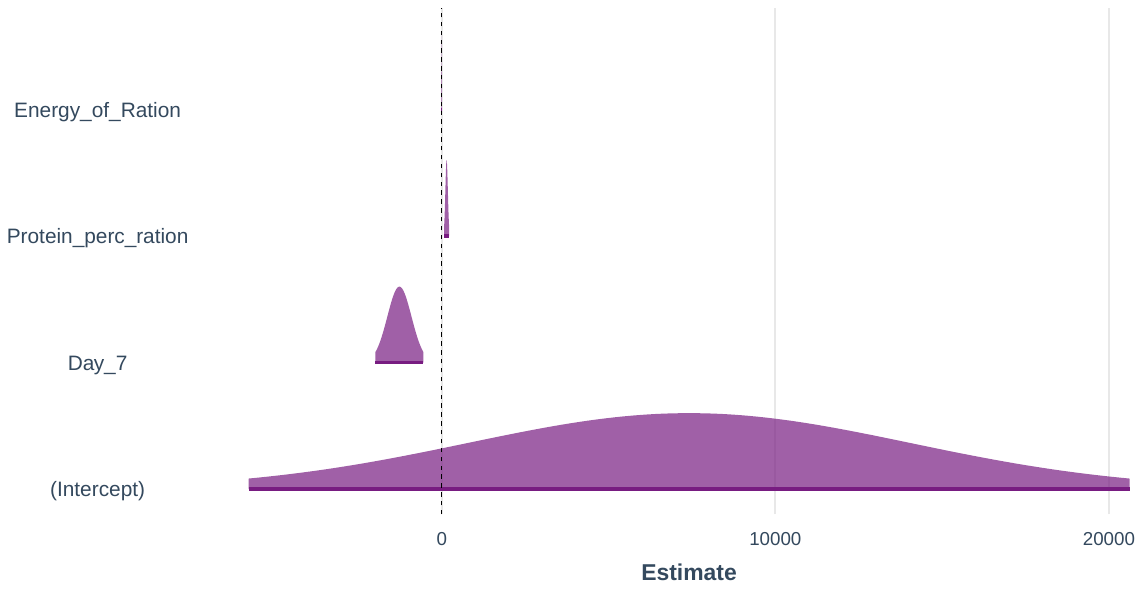

|  | **Richness – M5** | | | | | | |
| --- | --- | --- | --- | --- | --- | --- | --- |
| *Predictors* | *Estimates* | *std. Error* | *std. Beta* | *CI* | *standardized CI* | *Statistic* | *p* |
| (Intercept) | -4763.92283 | 8419.13193 |  | -21525.12696 – 11997.28130 |  | -0.56584 | 0.573 |
| Feed_N | -92.62051 | 46.86606 | -0.28162 | -185.92366 – 0.68264 | -0.56092 – -0.00233 | -1.97628 | 0.052 |
| Day_30 | 1683.70926 ^***^ | 458.91906 | 5.15949 | 770.07159 – 2597.34693 | 2.40321 – 7.91577 | 3.66886 | **<0.001** |
| log_CFU_per_g_Campylobacter | 25.72693 ^*^ | 12.29499 | 0.33987 | 1.24949 – 50.20437 | 0.02152 – 0.65821 | 2.09247 | **0.040** |
| Protein_perc_ration | 369.99931 ^**^ | 115.40128 | 3.96432 | 140.25301 – 599.74561 | 1.54091 – 6.38772 | 3.20620 | **0.002** |
| Energy_of_Ration | -2.13571 | 5.32939 | -0.46942 | -12.74571 – 8.47429 | -2.76529 – 1.82645 | -0.40074 | 0.690 |
| Observations | 84 | | | | | | |
| R^2^ / R^2^ adjusted | 0.872 / 0.864 | | | | | | |
| ** p<0.05   ** p<0.01   *** p<0.001* | | | | | | | |

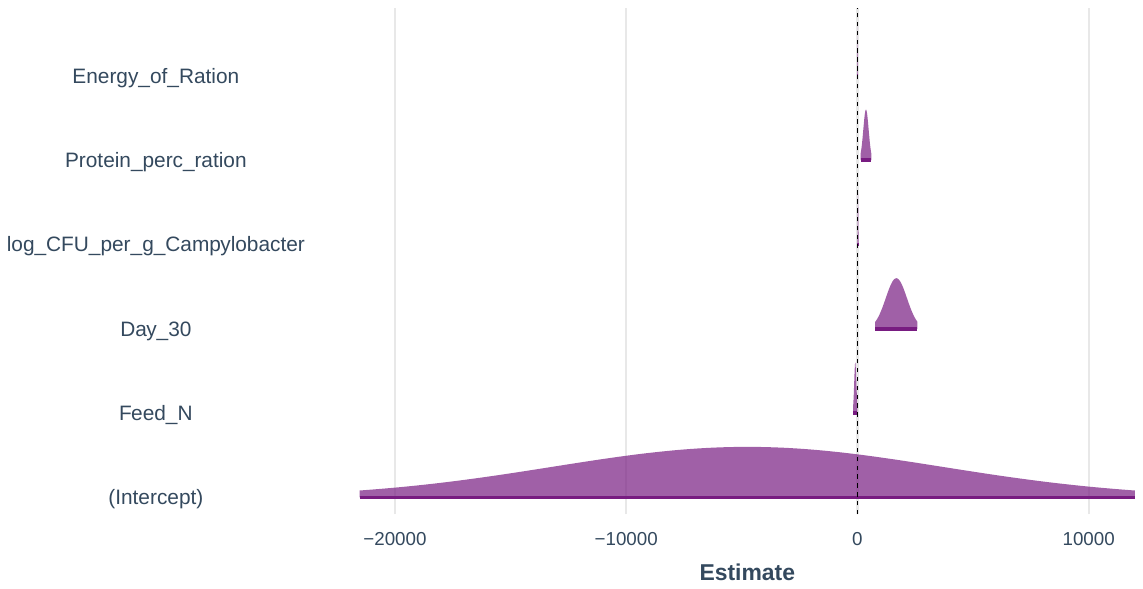

|  | **Richness – M6** | | | | | | |
| --- | --- | --- | --- | --- | --- | --- | --- |
| *Predictors* | *Estimates* | *std. Error* | *std. Beta* | *CI* | *standardized CI* | *Statistic* | *p* |
| (Intercept) | -1305.13476 | 9957.50912 |  | -21133.06856 – 18522.79904 |  | -0.13107 | 0.896 |
| Feed_N | -116.12951 | 80.66215 | -0.35311 | -276.74836 – 44.48935 | -0.83381 – 0.12760 | -1.43970 | 0.154 |
| Day_7 | -1795.65600 ^**^ | 556.91340 | -5.50253 | -2904.61225 – -686.69975 | -8.84737 – -2.15769 | -3.22430 | **0.002** |
| Total_Water_Consumption | 0.00059 | 0.00165 | 0.06772 | -0.00270 – 0.00389 | -0.30189 – 0.43734 | 0.35913 | 0.720 |
| log_CFU_per_g_Campylobacter | 25.72693 ^*^ | 12.36422 | 0.33987 | 1.10663 – 50.34724 | 0.01973 – 0.66000 | 2.08076 | **0.041** |
| Protein_perc_ration | 375.12560 ^**^ | 116.92563 | 4.01924 | 142.29691 – 607.95428 | 1.56382 – 6.47466 | 3.20824 | **0.002** |
| Energy_of_Ration | -3.44861 | 6.48752 | -0.75799 | -16.36690 – 9.46969 | -3.55278 – 2.03679 | -0.53158 | 0.597 |
| Observations | 84 | | | | | | |
| R^2^ / R^2^ adjusted | 0.873 / 0.863 | | | | | | |
| ** p<0.05   ** p<0.01   *** p<0.001* | | | | | | | |

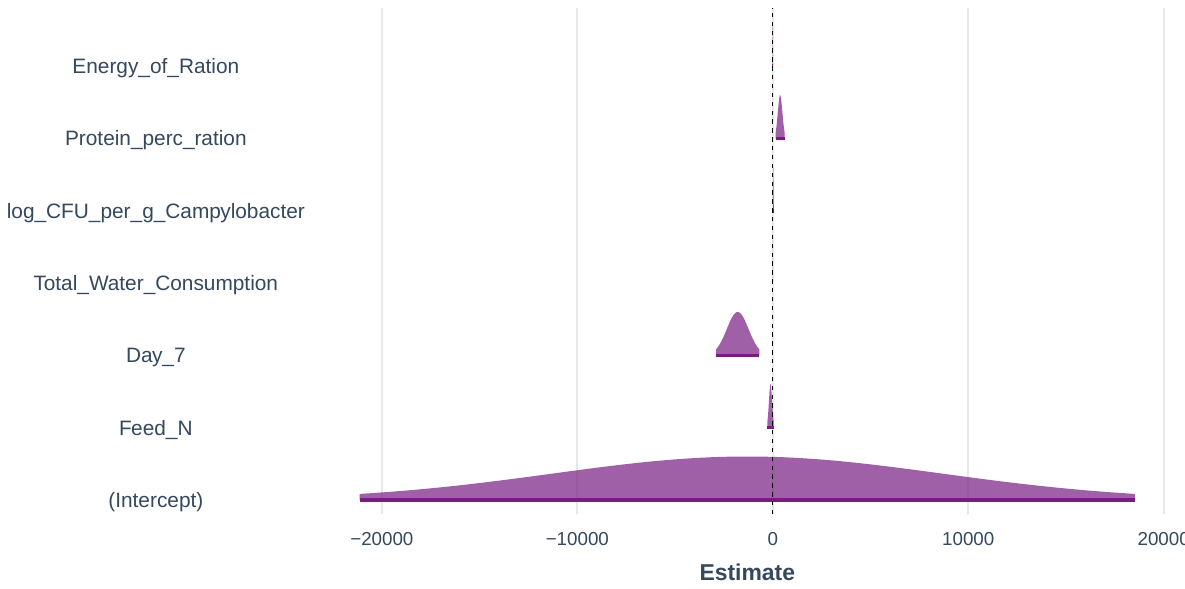

|  | **Richness – M1** | | | | | | |
| --- | --- | --- | --- | --- | --- | --- | --- |
| *Predictors* | *Estimates* | *std. Error* | *std. Beta* | *CI* | *standardized CI* | *Statistic* | *p* |
| (Intercept) | 253.89592 ^***^ | 8.68257 |  | 236.62351 – 271.16832 |  | 29.24201 | **<0.001** |
| Day_30 | 298.63698 ^***^ | 14.52872 | 0.91513 | 269.73472 – 327.53924 | 0.82787 – 1.00239 | 20.55493 | **<0.001** |
| Observations | 84 | | | | | | |
| R^2^ / R^2^ adjusted | 0.837 / 0.835 | | | | | | |
| ** p<0.05   ** p<0.01   *** p<0.001* | | | | | | | |

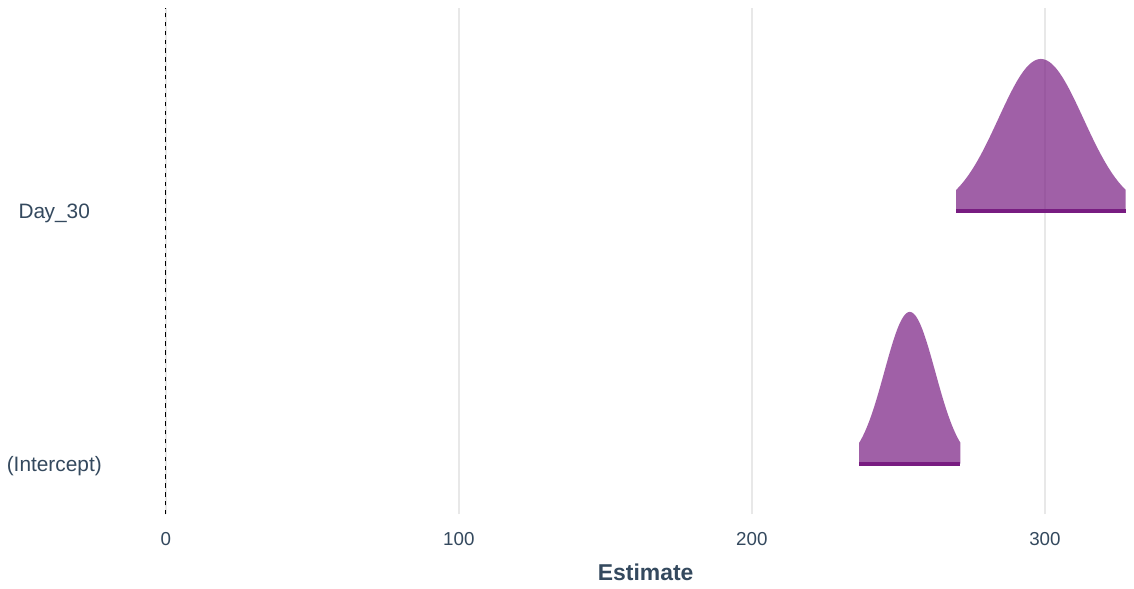
