## Supplementary S5-Shannon Entropy Subset Regression for "Impact of industrial production system parameters on chicken microbiomes: mechanisms to improve performance and reduce *Campylobacter*"

**Top models (Model parameters given afterwards with significant positive influencers highlighted in orange and negative in blue):**

|  | Model | Cross-validation Errors (5-Folds Average RMSE) |
| --- | --- | --- |
| M2 | Shannon ~ Protein_perc_ration + Energy_of_Ration | 0.39458 |
| M3 | Shannon ~ Day_7 + Weight_Gain_per_Day + Protein_perc_ration | 0.39658 |
| M4 | Shannon ~ Day_30 + Feed_Change_Age_Grower_to_Finisher_23 + log_CFU_per_g_Campylobacter + Protein_perc_ration | 0.39891 |
| M5 | Shannon ~ Day_30 + Feed_Change_Age_Grower_to_Finisher_23 + log_CFU_per_g_Campylobacter + Protein_perc_ration + Energy_of_Ration | 0.41012 |
| M6 | Shannon ~ Day_30 + Birds_Placed + Total_Water_Consumption + log_CFU_per_g_Campylobacter + Protein_perc_ration + Energy_of_Ration | 0.41396 |
| M1 | Shannon ~ Energy_of_Ration | 0.42019 |

|  | **Shannon – M2** | | | | | | |
| --- | --- | --- | --- | --- | --- | --- | --- |
| *Predictors* | *Estimates* | *std. Error* | *std. Beta* | *CI* | *standardized CI* | *Statistic* | *p* |
| (Intercept) | -92.38419 ^***^ | 20.54348 |  | -133.25927 – -51.50911 |  | -4.49701 | **<0.001** |
| Protein_perc_ration | 0.81695 ^***^ | 0.23067 | 1.91271 | 0.35799 – 1.27590 | 0.85421 – 2.97120 | 3.54166 | **0.001** |
| Energy_of_Ration | 0.05625 ^***^ | 0.01124 | 2.70165 | 0.03388 – 0.07862 | 1.64315 – 3.76014 | 5.00250 | **<0.001** |
| Observations | 84 | | | | | | |
| R^2^ / R^2^ adjusted | 0.690 / 0.683 | | | | | | |
| ** p<0.05   ** p<0.01   *** p<0.001* | | | | | | | |

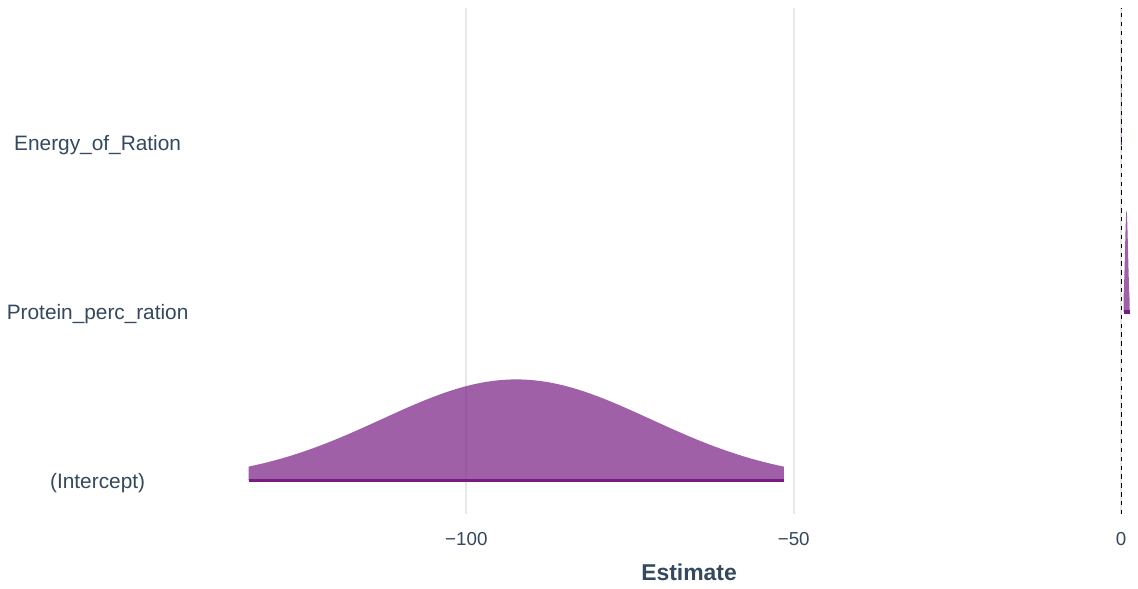

|  | **Shannon – M3** | | | | | | |
| --- | --- | --- | --- | --- | --- | --- | --- |
| *Predictors* | *Estimates* | *std. Error* | *std. Beta* | *CI* | *standardized CI* | *Statistic* | *p* |
| (Intercept) | -10.37590 ^*^ | 4.69614 |  | -19.72151 – -1.03028 |  | -2.20945 | **0.030** |
| Day_7 | -4.50042 ^***^ | 0.88067 | -3.01356 | -6.25300 – -2.74785 | -4.16937 – -1.85776 | -5.11026 | **<0.001** |
| Weight_Gain_per_Day | -40.40297 | 21.57871 | -0.11453 | -83.34597 – 2.54004 | -0.23443 – 0.00536 | -1.87235 | 0.065 |
| Protein_perc_ration | 0.95048 ^***^ | 0.25186 | 2.22535 | 0.44926 – 1.45170 | 1.06960 – 3.38109 | 3.77384 | **<0.001** |
| Observations | 84 | | | | | | |
| R^2^ / R^2^ adjusted | 0.702 / 0.690 | | | | | | |
| ** p<0.05   ** p<0.01   *** p<0.001* | | | | | | | |

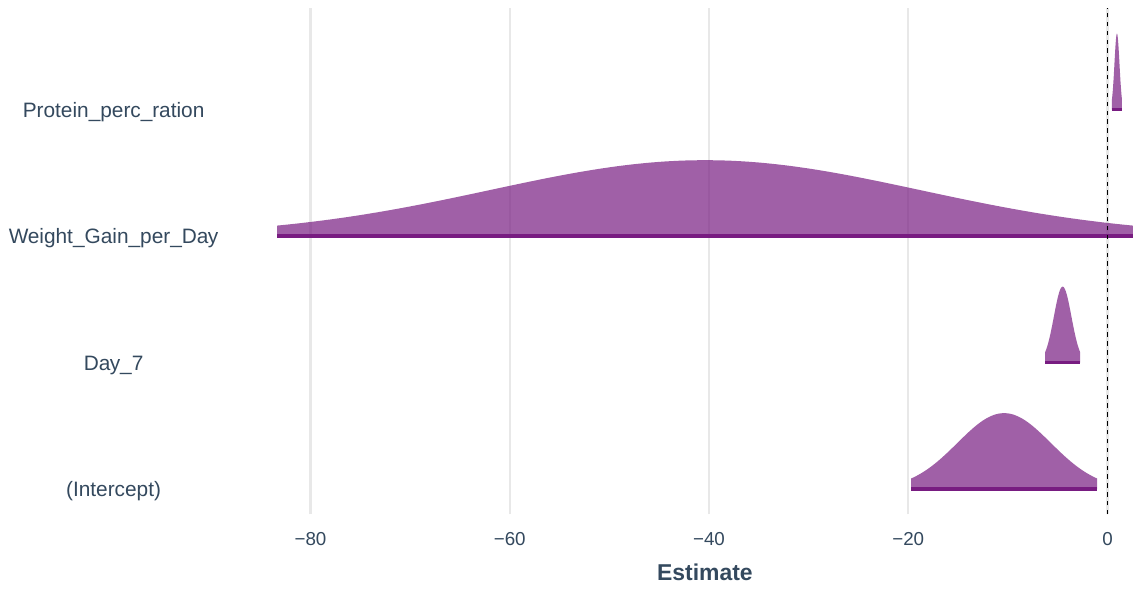

|  | **Shannon – M4** | | | | | | |
| --- | --- | --- | --- | --- | --- | --- | --- |
| *Predictors* | *Estimates* | *std. Error* | *std. Beta* | *CI* | *standardized CI* | *Statistic* | *p* |
| (Intercept) | -23.17967 ^**^ | 6.86977 |  | -36.85360 – -9.50574 |  | -3.37416 | **0.001** |
| Day_30 | 5.36982 ^***^ | 1.07307 | 3.59573 | 3.23392 – 7.50573 | 2.18740 – 5.00406 | 5.00415 | **<0.001** |
| Feed_Change_Age_Grower_to_Finisher_23 | 0.21442 | 0.11222 | 0.14126 | -0.00893 – 0.43778 | -0.00363 – 0.28615 | 1.91083 | 0.060 |
| log_CFU_per_g_Campylobacter | 0.02450 | 0.02988 | 0.07073 | -0.03497 – 0.08397 | -0.09831 – 0.23977 | 0.82008 | 0.415 |
| Protein_perc_ration | 1.21561 ^***^ | 0.31652 | 2.84608 | 0.58559 – 1.84562 | 1.39363 – 4.29854 | 3.84055 | **<0.001** |
| Observations | 84 | | | | | | |
| R^2^ / R^2^ adjusted | 0.702 / 0.687 | | | | | | |
| ** p<0.05   ** p<0.01   *** p<0.001* | | | | | | | |

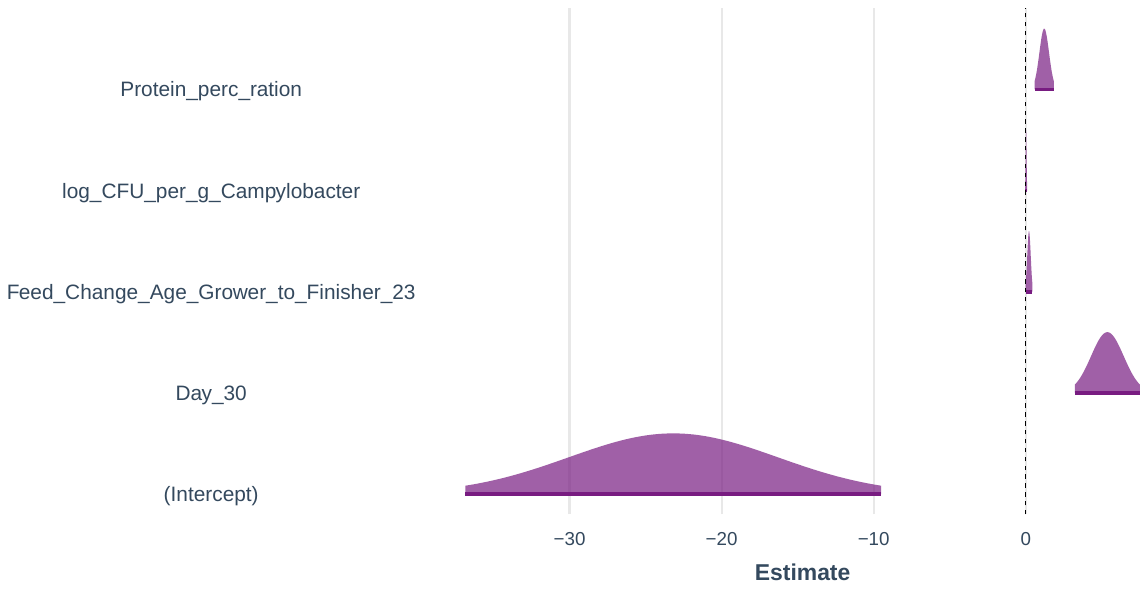

|  | **Shannon – M5** | | | | | | |
| --- | --- | --- | --- | --- | --- | --- | --- |
| *Predictors* | *Estimates* | *std. Error* | *std. Beta* | *CI* | *standardized CI* | *Statistic* | *p* |
| (Intercept) | -29.07548 | 59.35852 |  | -147.24922 – 89.09826 |  | -0.48983 | 0.626 |
| Day_30 | 5.03354 | 3.53180 | 3.37055 | -1.99774 – 12.06482 | -1.26468 – 8.00578 | 1.42520 | 0.158 |
| Feed_Change_Age_Grower_to_Finisher_23 | 0.20676 | 0.13646 | 0.13621 | -0.06490 – 0.47843 | -0.03998 – 0.31240 | 1.51525 | 0.134 |
| log_CFU_per_g_Campylobacter | 0.02532 | 0.03115 | 0.07308 | -0.03670 – 0.08733 | -0.10315 – 0.24931 | 0.81274 | 0.419 |
| Protein_perc_ration | 1.20921 ^***^ | 0.32488 | 2.83111 | 0.56243 – 1.85599 | 1.34031 – 4.32191 | 3.72207 | **<0.001** |
| Energy_of_Ration | 0.00436 | 0.04358 | 0.20935 | -0.08241 – 0.09113 | -3.89351 – 4.31220 | 0.10001 | 0.921 |
| Observations | 84 | | | | | | |
| R^2^ / R^2^ adjusted | 0.702 / 0.683 | | | | | | |
| ** p<0.05   ** p<0.01   *** p<0.001* | | | | | | | |

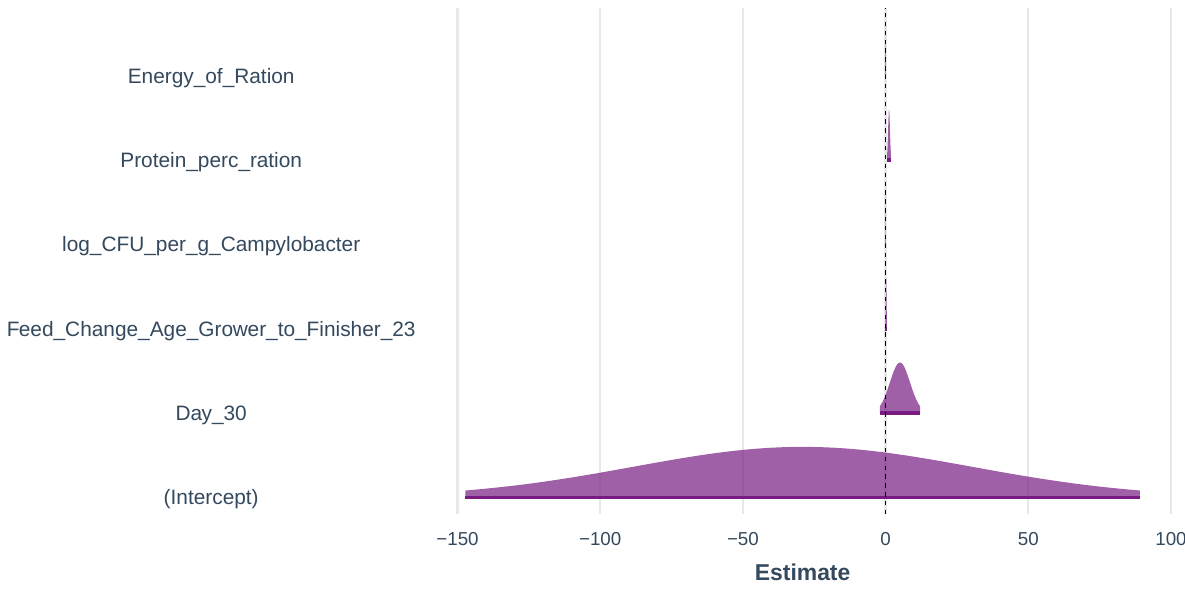

|  | **Shannon – M6** | | | | | | |
| --- | --- | --- | --- | --- | --- | --- | --- |
| *Predictors* | *Estimates* | *std. Error* | *std. Beta* | *CI* | *standardized CI* | *Statistic* | *p* |
| (Intercept) | -31.74641 | 67.58004 |  | -166.31545 – 102.82264 |  | -0.46976 | 0.640 |
| Day_30 | 5.12944 | 3.89717 | 3.43476 | -2.63083 – 12.88970 | -1.67999 – 8.54951 | 1.31619 | 0.192 |
| Birds_Placed | 0.00020 | 0.00015 | 0.60711 | -0.00010 – 0.00049 | -0.28964 – 1.50385 | 1.32692 | 0.188 |
| Total_Water_Consumption | -0.00003 | 0.00002 | -0.64033 | -0.00007 – 0.00002 | -1.81134 – 0.53067 | -1.07176 | 0.287 |
| log_CFU_per_g_Campylobacter | 0.03016 | 0.08652 | 0.08705 | -0.14213 – 0.20244 | -0.40249 – 0.57659 | 0.34852 | 0.728 |
| Protein_perc_ration | 1.25423 | 0.81822 | 2.93651 | -0.37506 – 2.88352 | -0.81819 – 6.69120 | 1.53287 | 0.129 |
| Energy_of_Ration | 0.00506 | 0.04540 | 0.24307 | -0.08534 – 0.09546 | -4.03057 – 4.51671 | 0.11148 | 0.912 |
| Observations | 84 | | | | | | |
| R^2^ / R^2^ adjusted | 0.702 / 0.679 | | | | | | |
| ** p<0.05   ** p<0.01   *** p<0.001* | | | | | | | |

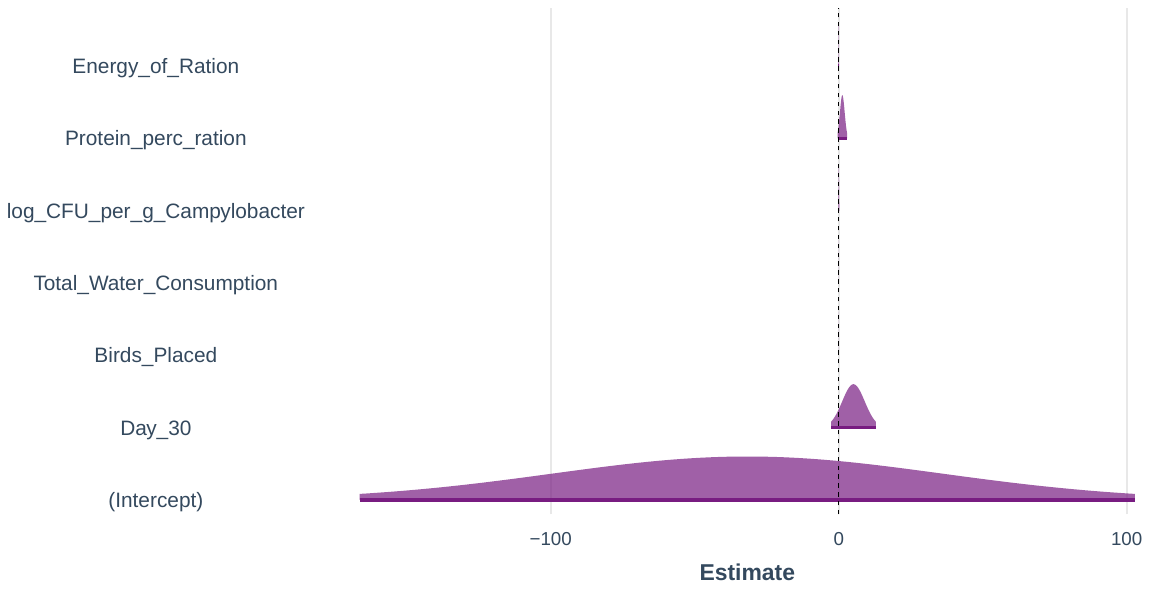

|  | **Shannon – M1** | | | | | | |
| --- | --- | --- | --- | --- | --- | --- | --- |
| *Predictors* | *Estimates* | *std. Error* | *std. Beta* | *CI* | *standardized CI* | *Statistic* | *p* |
| (Intercept) | -19.91118 ^***^ | 1.93998 |  | -23.77043 – -16.05194 |  | -10.26359 | **<0.001** |
| Energy_of_Ration | 0.01669 ^***^ | 0.00137 | 0.80152 | 0.01395 – 0.01942 | 0.67209 – 0.93094 | 12.13768 | **<0.001** |
| Observations | 84 | | | | | | |
| R^2^ / R^2^ adjusted | 0.642 / 0.638 | | | | | | |
| ** p<0.05   ** p<0.01   *** p<0.001* | | | | | | | |

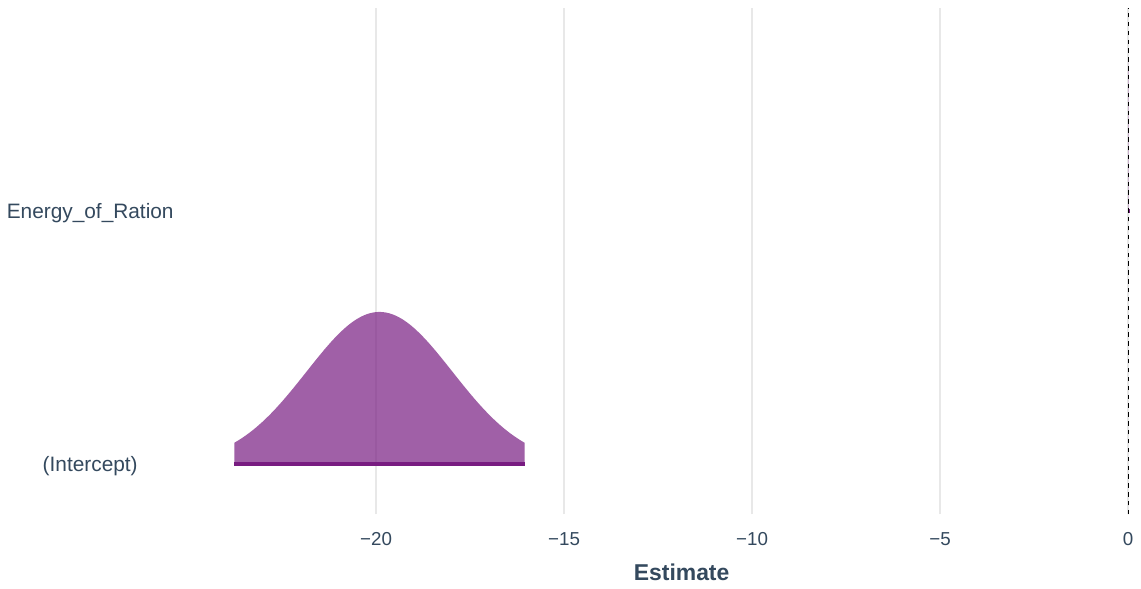
