## Supplementary S6-NRI Subset Regression for "Impact of industrial production system parameters on chicken microbiomes: mechanisms to improve performance and reduce *Campylobacter*"

**Top models (Model parameters given afterwards with significant positive influencers highlighted in orange and negative in blue):**

|  | Model | Cross-validation Errors (5-Folds Average RMSE) |
| --- | --- | --- |
| M1 | NRI ~ Protein_perc_ration | 1.99595 |
| M3 | NRI ~ Feed_Change_Age_Finisher_to_Withdrawal_34 + log_CFU_per_g_Campylobacter + Protein_perc_ration | 2.00044 |
| M4 | NRI ~ Feed_N + Feed_HW + log_CFU_per_g_Campylobacter + Protein_perc_ration | 2.00251 |
| M5 | NRI ~ Feed_N + Feed_HW + log_CFU_per_g_Campylobacter + Protein_perc_ration + Energy_of_Ration | 2.00776 |
| M6 | NRI ~ Day_7 + Birds_Killed + Feed_Change_Age_Grower_to_Finisher_22 + log_CFU_per_g_Campylobacter + Protein_perc_ration + Energy_of_Ration | 2.01021 |
| M2 | NRI ~ Water_Consumption_per_Bird + log_CFU_per_g_Campylobacter | 2.01483 |

|  | **NRI – M1** | | | | | | |
| --- | --- | --- | --- | --- | --- | --- | --- |
| *Predictors* | *Estimates* | *std. Error* | *std. Beta* | *CI* | *standardized CI* | *Statistic* | *p* |
| (Intercept) | 12.62945 ^***^ | 2.74327 |  | 7.17221 – 18.08670 |  | 4.60379 | **<0.001** |
| Protein_perc_ration | -0.42926 ^**^ | 0.13401 | -0.33349 | -0.69584 – -0.16267 | -0.53754 – -0.12944 | -3.20322 | **0.002** |
| Observations | 84 | | | | | | |
| R^2^ / R^2^ adjusted | 0.111 / 0.100 | | | | | | |
| ** p<0.05   ** p<0.01   *** p<0.001* | | | | | | | |

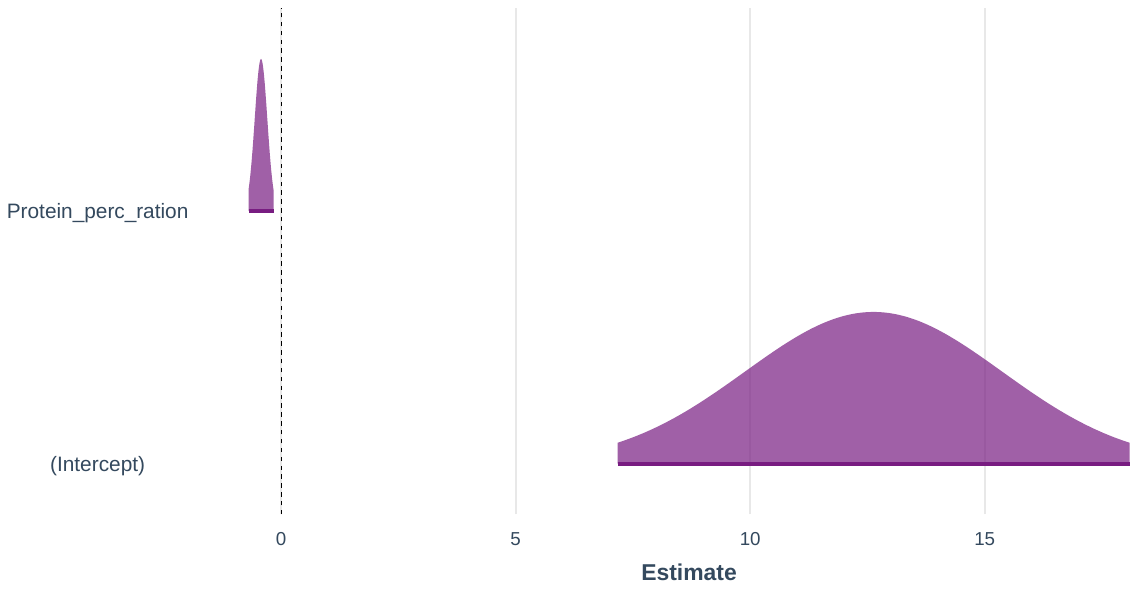

|  | **NRI – M3** | | | | | | |
| --- | --- | --- | --- | --- | --- | --- | --- |
| *Predictors* | *Estimates* | *std. Error* | *std. Beta* | *CI* | *standardized CI* | *Statistic* | *p* |
| (Intercept) | 8.70358 ^*^ | 3.32326 |  | 2.09008 – 15.31709 |  | 2.61899 | **0.011** |
| Feed_Change_Age_Finisher_to_Withdrawal_34 | -1.92001 ^**^ | 0.56685 | -0.42331 | -3.04808 – -0.79194 | -0.66826 – -0.17836 | -3.38716 | **0.001** |
| log_CFU_per_g_Campylobacter | 0.28137 | 0.15137 | 0.26952 | -0.01986 – 0.58261 | -0.01466 – 0.55371 | 1.85886 | 0.067 |
| Protein_perc_ration | -0.21449 | 0.16305 | -0.16663 | -0.53897 – 0.10999 | -0.41491 – 0.08164 | -1.31548 | 0.192 |
| Observations | 84 | | | | | | |
| R^2^ / R^2^ adjusted | 0.223 / 0.194 | | | | | | |
| ** p<0.05   ** p<0.01   *** p<0.001* | | | | | | | |

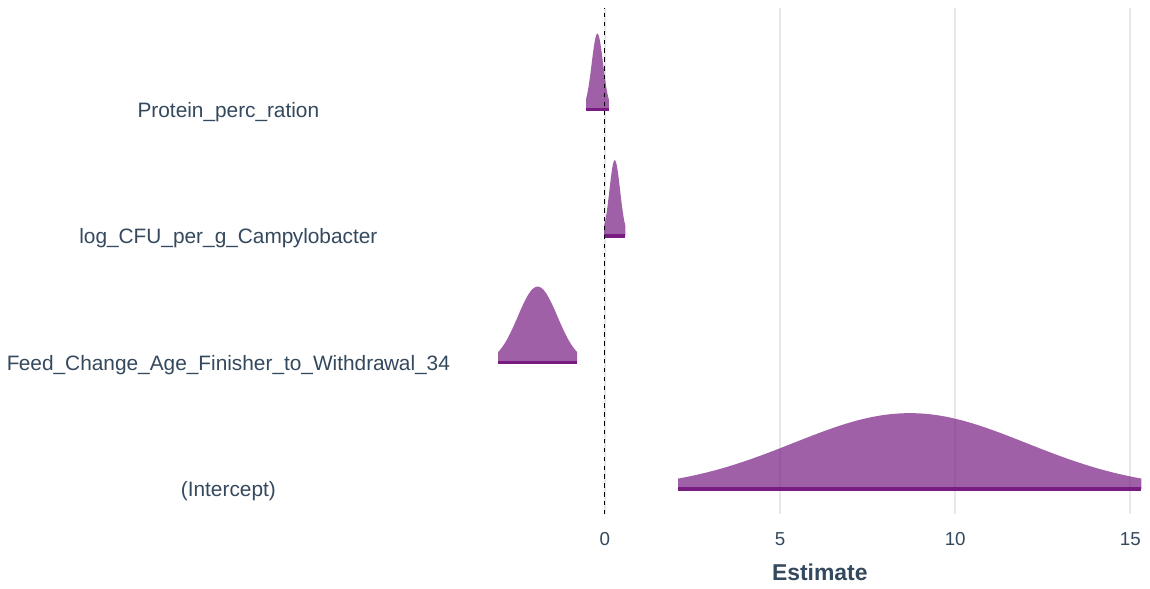

|  | **NRI – M4** | | | | | | |
| --- | --- | --- | --- | --- | --- | --- | --- |
| *Predictors* | *Estimates* | *std. Error* | *std. Beta* | *CI* | *standardized CI* | *Statistic* | *p* |
| (Intercept) | 8.58609 ^*^ | 3.35023 |  | 1.91761 – 15.25456 |  | 2.56283 | **0.012** |
| Feed_N | -1.79923 ^**^ | 0.63012 | -0.39668 | -3.05346 – -0.54500 | -0.66897 – -0.12439 | -2.85535 | **0.005** |
| Feed_HW | 0.23681 | 0.52791 | 0.05177 | -0.81397 – 1.28759 | -0.17441 – 0.27795 | 0.44859 | 0.655 |
| log_CFU_per_g_Campylobacter | 0.28129 | 0.15213 | 0.26944 | -0.02152 – 0.58410 | -0.01617 – 0.55505 | 1.84898 | 0.068 |
| Protein_perc_ration | -0.21464 | 0.16387 | -0.16675 | -0.54081 – 0.11154 | -0.41627 – 0.08277 | -1.30981 | 0.194 |
| Observations | 84 | | | | | | |
| R^2^ / R^2^ adjusted | 0.225 / 0.186 | | | | | | |
| ** p<0.05   ** p<0.01   *** p<0.001* | | | | | | | |

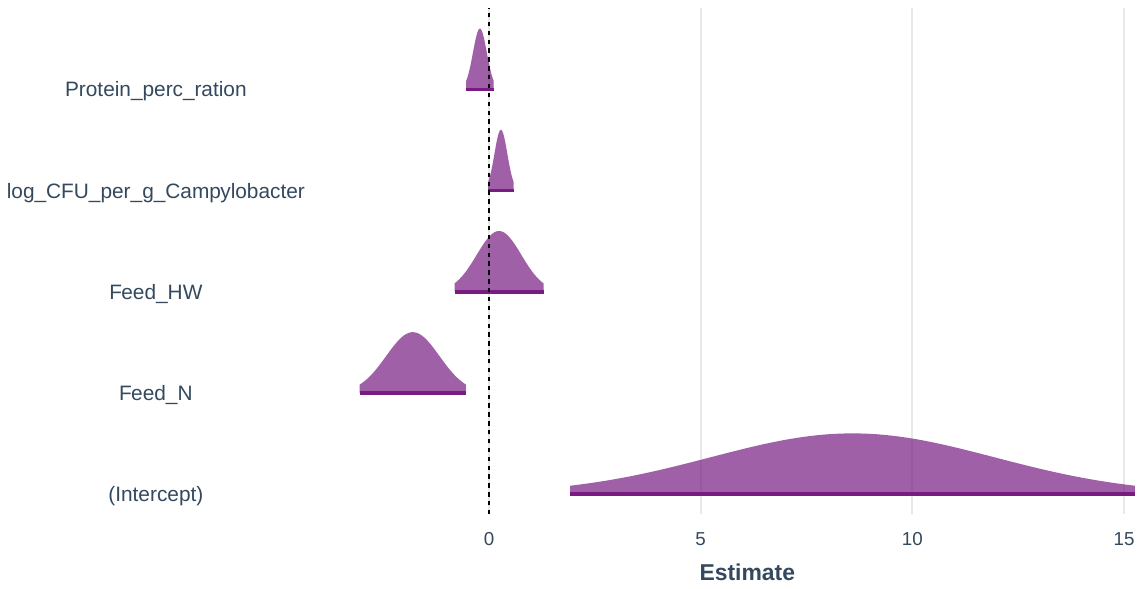

|  | **NRI – M5** | | | | | | |
| --- | --- | --- | --- | --- | --- | --- | --- |
| *Predictors* | *Estimates* | *std. Error* | *std. Beta* | *CI* | *standardized CI* | *Statistic* | *p* |
| (Intercept) | 59.58578 | 277.11862 |  | -492.11501 – 611.28658 |  | 0.21502 | 0.830 |
| Feed_N | -1.55742 | 1.45881 | -0.34337 | -4.46169 – 1.34685 | -0.97375 – 0.28701 | -1.06759 | 0.289 |
| Feed_HW | 0.26440 | 0.55191 | 0.05780 | -0.83437 – 1.36318 | -0.17867 – 0.29426 | 0.47907 | 0.633 |
| log_CFU_per_g_Campylobacter | 0.21491 | 0.39180 | 0.20586 | -0.56510 – 0.99492 | -0.52971 – 0.94143 | 0.54852 | 0.585 |
| Protein_perc_ration | -0.81993 | 3.29287 | -0.63699 | -7.37552 – 5.73566 | -5.65097 – 4.37698 | -0.24900 | 0.804 |
| Energy_of_Ration | -0.02743 | 0.14904 | -0.43718 | -0.32415 – 0.26929 | -5.09275 – 4.21840 | -0.18405 | 0.854 |
| Observations | 84 | | | | | | |
| R^2^ / R^2^ adjusted | 0.226 / 0.176 | | | | | | |
| ** p<0.05   ** p<0.01   *** p<0.001* | | | | | | | |

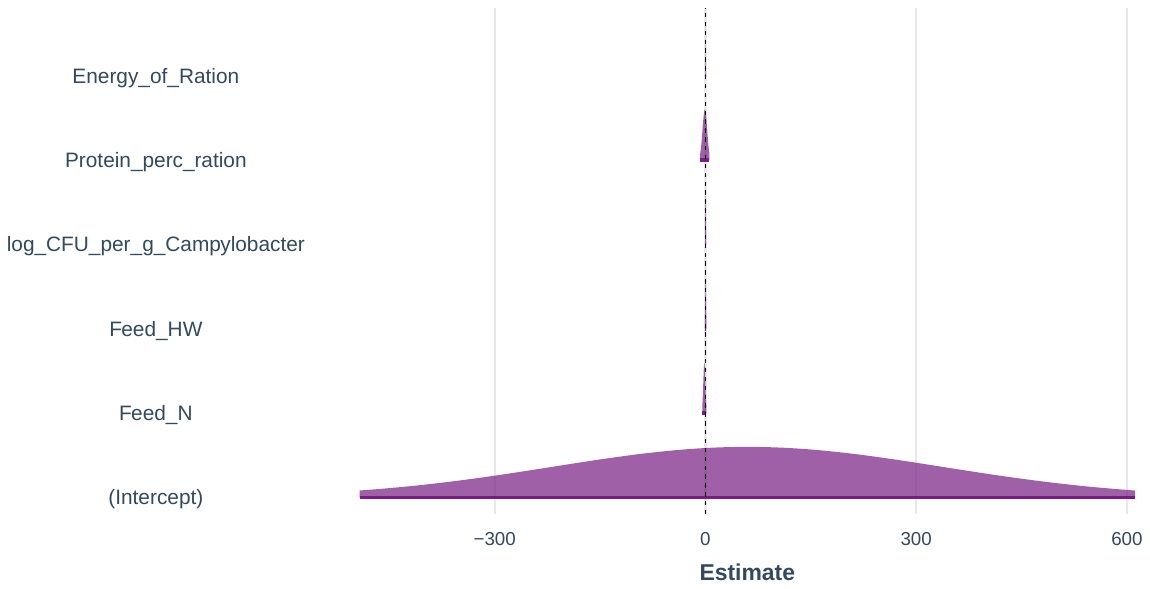

|  | **NRI – M6** | | | | | | |
| --- | --- | --- | --- | --- | --- | --- | --- |
| *Predictors* | *Estimates* | *std. Error* | *std. Beta* | *CI* | *standardized CI* | *Statistic* | *p* |
| (Intercept) | 75.29773 | 338.70584 |  | -599.15176 – 749.74722 |  | 0.22231 | 0.825 |
| Day_7 | -1.68924 | 18.94286 | -0.37534 | -39.40930 – 36.03081 | -8.62481 – 7.87413 | -0.08918 | 0.929 |
| Birds_Killed | 0.00014 | 0.00031 | 0.14298 | -0.00049 – 0.00077 | -0.48764 – 0.77359 | 0.44437 | 0.658 |
| Feed_Change_Age_Grower_to_Finisher_22 | -2.36484 | 2.22217 | -0.52138 | -6.78974 – 2.06007 | -1.48162 – 0.43886 | -1.06420 | 0.291 |
| log_CFU_per_g_Campylobacter | 0.22795 | 0.42056 | 0.21835 | -0.60949 – 1.06538 | -0.57121 – 1.00791 | 0.54202 | 0.589 |
| Protein_perc_ration | -0.62385 | 3.97711 | -0.48466 | -8.54329 – 7.29559 | -6.54052 – 5.57120 | -0.15686 | 0.876 |
| Energy_of_Ration | -0.04186 | 0.22067 | -0.66720 | -0.48127 – 0.39754 | -7.56005 – 6.22566 | -0.18972 | 0.850 |
| Observations | 84 | | | | | | |
| R^2^ / R^2^ adjusted | 0.226 / 0.165 | | | | | | |
| ** p<0.05   ** p<0.01   *** p<0.001* | | | | | | | |

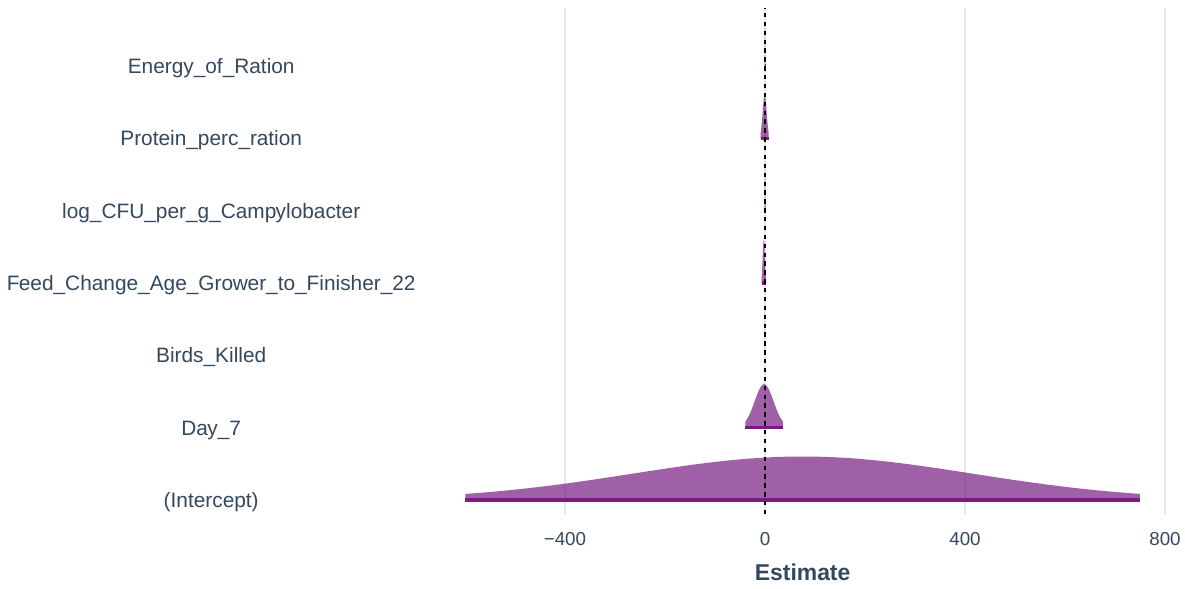

|  | **NRI – M2** | | | | | | |
| --- | --- | --- | --- | --- | --- | --- | --- |
| *Predictors* | *Estimates* | *std. Error* | *std. Beta* | *CI* | *standardized CI* | *Statistic* | *p* |
| (Intercept) | 22.87362 ^***^ | 4.40551 |  | 14.10804 – 31.63920 |  | 5.19205 | **<0.001** |
| Water_Consumption_per_Bird | -3.29095 ^***^ | 0.75748 | -0.49338 | -4.79809 – -1.78380 | -0.71596 – -0.27081 | -4.34461 | **<0.001** |
| log_CFU_per_g_Campylobacter | 0.40605 ^***^ | 0.11856 | 0.38895 | 0.17016 – 0.64194 | 0.16637 – 0.61153 | 3.42500 | **0.001** |
| Observations | 84 | | | | | | |
| R^2^ / R^2^ adjusted | 0.207 / 0.187 | | | | | | |
| ** p<0.05   ** p<0.01   *** p<0.001* | | | | | | | |

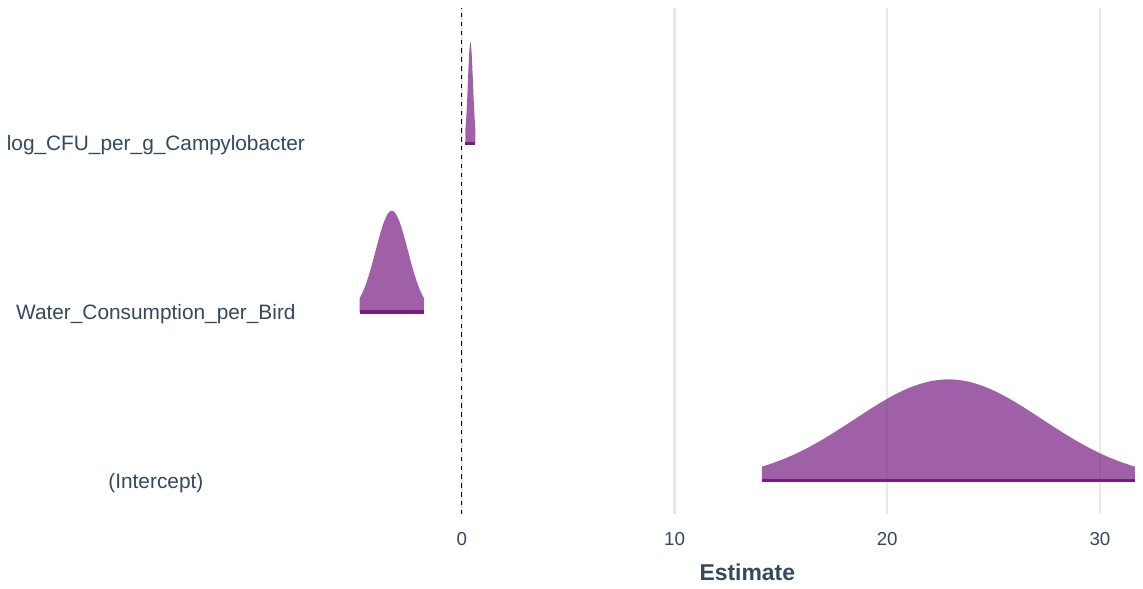
