## Supplementary S7-NTI Subset Regression for "Impact of industrial production system parameters on chicken microbiomes: mechanisms to improve performance and reduce *Campylobacter*"

**Top models (Model parameters given afterwards with significant positive influencers highlighted in orange and negative in blue):**

|  | Model | Cross-validation Errors (5-Folds Average RMSE) |
| --- | --- | --- |
| M3 | NTI ~ Day_7 + Age_At_Thin + Protein_perc_ration | 1.60290 |
| M5 | NTI ~ Day_30 + Total_Water_Consumption + log_CFU_per_g_Campylobacter + Protein_perc_ration + Energy_of_Ration | 1.60347 |
| M4 | NTI ~ Day_7 + Placement_Birds_per_m2 + log_CFU_per_g_Campylobacter + Protein_perc_ration | 1.60392 |
| M6 | NTI ~ Feed_N + Feed_HW + Day_7 + log_CFU_per_g_Campylobacter + Protein_perc_ration + Energy_of_Ration | 1.60769 |
| M2 | NTI ~ Food_Conversion_Ratio + log_CFU_per_g_Campylobacter | 1.64608 |
| M1 | NTI ~ Hockmark_percentage | 1.80411 |

|  | **NTI – M3** | | | | | | |
| --- | --- | --- | --- | --- | --- | --- | --- |
| *Predictors* | *Estimates* | *std. Error* | *std. Beta* | *CI* | *standardized CI* | *Statistic* | *p* |
| (Intercept) | 86.98727 ^***^ | 18.44338 |  | 50.28378 – 123.69076 |  | 4.71645 | **<0.001** |
| Day_7 | 10.04552 ^*^ | 3.86552 | 2.39868 | 2.35290 – 17.73815 | 0.58961 – 4.20775 | 2.59875 | **0.011** |
| Age_At_Thin | -0.61536 ^***^ | 0.16508 | -0.35767 | -0.94387 – -0.28685 | -0.54572 – -0.16962 | -3.72776 | **<0.001** |
| Protein_perc_ration | -3.26620 ^**^ | 1.10767 | -2.72691 | -5.47054 – -1.06187 | -4.53944 – -0.91437 | -2.94871 | **0.004** |
| Observations | 84 | | | | | | |
| R^2^ / R^2^ adjusted | 0.397 / 0.374 | | | | | | |
| ** p<0.05   ** p<0.01   *** p<0.001* | | | | | | | |

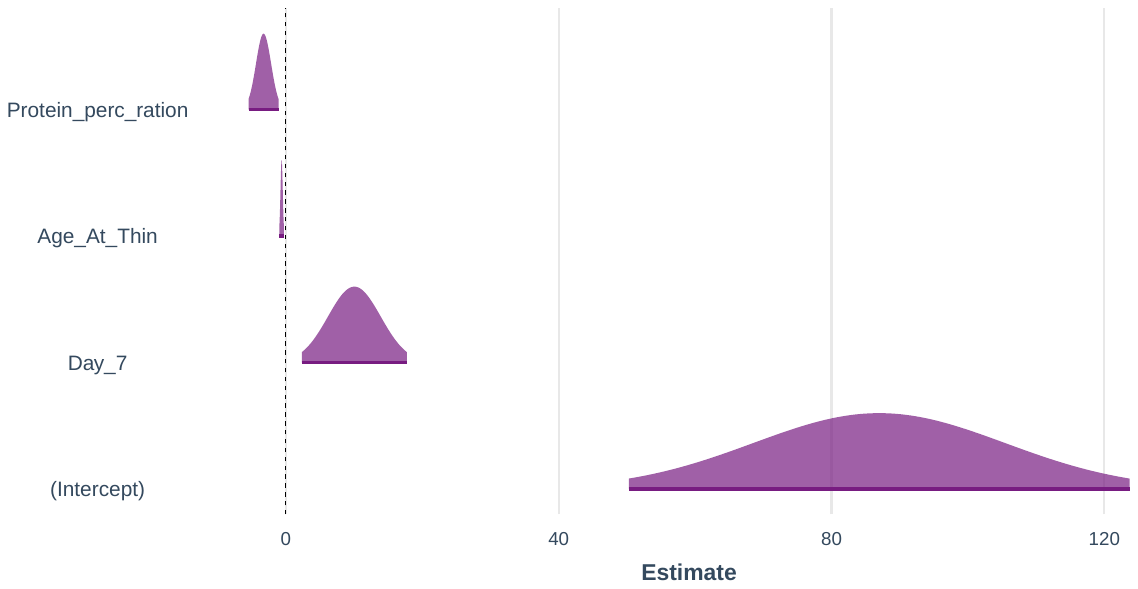

|  | **NTI – M5** | | | | | | |
| --- | --- | --- | --- | --- | --- | --- | --- |
| *Predictors* | *Estimates* | *std. Error* | *std. Beta* | *CI* | *standardized CI* | *Statistic* | *p* |
| (Intercept) | 2.43091 | 265.07201 |  | -525.28693 – 530.14874 |  | 0.00917 | 0.993 |
| Day_30 | -9.10195 | 10.64117 | -2.17337 | -30.28689 – 12.08300 | -7.15345 – 2.80671 | -0.85535 | 0.395 |
| Total_Water_Consumption | -0.00005 | 0.00003 | -0.46403 | -0.00011 – 0.00000 | -0.93052 – 0.00245 | -1.94967 | 0.055 |
| log_CFU_per_g_Campylobacter | 0.13477 | 0.28696 | 0.13873 | -0.43652 – 0.70607 | -0.44023 – 0.71770 | 0.46965 | 0.640 |
| Protein_perc_ration | -2.11967 | 2.48137 | -1.76968 | -7.05970 – 2.82036 | -5.83007 – 2.29070 | -0.85423 | 0.396 |
| Energy_of_Ration | 0.03847 | 0.16938 | 0.65894 | -0.29874 – 0.37568 | -5.02686 – 6.34475 | 0.22715 | 0.821 |
| Observations | 84 | | | | | | |
| R^2^ / R^2^ adjusted | 0.398 / 0.360 | | | | | | |
| ** p<0.05   ** p<0.01   *** p<0.001* | | | | | | | |

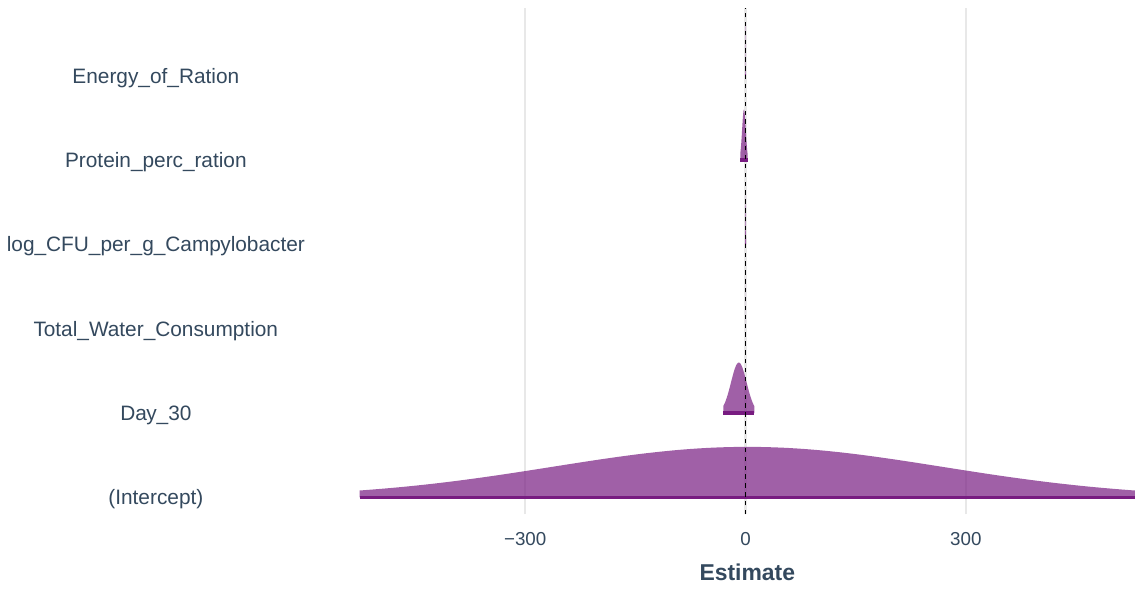

|  | **NTI – M4** | | | | | | |
| --- | --- | --- | --- | --- | --- | --- | --- |
| *Predictors* | *Estimates* | *std. Error* | *std. Beta* | *CI* | *standardized CI* | *Statistic* | *p* |
| (Intercept) | 54.29686 | 36.98352 |  | -19.31701 – 127.91072 |  | 1.46814 | 0.146 |
| Day_7 | 6.91297 | 7.07798 | 1.65068 | -7.17539 – 21.00133 | -1.66182 – 4.96319 | 0.97669 | 0.332 |
| Placement_Birds_per_m2 | -0.34522 ^*^ | 0.15913 | -0.44538 | -0.66196 – -0.02848 | -0.84775 – -0.04300 | -2.16944 | **0.033** |
| log_CFU_per_g_Campylobacter | 0.10333 | 0.22028 | 0.10636 | -0.33513 – 0.54178 | -0.33807 – 0.55079 | 0.46907 | 0.640 |
| Protein_perc_ration | -2.30223 | 2.14685 | -1.92210 | -6.57543 – 1.97096 | -5.43509 – 1.59088 | -1.07238 | 0.287 |
| Observations | 84 | | | | | | |
| R^2^ / R^2^ adjusted | 0.398 / 0.368 | | | | | | |
| ** p<0.05   ** p<0.01   *** p<0.001* | | | | | | | |

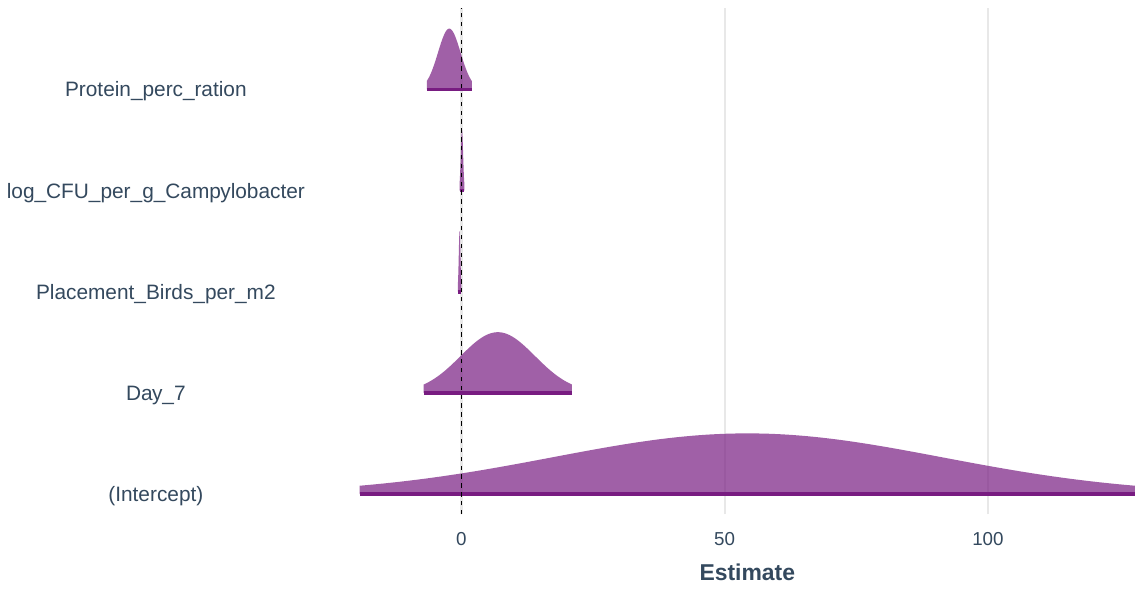

|  | **NTI – M6** | | | | | | |
| --- | --- | --- | --- | --- | --- | --- | --- |
| *Predictors* | *Estimates* | *std. Error* | *std. Beta* | *CI* | *standardized CI* | *Statistic* | *p* |
| (Intercept) | -10.13034 | 279.42571 |  | -566.53801 – 546.27733 |  | -0.03625 | 0.971 |
| Feed_N | -2.20480 | 1.32035 | -0.52239 | -4.83395 – 0.42435 | -1.13552 – 0.09075 | -1.66986 | 0.099 |
| Feed_HW | -0.62071 | 0.54981 | -0.14581 | -1.71552 – 0.47411 | -0.39896 – 0.10733 | -1.12895 | 0.262 |
| Day_7 | 9.06174 | 15.53797 | 2.16377 | -21.87832 – 40.00179 | -5.10803 – 9.43556 | 0.58320 | 0.561 |
| log_CFU_per_g_Campylobacter | 0.13545 | 0.34496 | 0.13943 | -0.55146 – 0.82236 | -0.55656 – 0.83541 | 0.39264 | 0.696 |
| Protein_perc_ration | -2.11218 | 3.26224 | -1.76342 | -8.60813 – 4.38378 | -7.10158 – 3.57473 | -0.64746 | 0.519 |
| Energy_of_Ration | 0.03826 | 0.18100 | 0.65522 | -0.32217 – 0.39868 | -5.42074 – 6.73118 | 0.21136 | 0.833 |
| Observations | 84 | | | | | | |
| R^2^ / R^2^ adjusted | 0.398 / 0.351 | | | | | | |
| ** p<0.05   ** p<0.01   *** p<0.001* | | | | | | | |

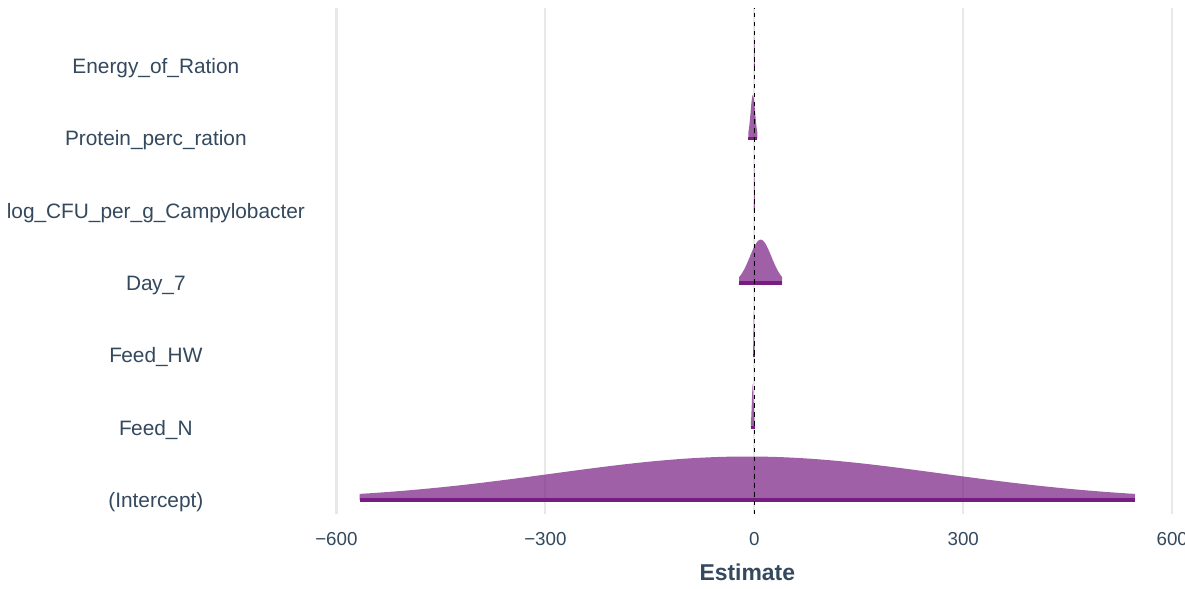

|  | **NTI – M2** | | | | | | |
| --- | --- | --- | --- | --- | --- | --- | --- |
| *Predictors* | *Estimates* | *std. Error* | *std. Beta* | *CI* | *standardized CI* | *Statistic* | *p* |
| (Intercept) | -21.51722 ^***^ | 3.91548 |  | -29.30781 – -13.72664 |  | -5.49542 | **<0.001** |
| Food_Conversion_Ratio | 17.23651 ^***^ | 2.48274 | 0.69699 | 12.29662 – 22.17639 | 0.50022 – 0.89376 | 6.94252 | **<0.001** |
| log_CFU_per_g_Campylobacter | 0.41591 ^***^ | 0.09753 | 0.42813 | 0.22186 – 0.60996 | 0.23136 – 0.62490 | 4.26452 | **<0.001** |
| Observations | 84 | | | | | | |
| R^2^ / R^2^ adjusted | 0.378 / 0.363 | | | | | | |
| ** p<0.05   ** p<0.01   *** p<0.001* | | | | | | | |

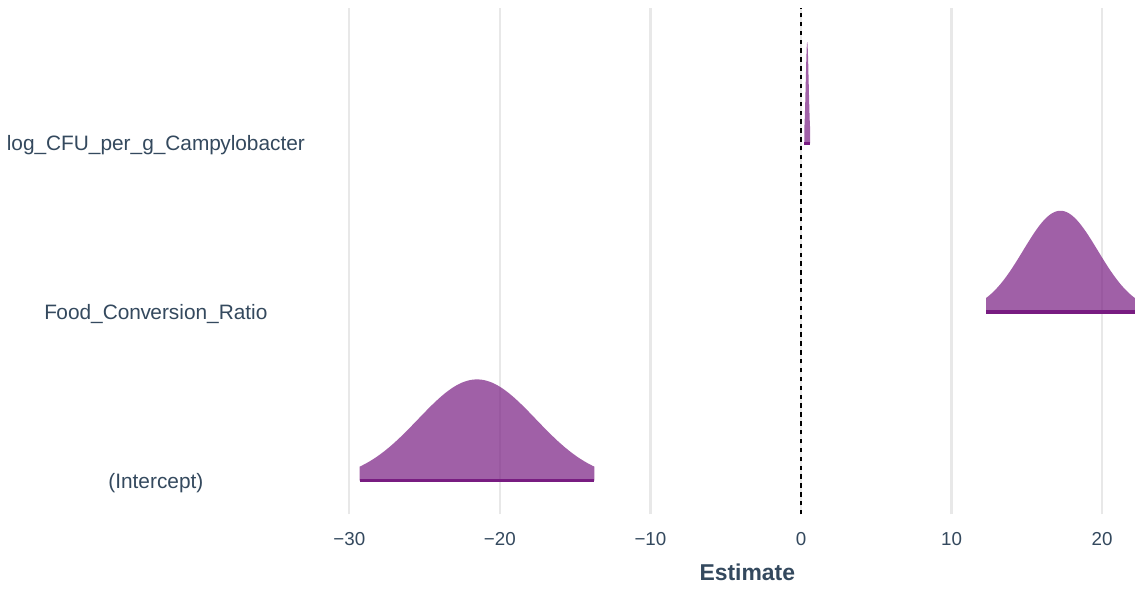

|  | **NTI – M1** | | | | | | |
| --- | --- | --- | --- | --- | --- | --- | --- |
| *Predictors* | *Estimates* | *std. Error* | *std. Beta* | *CI* | *standardized CI* | *Statistic* | *p* |
| (Intercept) | 6.82797 ^***^ | 0.29417 |  | 6.24276 – 7.41317 |  | 23.21057 | **<0.001** |
| Hockmark_percentage | -0.11158 ^***^ | 0.02191 | -0.49015 | -0.15517 – -0.06799 | -0.67881 – -0.30149 | -5.09217 | **<0.001** |
| Observations | 84 | | | | | | |
| R^2^ / R^2^ adjusted | 0.240 / 0.231 | | | | | | |
| ** p<0.05   ** p<0.01   *** p<0.001* | | | | | | | |

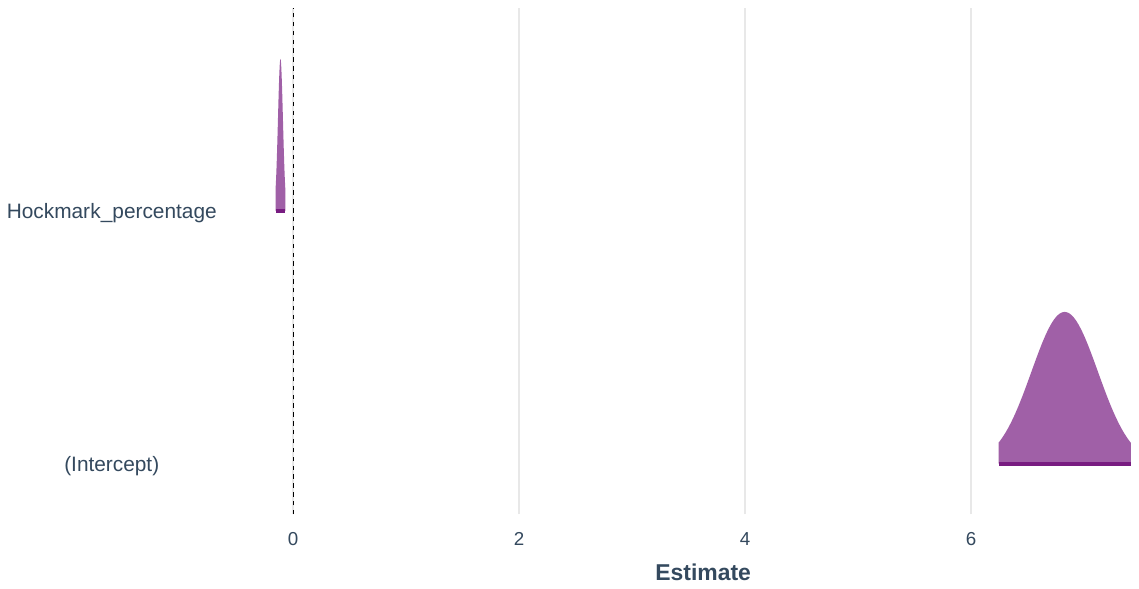
