## Supplementary S8-LCBD Bray-Curtis Subset Regression for "Impact of industrial production system parameters on chicken microbiomes: mechanisms to improve performance and reduce *Campylobacter*"

**Top models (Model parameters given afterwards with significant positive influencers highlighted in orange and negative in blue):**

|  | Model | Cross-validation Errors (5-Folds Average RMSE) |
| --- | --- | --- |
| M2 | LCBD ~ log_CFU_per_g_Campylobacter + Protein_perc_ration | 0.00138 |
| M4 | LCBD ~ Day_7 + Total_Mortality_percentage + Protein_perc_ration + Energy_of_Ration | 0.00140 |
| M3 | LCBD ~ Day_7 + Total_Mortality_percentage + Protein_perc_ration | 0.00141 |
| M1 | LCBD ~ Protein_perc_ration | 0.00141 |
| M5 | LCBD ~ Day_7 + No_of_Parent_Flocks_Used_3 + log_CFU_per_g_Campylobacter + Protein_perc_ration + Energy_of_Ration | 0.00142 |
| M6 | LCBD ~ Feed_N + Feed_HW + Day_7 + log_CFU_per_g_Campylobacter + Protein_perc_ration + Energy_of_Ration | 0.00143 |

|  | **LCBD – M2** | | | | | | |
| --- | --- | --- | --- | --- | --- | --- | --- |
| *Predictors* | *Estimates* | *std. Error* | *std. Beta* | *CI* | *standardized CI* | *Statistic* | *p* |
| (Intercept) | 0.01989 ^***^ | 0.00218 |  | 0.01555 – 0.02423 |  | 9.12370 | **<0.001** |
| log_CFU_per_g_Campylobacter | 0.00018 ^*^ | 0.00009 | 0.23148 | 0.00001 – 0.00035 | 0.02173 – 0.44123 | 2.16300 | **0.033** |
| Protein_perc_ration | -0.00040 ^***^ | 0.00010 | -0.40572 | -0.00061 – -0.00019 | -0.61548 – -0.19597 | -3.79114 | **<0.001** |
| Observations | 84 | | | | | | |
| R^2^ / R^2^ adjusted | 0.314 / 0.297 | | | | | | |
| ** p<0.05   ** p<0.01   *** p<0.001* | | | | | | | |

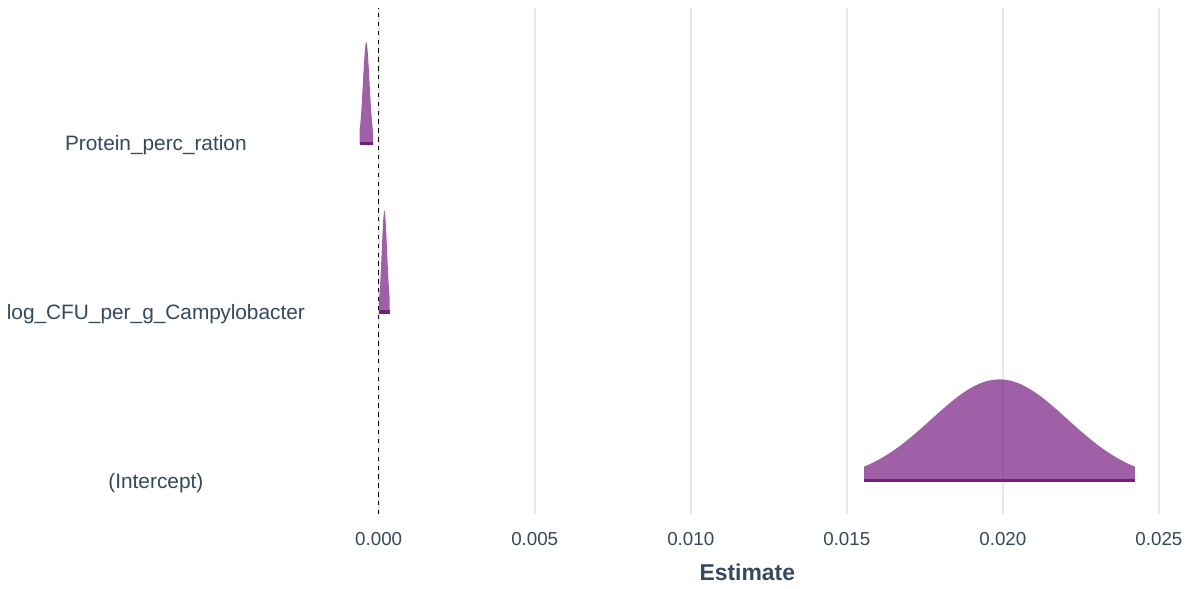

|  | **LCBD – M4** | | | | | | |
| --- | --- | --- | --- | --- | --- | --- | --- |
| *Predictors* | *Estimates* | *std. Error* | *std. Beta* | *CI* | *standardized CI* | *Statistic* | *p* |
| (Intercept) | -0.12247 | 0.19096 |  | -0.50258 – 0.25763 |  | -0.64133 | 0.523 |
| Day_7 | 0.02109 | 0.01187 | 6.14759 | -0.00255 – 0.04472 | -0.63680 – 12.93198 | 1.77600 | 0.080 |
| Total_Mortality_percentage | -0.00050 ^*^ | 0.00022 | -0.29967 | -0.00094 – -0.00006 | -0.55832 – -0.04101 | -2.27070 | **0.026** |
| Protein_perc_ration | -0.00367 ^***^ | 0.00107 | -3.73833 | -0.00580 – -0.00154 | -5.87596 – -1.60070 | -3.42762 | **0.001** |
| Energy_of_Ration | 0.00014 | 0.00014 | 2.93688 | -0.00013 – 0.00041 | -2.66494 – 8.53870 | 1.02756 | 0.307 |
| Observations | 84 | | | | | | |
| R^2^ / R^2^ adjusted | 0.353 / 0.320 | | | | | | |
| ** p<0.05   ** p<0.01   *** p<0.001* | | | | | | | |

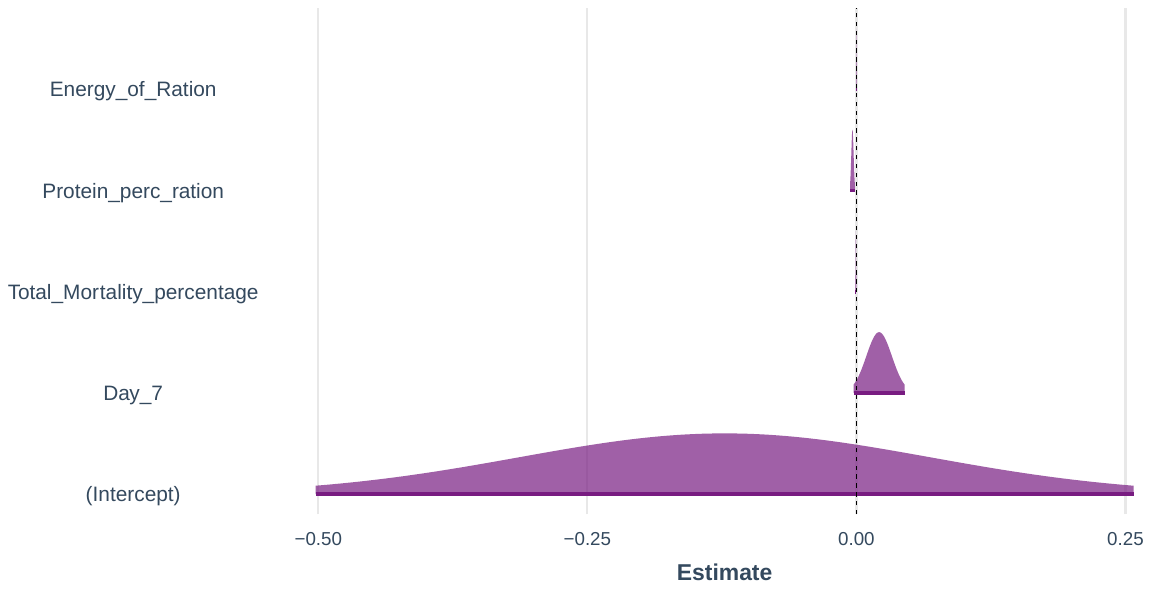

|  | **LCBD – M3** | | | | | | |
| --- | --- | --- | --- | --- | --- | --- | --- |
| *Predictors* | *Estimates* | *std. Error* | *std. Beta* | *CI* | *standardized CI* | *Statistic* | *p* |
| (Intercept) | 0.07288 ^***^ | 0.01799 |  | 0.03707 – 0.10869 |  | 4.05043 | **<0.001** |
| Day_7 | 0.00939 ^**^ | 0.00338 | 2.73756 | 0.00267 – 0.01611 | 0.80764 – 4.66747 | 2.78018 | **0.007** |
| Total_Mortality_percentage | -0.00036 ^*^ | 0.00017 | -0.21397 | -0.00070 – -0.00002 | -0.41450 – -0.01344 | -2.09129 | **0.040** |
| Protein_perc_ration | -0.00320 ^**^ | 0.00097 | -3.25924 | -0.00512 – -0.00127 | -5.19237 – -1.32610 | -3.30447 | **0.001** |
| Observations | 84 | | | | | | |
| R^2^ / R^2^ adjusted | 0.345 / 0.320 | | | | | | |
| ** p<0.05   ** p<0.01   *** p<0.001* | | | | | | | |

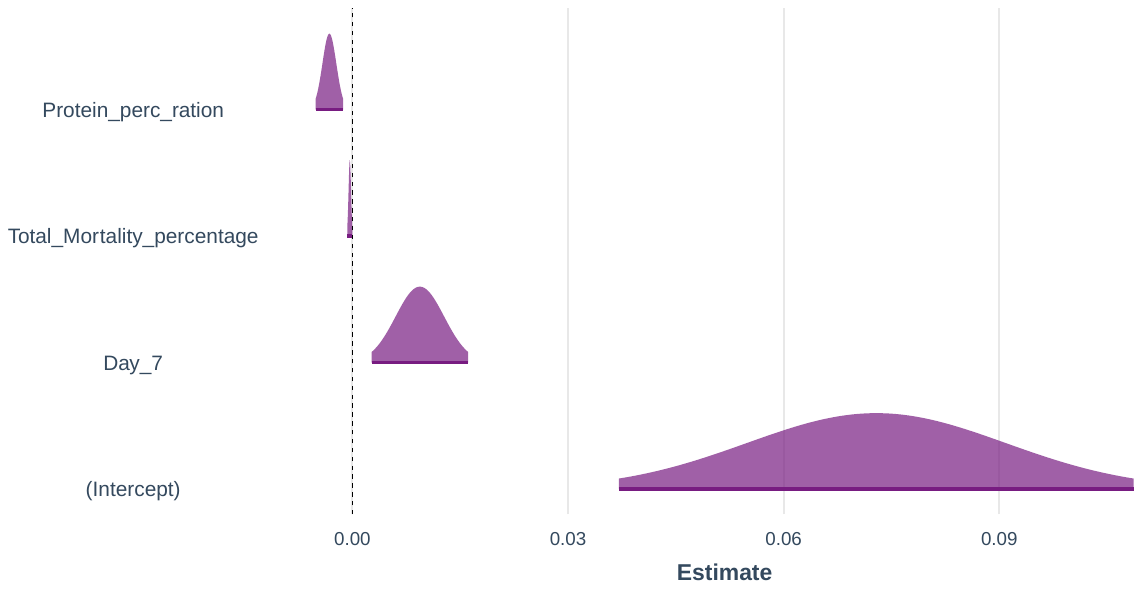

|  | **LCBD – M1** | | | | | | |
| --- | --- | --- | --- | --- | --- | --- | --- |
| *Predictors* | *Estimates* | *std. Error* | *std. Beta* | *CI* | *standardized CI* | *Statistic* | *p* |
| (Intercept) | 0.02239 ^***^ | 0.00189 |  | 0.01863 – 0.02615 |  | 11.85230 | **<0.001** |
| Protein_perc_ration | -0.00051 ^***^ | 0.00009 | -0.52390 | -0.00070 – -0.00033 | -0.70826 – -0.33954 | -5.56962 | **<0.001** |
| Observations | 84 | | | | | | |
| R^2^ / R^2^ adjusted | 0.274 / 0.266 | | | | | | |
| ** p<0.05   ** p<0.01   *** p<0.001* | | | | | | | |

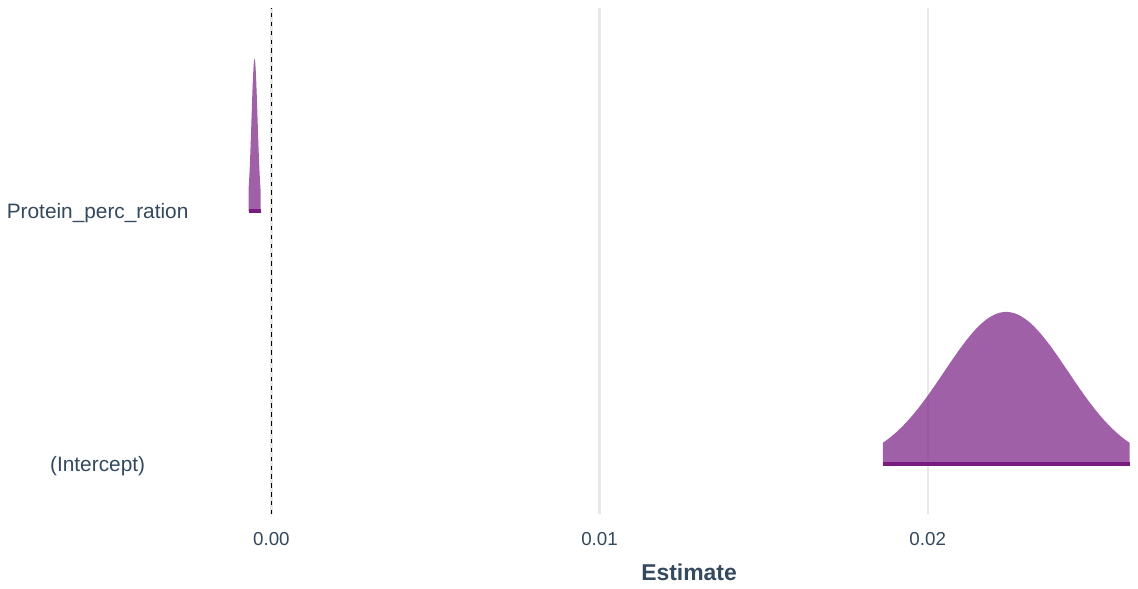

|  | **LCBD – M5** | | | | | | |
| --- | --- | --- | --- | --- | --- | --- | --- |
| *Predictors* | *Estimates* | *std. Error* | *std. Beta* | *CI* | *standardized CI* | *Statistic* | *p* |
| (Intercept) | -0.20926 | 0.21120 |  | -0.62972 – 0.21121 |  | -0.99081 | 0.325 |
| Day_7 | 0.02111 | 0.01192 | 6.15242 | -0.00262 – 0.04483 | -0.65645 – 12.96129 | 1.77100 | 0.080 |
| No_of_Parent_Flocks_Used_3 | -0.00084 | 0.00046 | -0.24020 | -0.00175 – 0.00008 | -0.49901 – 0.01861 | -1.81901 | 0.073 |
| log_CFU_per_g_Campylobacter | 0.00011 | 0.00011 | 0.14152 | -0.00010 – 0.00032 | -0.11735 – 0.40040 | 1.07151 | 0.287 |
| Protein_perc_ration | -0.00267 ^*^ | 0.00110 | -2.71812 | -0.00485 – -0.00048 | -4.90802 – -0.52822 | -2.43272 | **0.017** |
| Energy_of_Ration | 0.00019 | 0.00015 | 3.88650 | -0.00011 – 0.00048 | -2.14034 – 9.91335 | 1.26391 | 0.210 |
| Observations | 84 | | | | | | |
| R^2^ / R^2^ adjusted | 0.358 / 0.317 | | | | | | |
| ** p<0.05   ** p<0.01   *** p<0.001* | | | | | | | |

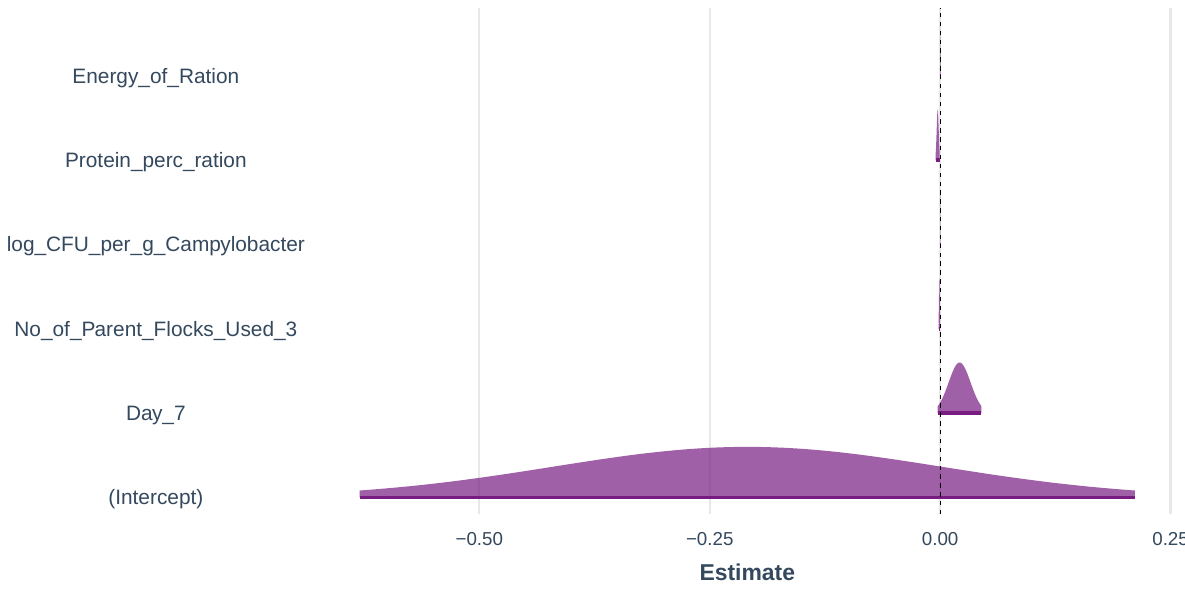

|  | **LCBD – M6** | | | | | | |
| --- | --- | --- | --- | --- | --- | --- | --- |
| *Predictors* | *Estimates* | *std. Error* | *std. Beta* | *CI* | *standardized CI* | *Statistic* | *p* |
| (Intercept) | -0.21670 | 0.23648 |  | -0.68760 – 0.25421 |  | -0.91632 | 0.362 |
| Feed_N | -0.00008 | 0.00112 | -0.02319 | -0.00231 – 0.00214 | -0.65668 – 0.61030 | -0.07175 | 0.943 |
| Feed_HW | -0.00084 | 0.00047 | -0.24107 | -0.00177 – 0.00009 | -0.50262 – 0.02049 | -1.80645 | 0.075 |
| Day_7 | 0.02072 | 0.01315 | 6.03965 | -0.00547 – 0.04690 | -1.47354 – 13.55284 | 1.57556 | 0.119 |
| log_CFU_per_g_Campylobacter | 0.00013 | 0.00029 | 0.16606 | -0.00045 – 0.00071 | -0.55303 – 0.88515 | 0.45262 | 0.652 |
| Protein_perc_ration | -0.00249 | 0.00276 | -2.53303 | -0.00798 – 0.00301 | -8.04839 – 2.98233 | -0.90015 | 0.371 |
| Energy_of_Ration | 0.00019 | 0.00015 | 3.94572 | -0.00012 – 0.00049 | -2.33193 – 10.22338 | 1.23191 | 0.222 |
| Observations | 84 | | | | | | |
| R^2^ / R^2^ adjusted | 0.358 / 0.308 | | | | | | |
| ** p<0.05   ** p<0.01   *** p<0.001* | | | | | | | |

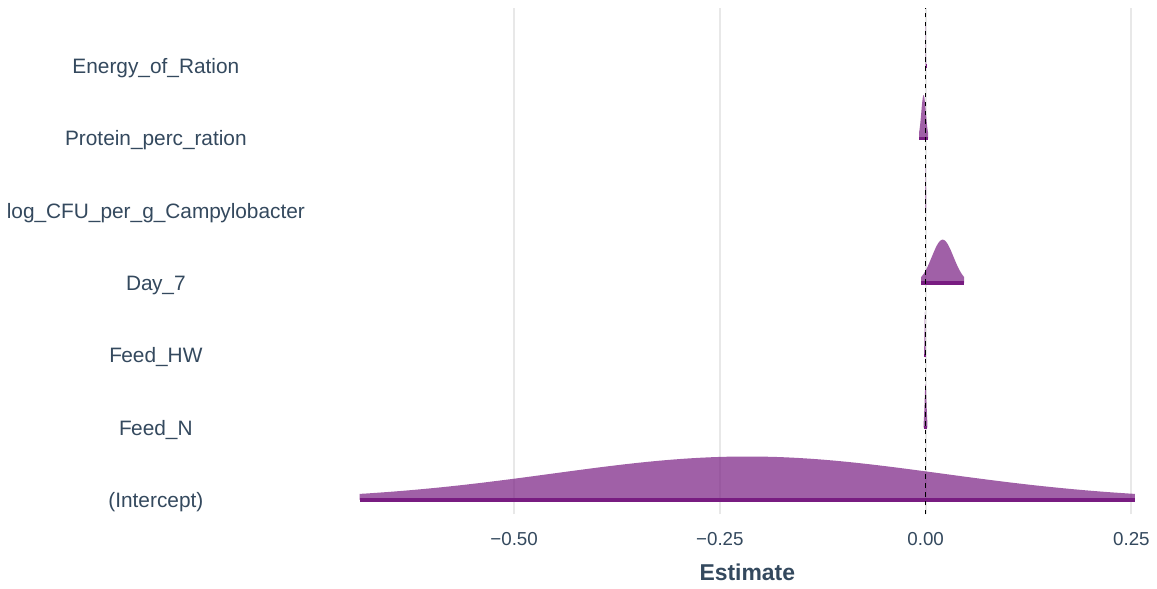
