## Supplementary S9-LCBD Unweighted UniFrac Subset Regression for "Impact of industrial production system parameters on chicken microbiomes: mechanisms to improve performance and reduce *Campylobacter*"

**Top models (Model parameters given afterwards with significant positive influencers highlighted in orange and negative in blue):**

|  | Model | Cross-validation Errors (5-Folds Average RMSE) |
| --- | --- | --- |
| M2 | LCBD ~ Day_30 + Protein_perc_ration | 0.00154 |
| M3 | LCBD ~ Day_30 + Pododermatitis_percentage + Protein_perc_ration | 0.00156 |
| M4 | LCBD ~ Feed_N + Day_7 + Protein_perc_ration + Energy_of_Ration | 0.00158 |
| M1 | LCBD ~ Pododermatitis_percentage | 0.00161 |
| M5 | LCBD ~ Day_30 + Feed_Change_Age_Grower_to_Finisher_20 + Total_Mortality_percentage + Protein_perc_ration + Energy_of_Ration | 0.00162 |
| M6 | LCBD ~ Feed_N + Feed_HW + Day_7 + log_CFU_per_g_Campylobacter + Protein_perc_ration + Energy_of_Ration | 0.00162 |

|  | **LCBD – M2** | | | | | | |
| --- | --- | --- | --- | --- | --- | --- | --- |
| *Predictors* | *Estimates* | *std. Error* | *std. Beta* | *CI* | *standardized CI* | *Statistic* | *p* |
| (Intercept) | -0.08467 ^***^ | 0.02056 |  | -0.12558 – -0.04377 |  | -4.11861 | **<0.001** |
| Day_30 | 0.01534 ^***^ | 0.00332 | 4.38030 | 0.00873 – 0.02195 | 2.52201 – 6.23858 | 4.61997 | **<0.001** |
| Protein_perc_ration | 0.00447 ^***^ | 0.00095 | 4.45725 | 0.00258 – 0.00635 | 2.59896 – 6.31553 | 4.70113 | **<0.001** |
| Observations | 84 | | | | | | |
| R^2^ / R^2^ adjusted | 0.217 / 0.197 | | | | | | |
| ** p<0.05   ** p<0.01   *** p<0.001* | | | | | | | |

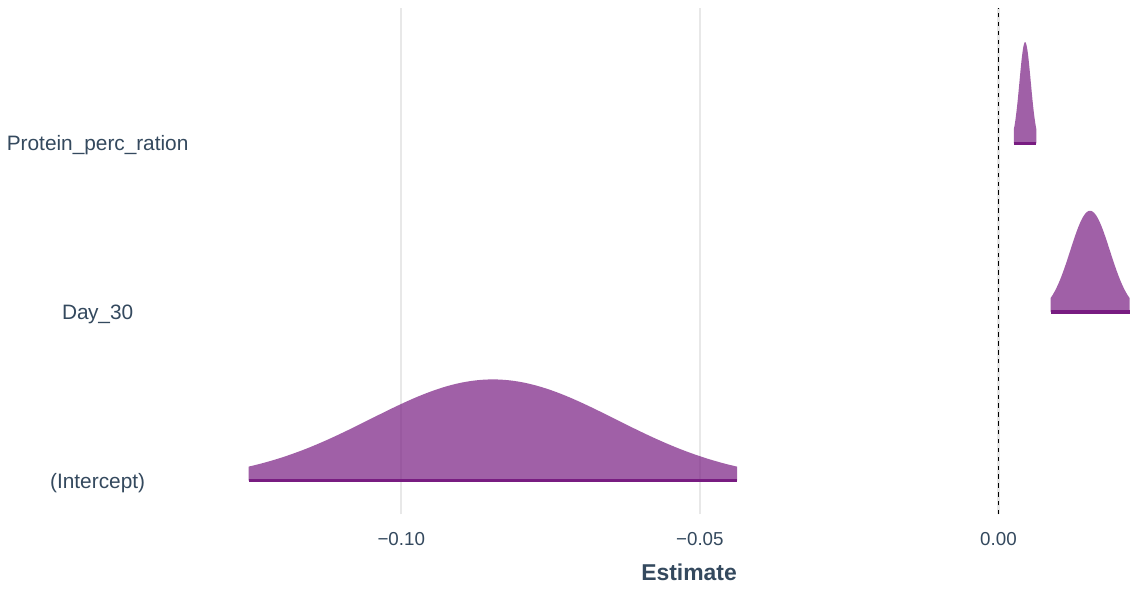

|  | **LCBD – M3** | | | | | | |
| --- | --- | --- | --- | --- | --- | --- | --- |
| *Predictors* | *Estimates* | *std. Error* | *std. Beta* | *CI* | *standardized CI* | *Statistic* | *p* |
| (Intercept) | -0.07194 ^**^ | 0.02313 |  | -0.11797 – -0.02591 |  | -3.11052 | **0.003** |
| Day_30 | 0.01321 ^***^ | 0.00377 | 3.77082 | 0.00571 – 0.02070 | 1.66304 – 5.87859 | 3.50638 | **0.001** |
| Pododermatitis_percentage | 0.00002 | 0.00002 | 0.13303 | -0.00001 – 0.00005 | -0.08602 – 0.35208 | 1.19029 | 0.237 |
| Protein_perc_ration | 0.00385 ^***^ | 0.00108 | 3.84286 | 0.00170 – 0.00600 | 1.73122 – 5.95450 | 3.56683 | **0.001** |
| Observations | 84 | | | | | | |
| R^2^ / R^2^ adjusted | 0.230 / 0.201 | | | | | | |
| ** p<0.05   ** p<0.01   *** p<0.001* | | | | | | | |

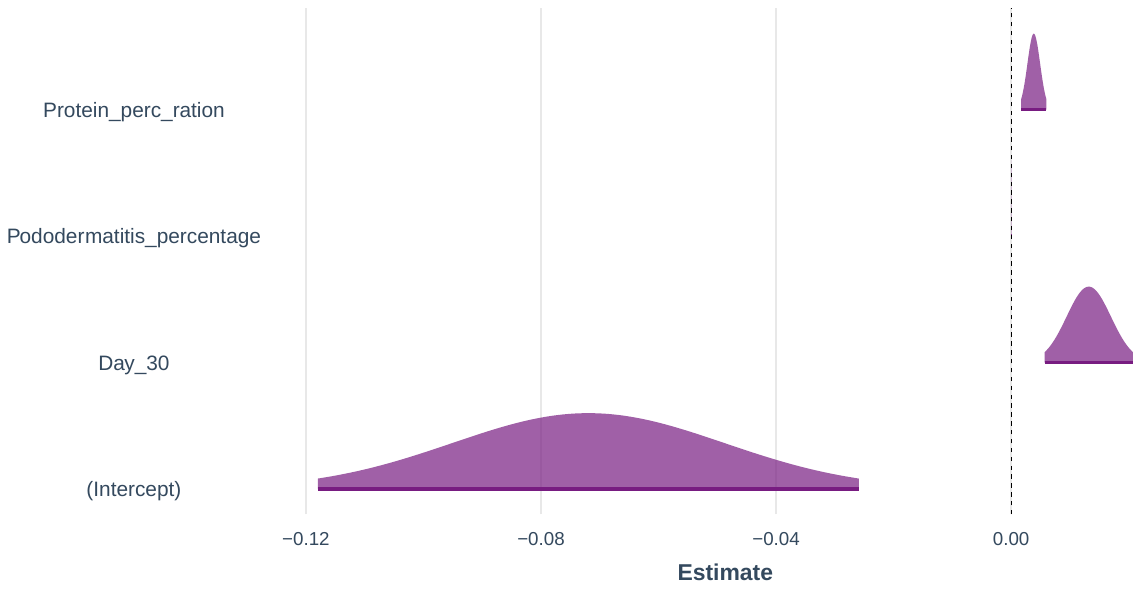

|  | **LCBD – M4** | | | | | | |
| --- | --- | --- | --- | --- | --- | --- | --- |
| *Predictors* | *Estimates* | *std. Error* | *std. Beta* | *CI* | *standardized CI* | *Statistic* | *p* |
| (Intercept) | 0.02980 | 0.18184 |  | -0.33214 – 0.39174 |  | 0.16389 | 0.870 |
| Feed_N | 0.00036 | 0.00043 | 0.10068 | -0.00051 – 0.00122 | -0.13949 – 0.34084 | 0.82161 | 0.414 |
| Day_7 | -0.01834 | 0.01091 | -5.23634 | -0.04006 – 0.00338 | -11.34319 – 0.87051 | -1.68058 | 0.097 |
| Protein_perc_ration | 0.00403 ^***^ | 0.00113 | 4.02538 | 0.00178 – 0.00629 | 1.80714 – 6.24362 | 3.55670 | **0.001** |
| Energy_of_Ration | -0.00006 | 0.00013 | -1.28478 | -0.00032 – 0.00019 | -6.41729 – 3.84773 | -0.49062 | 0.625 |
| Observations | 84 | | | | | | |
| R^2^ / R^2^ adjusted | 0.231 / 0.192 | | | | | | |
| ** p<0.05   ** p<0.01   *** p<0.001* | | | | | | | |

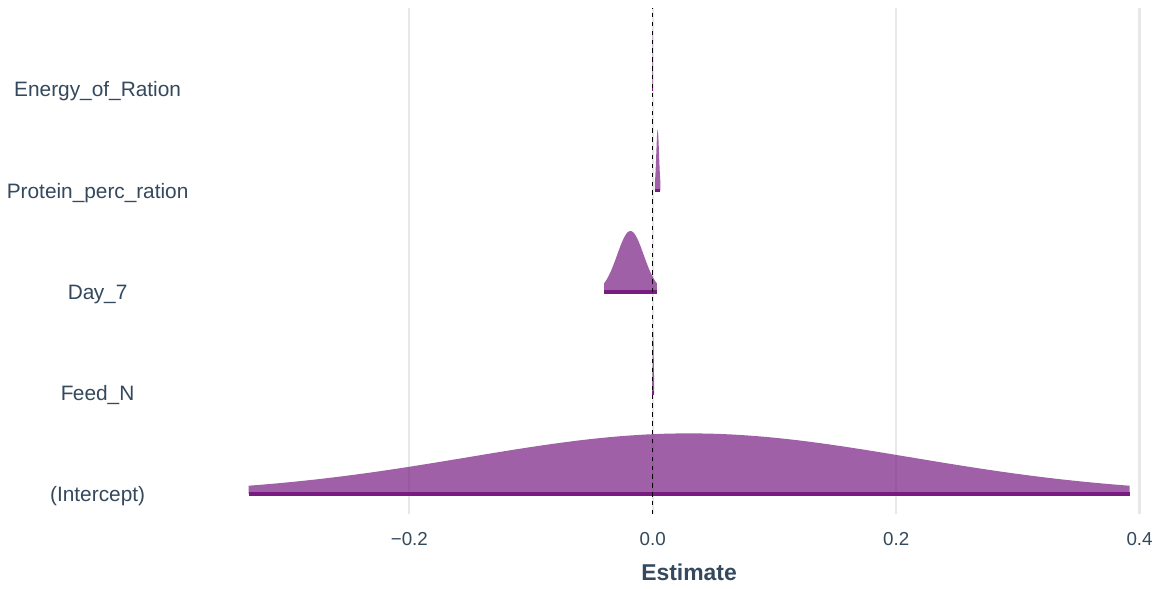

|  | **LCBD – M1** | | | | | | |
| --- | --- | --- | --- | --- | --- | --- | --- |
| *Predictors* | *Estimates* | *std. Error* | *std. Beta* | *CI* | *standardized CI* | *Statistic* | *p* |
| (Intercept) | 0.01047 ^***^ | 0.00049 |  | 0.00949 – 0.01145 |  | 21.22874 | **<0.001** |
| Pododermatitis_percentage | 0.00005 ^**^ | 0.00002 | 0.32461 | 0.00002 – 0.00008 | 0.11988 – 0.52933 | 3.10771 | **0.003** |
| Observations | 84 | | | | | | |
| R^2^ / R^2^ adjusted | 0.105 / 0.094 | | | | | | |
| ** p<0.05   ** p<0.01   *** p<0.001* | | | | | | | |

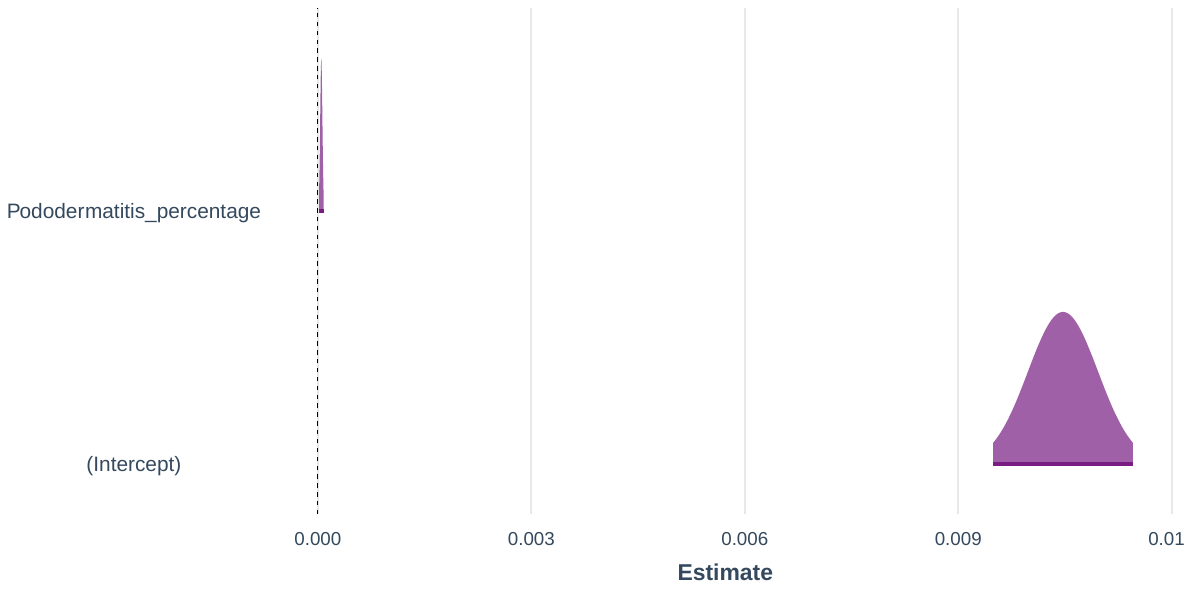

|  | **LCBD – M5** | | | | | | |
| --- | --- | --- | --- | --- | --- | --- | --- |
| *Predictors* | *Estimates* | *std. Error* | *std. Beta* | *CI* | *standardized CI* | *Statistic* | *p* |
| (Intercept) | -0.01404 | 0.21265 |  | -0.43738 – 0.40931 |  | -0.06601 | 0.948 |
| Day_30 | 0.01660 | 0.01369 | 4.73889 | -0.01065 – 0.04385 | -2.91957 – 12.39735 | 1.21278 | 0.229 |
| Feed_Change_Age_Grower_to_Finisher_20 | -0.00012 | 0.00038 | -0.03310 | -0.00088 – 0.00064 | -0.24085 – 0.17464 | -0.31232 | 0.756 |
| Total_Mortality_percentage | -0.00018 | 0.00025 | -0.10836 | -0.00068 – 0.00031 | -0.39405 – 0.17732 | -0.74344 | 0.459 |
| Protein_perc_ration | 0.00395 ^**^ | 0.00120 | 3.94574 | 0.00156 – 0.00634 | 1.59699 – 6.29448 | 3.29261 | **0.001** |
| Energy_of_Ration | -0.00004 | 0.00016 | -0.86632 | -0.00036 – 0.00028 | -7.30494 – 5.57230 | -0.26371 | 0.793 |
| Observations | 84 | | | | | | |
| R^2^ / R^2^ adjusted | 0.231 / 0.182 | | | | | | |
| ** p<0.05   ** p<0.01   *** p<0.001* | | | | | | | |

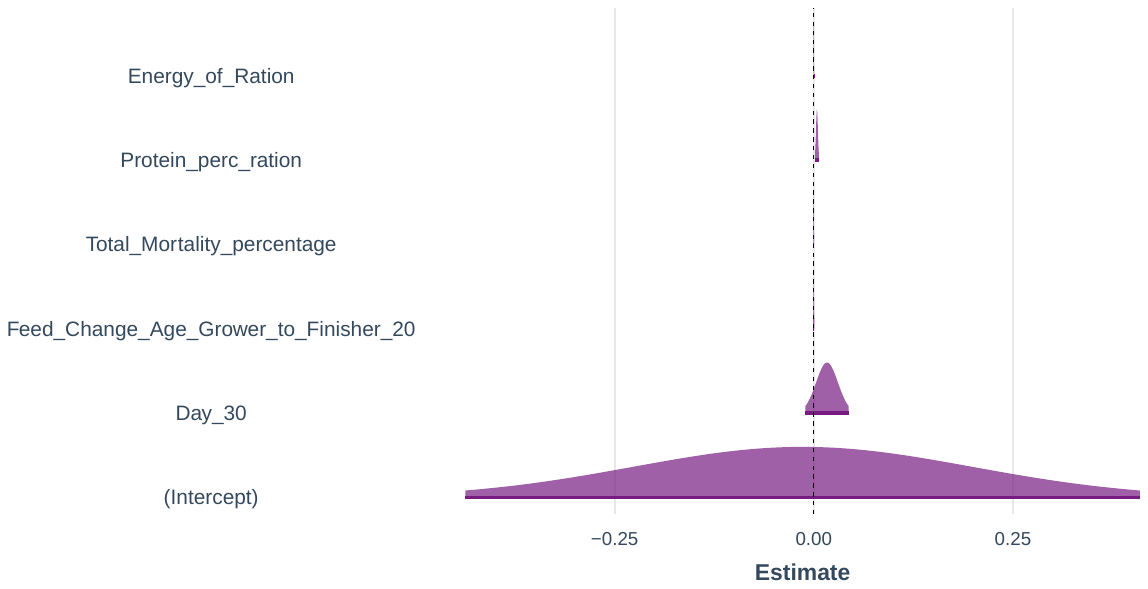

|  | **LCBD – M6** | | | | | | |
| --- | --- | --- | --- | --- | --- | --- | --- |
| *Predictors* | *Estimates* | *std. Error* | *std. Beta* | *CI* | *standardized CI* | *Statistic* | *p* |
| (Intercept) | -0.00759 | 0.26420 |  | -0.53368 – 0.51849 |  | -0.02875 | 0.977 |
| Feed_N | 0.00025 | 0.00125 | 0.07154 | -0.00223 – 0.00274 | -0.62161 – 0.76470 | 0.20229 | 0.840 |
| Feed_HW | -0.00011 | 0.00052 | -0.03094 | -0.00115 – 0.00092 | -0.31712 – 0.25525 | -0.21189 | 0.833 |
| Day_7 | -0.01694 | 0.01469 | -4.83780 | -0.04620 – 0.01231 | -13.05857 – 3.38298 | -1.15341 | 0.252 |
| log_CFU_per_g_Campylobacter | 0.00002 | 0.00033 | 0.02722 | -0.00063 – 0.00067 | -0.75959 – 0.81404 | 0.06782 | 0.946 |
| Protein_perc_ration | 0.00415 | 0.00308 | 4.13786 | -0.00200 – 0.01029 | -1.89693 – 10.17266 | 1.34388 | 0.183 |
| Energy_of_Ration | -0.00004 | 0.00017 | -0.78749 | -0.00038 – 0.00030 | -7.65637 – 6.08139 | -0.22470 | 0.823 |
| Observations | 84 | | | | | | |
| R^2^ / R^2^ adjusted | 0.231 / 0.171 | | | | | | |
| ** p<0.05   ** p<0.01   *** p<0.001* | | | | | | | |

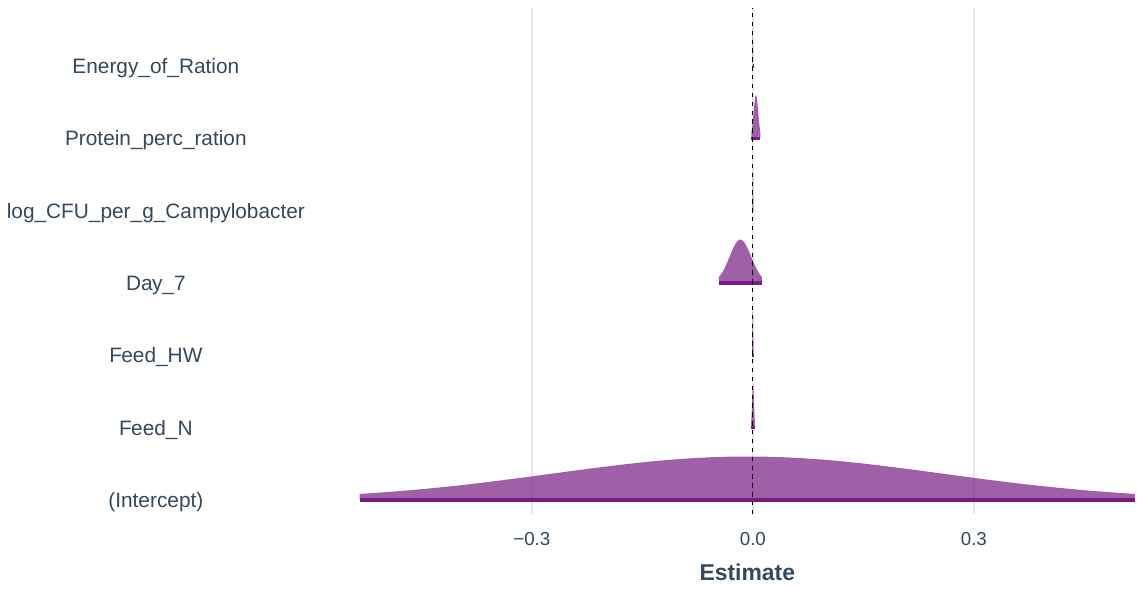
