## Supplementary S10-LCBD Weighted UniFrac Subset Regression for "Impact of industrial production system parameters on chicken microbiomes: mechanisms to improve performance and reduce *Campylobacter*"

**Top models (Model parameters given afterwards with significant positive influencers highlighted in orange and negative in blue):**

|  | Model | Cross-validation Errors (5-Folds Average RMSE) |
| --- | --- | --- |
| M1 | LCBD ~ Protein_perc_ration | 0.00258 |
| M3 | LCBD ~ Day_7 + Protein_perc_ration + Energy_of_Ration | 0.00263 |
| M4 | LCBD ~ Day_30 + Total_Mortality_percentage + Protein_perc_ration + Energy_of_Ration | 0.00263 |
| M2 | LCBD ~ Day_30 + Protein_perc_ration | 0.00265 |
| M5 | LCBD ~ Day_30 + EPEF + log_CFU_per_g_Campylobacter + Protein_perc_ration + Energy_of_Ration | 0.00271 |
| M6 | LCBD ~ Day_30 + No_of_Parent_Flocks_Used_3 + Feed_Change_Age_Finisher_to_Withdrawal_34 + log_CFU_per_g_Campylobacter + Protein_perc_ration + Energy_of_Ration | 0.00272 |

|  | **LCBD – M1** | | | | | | |
| --- | --- | --- | --- | --- | --- | --- | --- |
| *Predictors* | *Estimates* | *std. Error* | *std. Beta* | *CI* | *standardized CI* | *Statistic* | *p* |
| (Intercept) | 0.02942 ^***^ | 0.00349 |  | 0.02247 – 0.03637 |  | 8.42387 | **<0.001** |
| Protein_perc_ration | -0.00086 ^***^ | 0.00017 | -0.48572 | -0.00120 – -0.00052 | -0.67492 – -0.29653 | -5.03186 | **<0.001** |
| Observations | 84 | | | | | | |
| R^2^ / R^2^ adjusted | 0.236 / 0.227 | | | | | | |
| ** p<0.05   ** p<0.01   *** p<0.001* | | | | | | | |

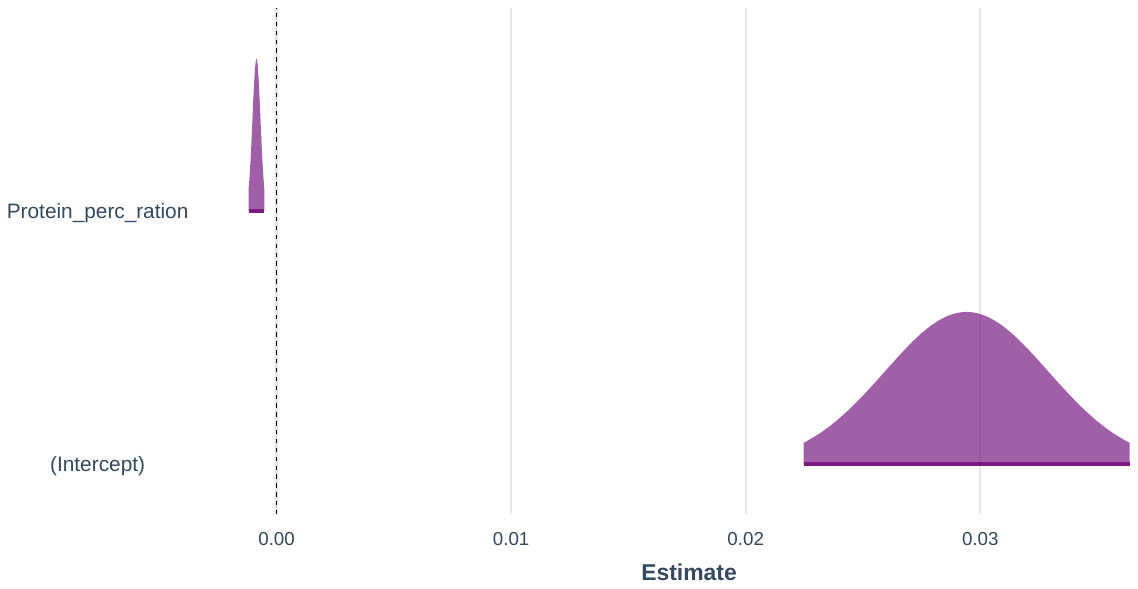

|  | **LCBD – M3** | | | | | | |
| --- | --- | --- | --- | --- | --- | --- | --- |
| *Predictors* | *Estimates* | *std. Error* | *std. Beta* | *CI* | *standardized CI* | *Statistic* | *p* |
| (Intercept) | -0.24349 | 0.29632 |  | -0.83319 – 0.34620 |  | -0.82172 | 0.414 |
| Day_7 | 0.01964 | 0.01615 | 3.17893 | -0.01250 – 0.05179 | -1.94468 – 8.30254 | 1.21606 | 0.228 |
| Protein_perc_ration | -0.00227 | 0.00165 | -1.28435 | -0.00556 – 0.00102 | -3.11863 – 0.54993 | -1.37235 | 0.174 |
| Energy_of_Ration | 0.00020 | 0.00020 | 2.37880 | -0.00020 – 0.00061 | -2.26355 – 7.02115 | 1.00431 | 0.318 |
| Observations | 84 | | | | | | |
| R^2^ / R^2^ adjusted | 0.251 / 0.223 | | | | | | |
| ** p<0.05   ** p<0.01   *** p<0.001* | | | | | | | |

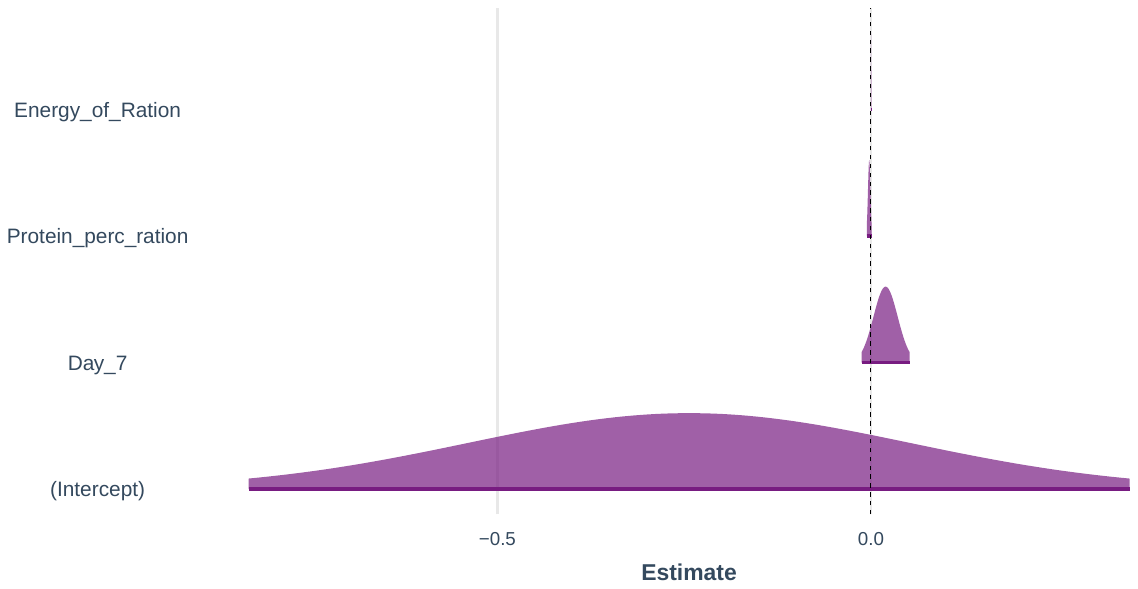

|  | **LCBD – M4** | | | | | | |
| --- | --- | --- | --- | --- | --- | --- | --- |
| *Predictors* | *Estimates* | *std. Error* | *std. Beta* | *CI* | *standardized CI* | *Statistic* | *p* |
| (Intercept) | -0.43676 | 0.34646 |  | -1.12636 – 0.25284 |  | -1.26065 | 0.211 |
| Day_30 | -0.03673 | 0.02286 | -5.94469 | -0.08223 – 0.00876 | -13.19406 – 1.30469 | -1.60722 | 0.112 |
| Total_Mortality_percentage | -0.00045 | 0.00042 | -0.14894 | -0.00129 – 0.00040 | -0.42533 – 0.12744 | -1.05622 | 0.294 |
| Protein_perc_ration | -0.00357 | 0.00206 | -2.01883 | -0.00767 – 0.00053 | -4.30297 – 0.26531 | -1.73230 | 0.087 |
| Energy_of_Ration | 0.00038 | 0.00026 | 4.41729 | -0.00014 – 0.00090 | -1.56846 – 10.40304 | 1.44639 | 0.152 |
| Observations | 84 | | | | | | |
| R^2^ / R^2^ adjusted | 0.261 / 0.224 | | | | | | |
| ** p<0.05   ** p<0.01   *** p<0.001* | | | | | | | |

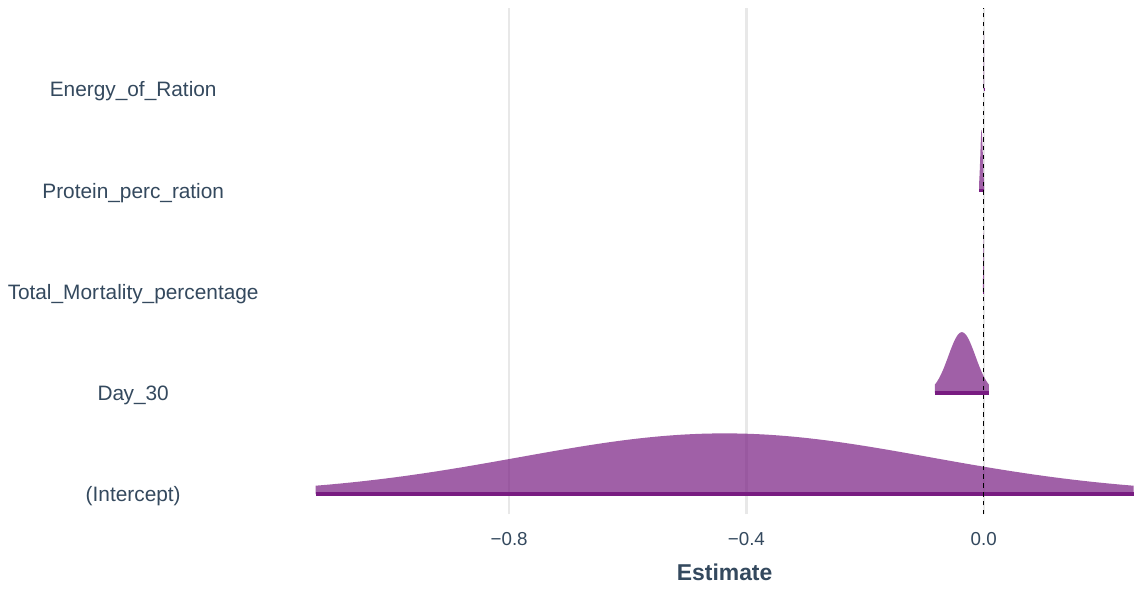

|  | **LCBD – M2** | | | | | | |
| --- | --- | --- | --- | --- | --- | --- | --- |
| *Predictors* | *Estimates* | *std. Error* | *std. Beta* | *CI* | *standardized CI* | *Statistic* | *p* |
| (Intercept) | 0.05707 | 0.03568 |  | -0.01393 – 0.12807 |  | 1.59929 | 0.114 |
| Day_30 | -0.00449 | 0.00576 | -0.72636 | -0.01596 – 0.00698 | -2.55470 – 1.10198 | -0.77865 | 0.438 |
| Protein_perc_ration | -0.00214 | 0.00165 | -1.20816 | -0.00542 – 0.00115 | -3.03651 – 0.62018 | -1.29514 | 0.199 |
| Observations | 84 | | | | | | |
| R^2^ / R^2^ adjusted | 0.242 / 0.223 | | | | | | |
| ** p<0.05   ** p<0.01   *** p<0.001* | | | | | | | |

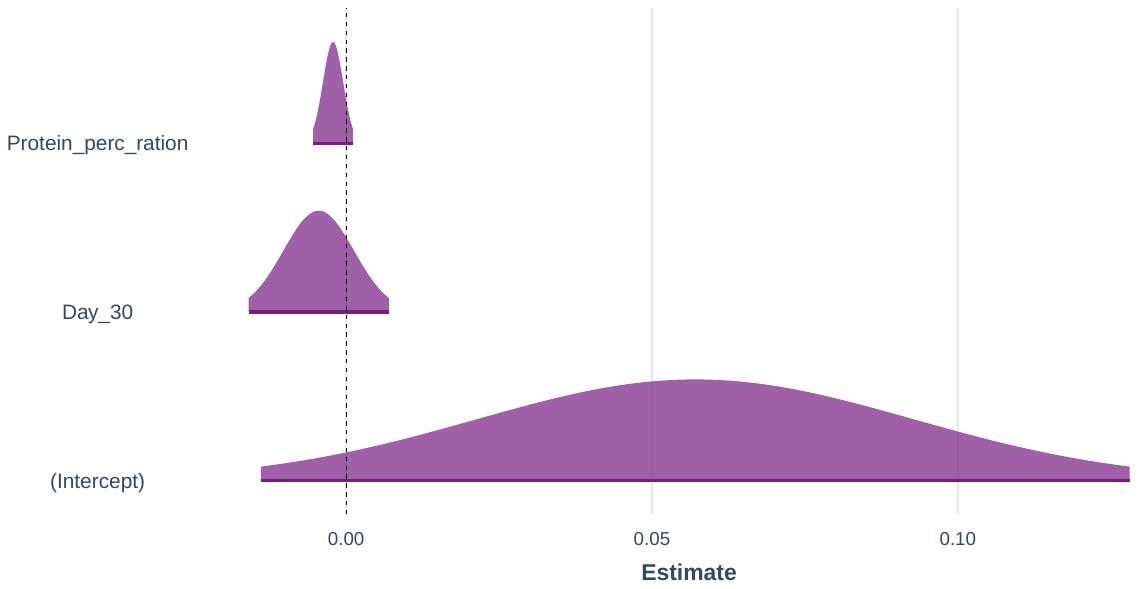

|  | **LCBD – M5** | | | | | | |
| --- | --- | --- | --- | --- | --- | --- | --- |
| *Predictors* | *Estimates* | *std. Error* | *std. Beta* | *CI* | *standardized CI* | *Statistic* | *p* |
| (Intercept) | -0.42641 | 0.34899 |  | -1.12120 – 0.26837 |  | -1.22185 | 0.225 |
| Day_30 | -0.03906 | 0.02501 | -6.32138 | -0.08886 – 0.01074 | -14.25550 – 1.61274 | -1.56157 | 0.122 |
| EPEF | 0.00002 | 0.00003 | 0.19768 | -0.00003 – 0.00008 | -0.25322 – 0.64858 | 0.85927 | 0.393 |
| log_CFU_per_g_Campylobacter | -0.00009 | 0.00032 | -0.06474 | -0.00074 – 0.00055 | -0.50675 – 0.37727 | -0.28707 | 0.775 |
| Protein_perc_ration | -0.00436 | 0.00348 | -2.46582 | -0.01129 – 0.00258 | -6.32965 – 1.39800 | -1.25082 | 0.215 |
| Energy_of_Ration | 0.00038 | 0.00026 | 4.38100 | -0.00014 – 0.00090 | -1.56849 – 10.33049 | 1.44325 | 0.153 |
| Observations | 84 | | | | | | |
| R^2^ / R^2^ adjusted | 0.262 / 0.215 | | | | | | |
| ** p<0.05   ** p<0.01   *** p<0.001* | | | | | | | |

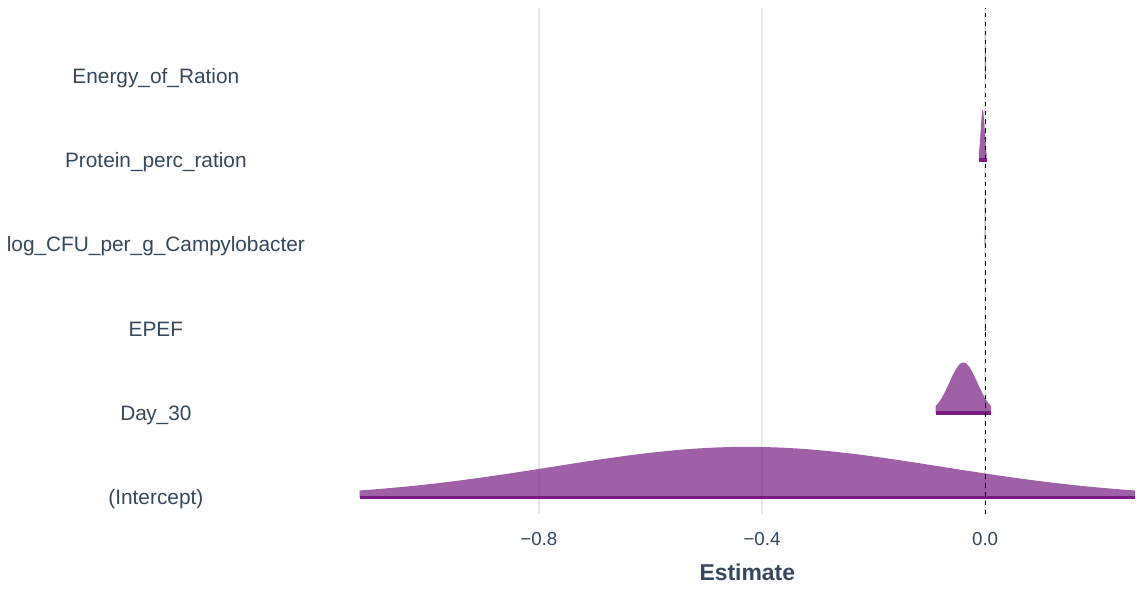

|  | **LCBD – M6** | | | | | | |
| --- | --- | --- | --- | --- | --- | --- | --- |
| *Predictors* | *Estimates* | *std. Error* | *std. Beta* | *CI* | *standardized CI* | *Statistic* | *p* |
| (Intercept) | -0.42001 | 0.44255 |  | -1.30124 – 0.46121 |  | -0.94908 | 0.346 |
| Day_30 | -0.03903 | 0.02539 | -6.31664 | -0.08959 – 0.01153 | -14.37068 – 1.73740 | -1.53716 | 0.128 |
| No_of_Parent_Flocks_Used_3 | -0.00067 | 0.00090 | -0.10719 | -0.00246 – 0.00112 | -0.38757 – 0.17319 | -0.74930 | 0.456 |
| Feed_Change_Age_Finisher_to_Withdrawal_34 | 0.00073 | 0.00216 | 0.11773 | -0.00356 – 0.00503 | -0.56137 – 0.79683 | 0.33979 | 0.735 |
| log_CFU_per_g_Campylobacter | -0.00009 | 0.00056 | -0.06189 | -0.00121 – 0.00103 | -0.83275 – 0.70896 | -0.15737 | 0.875 |
| Protein_perc_ration | -0.00432 | 0.00533 | -2.44569 | -0.01494 – 0.00629 | -8.35809 – 3.46671 | -0.81075 | 0.420 |
| Energy_of_Ration | 0.00038 | 0.00030 | 4.39488 | -0.00021 – 0.00097 | -2.33468 – 11.12445 | 1.28000 | 0.204 |
| Observations | 84 | | | | | | |
| R^2^ / R^2^ adjusted | 0.262 / 0.204 | | | | | | |
| ** p<0.05   ** p<0.01   *** p<0.001* | | | | | | | |

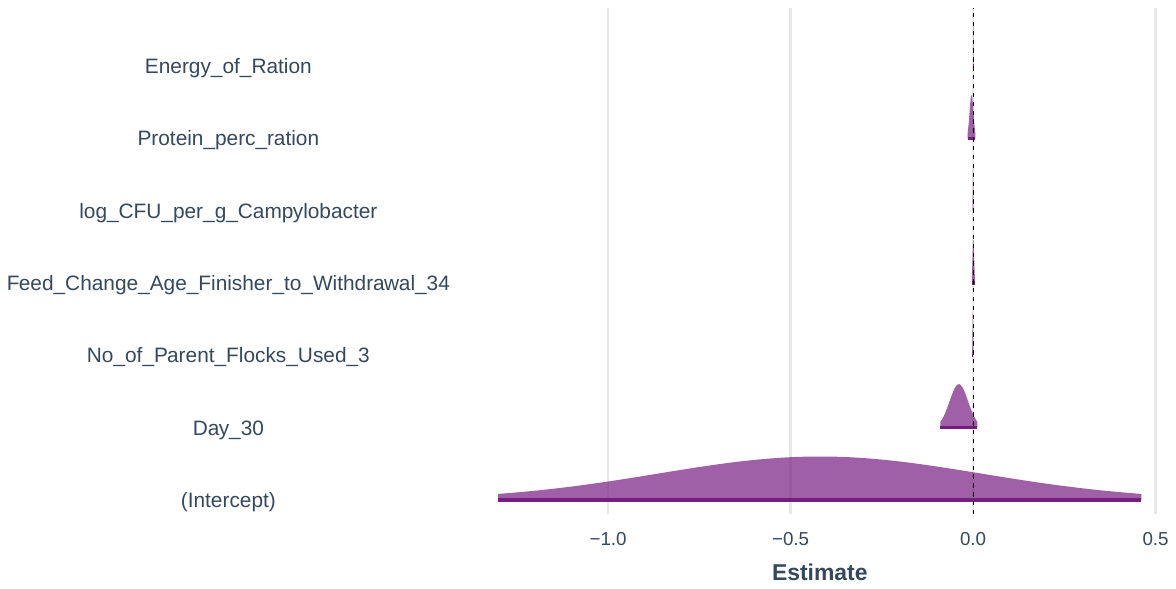
