## Supplementary S12-Bioinformatics_and_StatisticalAnalysis for "Impact of industrial production system parameters on chicken microbiomes: mechanisms to improve performance and reduce *Campylobacter*"

The software VSEARCH v2.3.4 (steps documented in <http://github.com/torognes/vsearch/wiki/VSEARCH-pipeline>) was used to generate the abundance table by constructing operational taxonomic units (OTUs), a proxy for species. Before using VSEARCH, the paired-end reads were pre-processed according to the recommendations given in author’s recent publications (Schirmer et al., 2015;D'Amore et al., 2016). Briefly, we quality trimmed (average Phred quality score of 20 using a sliding window approach) and filtered the reads using Sickle v1.200 (Joshi and Fass, 2011). Next, BayesHammer (Nikolenko et al., 2013) was used from the Spades v2.5.0 assembler, which error-corrected the paired-end reads. Following this, pandaseqv(2.4) (Masella et al., 2012) was used to assemble the forward and reverse reads into a single sequence spanning the entire V4 region with a minimum overlap of 10 bp. The reads were then pooled together, dereplicated, sorted in order of decreasing abundance and singletons were discarded. Next, the reads were clustered based on 97% similarity followed by a removal of clusters which had chimeric models built from more abundant reads (--uchime_denovo option in vsearch). In the next step, we employed a reference-based chimera filtering step (--uchime_ref option in vsearch) using a gold database (<https://www.mothur.org/w/images/f/f1/Silva.gold.bacteria.zip>). Finally, the OTU table was generated by matching the original barcoded reads against clean OTUs (a total of 3,815 OTUs for n=86 samples) at 97% similarity (a proxy for species-level separation) with summary read statistics for samples as follows: [1^st^ Quantile: 102,454, Median: 144,736, Mean: 190,358, 3^rd^ Quantile: 267,175, Max: 1,589,970]. We then used the assign_taxonomy.py script from the Qiime workflow (Caporaso et al., 2010) to taxonomically classify the representative OTUs against the SILVA SSU Ref NR database release v123 database. After, the OTUs were multisequence aligned using mafft v7.3 (Katoh and Standley, 2013) and subsequently, aligned sequences were used in FastTree v2.1.7 (Price et al., 2010) to generate the phylogenetic tree in NEWICK format. The biom file for the OTUs was then generated by combining the abundance table with taxonomy information using make_otu_table.py from the Qiime workflow.

**Statistical Analyses**

Statistical analyses were performed in R using the combined data generated from the bioinformatics as well as metadata associated with the study by relying on the vegan package (Oksanen et al., 2015) for alpha and beta diversity analyses. In the latter case, we have used the distance metrics: *Bray-Curtis* is a distance metric which considers only OTU abundance counts; *Unweighted Unifrac* is a phylogenetic distance metric which calculates the distance between samples by taking the proportion of the sum of unshared branch lengths in the sum of all the branch lengths of the phylogenetic tree for the OTUs observed in two samples, and without taking into account their abundances and; *Weighted Unifrac* is a phylogenetic distance metric combining phylogenetic distance with relative abundances. This places emphasis on dominant OTUs or taxa. Unifrac distances were calculated using the phyloseq package (McMurdie and Holmes, 2013).

Analysis of variance for explanatory variables (or sources of variation) was performed using Vegan’s adonis() against distance matrices (Bray-Curtis/UnweightedUniFrac/Weighted UniFrac). This function, referred to as PERMANOVA, fit linear models to distance matrices and used a permutation test with pseudo-F ratios to give sources of variation. We have used *Treatments* (N, HW, and O) and *Days* in beta diversity plots, and other recorded parameters (**Poultry Growth and Performance Measurements**): *Days*; *Weight Gain / Day (kg)*; *Food Conversion Ratio*; *EPEF*; *Total Mortality (%)*; *Total Leg Culls (%)*; *Pododermatitis (%)*; *Hockmark (%)*; *PMI Rejects (%)*; *Avg. Weight at Slaughter (kg)*; *Age at Thin*; *Age Total Depopulation*; *Total Water Consumption (l)*; *Water Consumption (l) / Bird*; log CFU/g *Campylobacter*; *Protein (% of Ration)*; and *Energy of Ration (Kcal/lb AME)*. These additional parameters were used in Table 2. R^2^ (if significant) returned by PERMANOVA then explained the percentage variability by these predictors.

To give an account of environmental filtering (phylogenetic overdispersion versus clustering), phylogenetic distances within each sample were further characterised by calculating the nearest taxa index (NTI) and net relatedness index (NRI). This analysis helps determine whether the community structure was stochastic (overdispersion and driven by competition among taxa) or deterministic (clustering and driven by strong environmental pressure). The NTI was calculated using mntd() and ses.mntd(), and the mean phylogenetic diversity (MPD) and NRI were calculated using mpd() and ses.mpd() functions from the picante package (Kembel et al., 2010). NTI and NRI represent the negatives of the output from ses.mntd() and ses.mpd(), respectively. Additionally, they quantify the number of standard deviations that separate the observed values from the mean of the null distribution (999 randomisation using null.model-‘richness’ in the ses.mntd() and ses.mpd() functions and only considering taxa as either present or absent regardless of their relative abundance). As opposed to authors’ recent work (Ijaz et al., 2018) OTUs collated at genera were used for the calculations.

Discriminant analyses were performed in two different settings. Since we have a nested design, i.e., microbial community data from production systems, N, O, and HW for both day 7 and 30, we used the Multivariate Integration (MINT) algorithm (Rohart et al., 2017) to compare the sampling time (Days 7 and 30). The algorithm is an extension of the multi-group Projection to Latent Structure (mgPLS), and it attempts to find a common projection space across all studies (three categories as mentioned-above), defined on a small subset of discriminative variables that consistently discriminate the outcome classes (Days 7 and 30). In MINT, we have combined $M=3$ datasets denoted $X^{\left( 1 \right)}\left( N_{1}\times P \right)$, $X^{\left( 2 \right)}\left( N_{2}\times P \right)$, …,$X^{\left( 3 \right)}\left( N_{3}\times P \right)$ , where all these datasets share the P genera whilst the number of samples differ, i.e., $N_{1}$, $N_{2}$,…,$N_{3}$. All studies have associated dummy indicator outcome $Y^{\left( 1 \right)}$, $Y^{\left( 2 \right)}$,…,$Y^{\left( 3 \right)}$ in which all the sampling times (Days 7 and 30) are represented. MINT then solves the problem: ${\text{max} \atop a_{h}, b_{h}}\sum_{m=1}^{M} N_{m}\text{cov}(X_{h}^{(m)}a_{h},Y_{h}^{(m)}b_{h})$, with the constraints$\left\| a_{h} \right\|_{2}=1$ and $\left\| a_{h} \right\|_{1}\leq\lambda$, where the covariance of scores between the datasets are maximised by finding the global loading vectors $a_{h}$ and $b_{h}$ common to all studies (akin to PCA analysis). The first constraint ensures the loading vector to have unit magnitude (requirement of the procedure) and the second constraint (also called $l_{1}$ penalty) to ensure that for the features that do not vary between the categories, the corresponding loading vector coefficients go to zero. This is done by using the sparsity control parameter $\lambda$ in the above equation, and by adjusting it enforces shrinkage of loading vector coefficients. According to the recommendations given in mixOmics package (http://www.mixomics.org), before applying the procedure splsda(), we pre-filter 1% of the lowest abundant genera and then perform TSS+CLR (Total Sum Scaling followed by Centralised Log Ratio) normalisation. To predict the number of latent components (associated loading vectors) and the number of discriminants, the perf.plsda() and tune.splsda() functions were used, respectively. In the latter case, we fine tune the model was applied using leave-one-out cross-validation by splitting the data into training and testing sets and then finding the classification error rates employing overall error rates, between the predicted latent variables with the centroid of the class labels (categories considered in this study) using the centroid distance

To find genera that are significantly different between different categories, we used DESeqDataSetFromMatrix() function from DESeq2 (Love et al., 2014) package with the adjusted p-value significance cut-off of 0.05 and log2 fold change cut-off of 2. This function uses negative binomial GLM to obtain maximum likelihood estimates for OTUs log fold change between two conditions. Then Bayesian shrinkage is applied to obtain shrunken log fold changes subsequently employing the Wald test for obtaining significances. While MINT gave the discriminants shared in a nested model on a global scale), DESeq2 identified changes on a local scale (in conjunction with beta diversity analysis) to identify genera that are causing the shift in microbial communities. Furthermore, we used R’s Metacoder package to generate differential heat trees (Foster et al., 2017) visualise differentially expressed lineages (using Wilcoxin p-value test adjusted with multiple comparison) comparing different categories.

The “BVSTEP” routine (Clarke and Ainsworth, 1993) was used to search for the highest correlation, in a Mantel test, by imploding the abundance table at genera level to absolute minimal set of genera that preserve the beta diversity between samples. To run this algorithm, bvStep() (from the sinkr package) (Taylor, 2014) was used as considered in author’s recent publication (Ijaz et al., 2018).

We performed subset regression against different microbiome metrics (Supplementary S4 to S10) by testing all possible combination of the explanatory variables, and then selecting the best model according to some statistical criteria, with recommendations given in (Kassambara, 2018) and code available at <http://www.sthda.com/english/articles/37-model-selection-essentials-in-r/155-best-subsets-regression-essentials-in-r/>. The R function regubsets() from leaps (Lumley and Miller, 2009) package was used to identify different best models of different sizes, by specifying the option nvmax, set to the maximum number of predictors to incorporate the model. Having obtained the best possible subsets, the k-fold cross-validation consisting of first dividing the data into k subsets. Each subset (10%) served successively as test data set and the remaining subset (90%) as training data. The average cross-validation error is then computed as the model prediction error. This was all done using a custom function utilising R’s train() function from the caret package (Kuhn, 2008). Finally R’s tab_model() function from sjPlot package (Lüdecke, 2018) was used to obtain the statistics for each model. To find the core microbiome, we have used R’s microbiome package (Lahti et al., 2017) and the recommendations given in (Shetty et al., 2017).

In majority of the figures displaying boxplots, pair-wise ANOVA was performed taking two categories at a time, and where significant (p ≤ 0.05), joined them together by a line and plotting significance on top (*: 0.01 ≤ p < 0.05; **: 0.05 ≤ p < 0.001; ***: p ≤ 0.001).

Caporaso, J.G., Kuczynski, J., Stombaugh, J., Bittinger, K., Bushman, F.D., Costello, E.K., Fierer, N., Pena, A.G., Goodrich, J.K., Gordon, J.I., Huttley, G.A., Kelley, S.T., Knights, D., Koenig, J.E., Ley, R.E., Lozupone, C.A., Mcdonald, D., Muegge, B.D., Pirrung, M., Reeder, J., Sevinsky, J.R., Turnbaugh, P.J., Walters, W.A., Widmann, J., Yatsunenko, T., Zaneveld, J., and Knight, R. (2010). QIIME allows analysis of high-throughput community sequencing data. *Nat Methods* 7**,** 335-336. doi: 10.1038/nmeth.f.303.

Clarke, K.R., and Ainsworth, M. (1993). A Method of Linking Multivariate Community Structure to Environmental Variables. *Marine Ecology Progress Series* 92**,** 205-219.

D'amore, R., Ijaz, U.Z., Schirmer, M., Kenny, J.G., Gregory, R., Darby, A.C., Shakya, M., Podar, M., Quince, C., and Hall, N. (2016). A comprehensive benchmarking study of protocols and sequencing platforms for 16S rRNA community profiling. *BMC Genomics* 17**,** 55. doi: 10.1186/s12864-015-2194-9.

Foster, Z.S., Sharpton, T.J., and Grünwald, N.J. (2017). Metacoder: an R package for visualization and manipulation of community taxonomic diversity data. *PLoS computational biology* 13**,** e1005404.

Ijaz, U.Z., Sivaloganathan, L., Mckenna, A., Richmond, A., Kelly, C., Linton, M., Stratakos, A.C., Lavery, U., Elmi, A., Wren, B.W., Dorrell, N., Corcionivoschi, N., and Gundogdu, O. (2018). Comprehensive Longitudinal Microbiome Analysis of the Chicken Cecum Reveals a Shift From Competitive to Environmental Drivers and a Window of Opportunity for Campylobacter. *Front Microbiol* 9**,** 2452. doi: 10.3389/fmicb.2018.02452.

Joshi, N., and Fass, J. (2011). "Sickle: A sliding-window, adaptive, quality-based trimming tool for FastQ files (Version 1.33)". 1.33 ed.).

Kassambara, A. (2018). *Machine Learning Essentials: Practical Guide in R.* sthda.

Katoh, K., and Standley, D.M. (2013). MAFFT multiple sequence alignment software version 7: improvements in performance and usability. *Mol Biol Evol* 30**,** 772-780. doi: 10.1093/molbev/mst010.

Kembel, S.W., Cowan, P.D., Helmus, M.R., Cornwell, W.K., Morlon, H., Ackerly, D.D., Blomberg, S.P., and Webb, C.O. (2010). Picante: R tools for integrating phylogenies and ecology. *Bioinformatics* 26**,** 1463-1464. doi: 10.1093/bioinformatics/btq166.

Kuhn, M. (2008). Building predictive models in R using the caret package. *Journal of statistical software* 28**,** 1-26.

Lahti, L., Shetty, S., Blake, T., and Salojarvi, J. (2017). microbiome R package. *Tools Microbiome Anal R*.

Love, M.I., Huber, W., and Anders, S. (2014). Moderated estimation of fold change and dispersion for RNA-seq data with DESeq2. *Genome Biol* 15**,** 550. doi: 10.1186/s13059-014-0550-8.

Lüdecke, D. (2018). sjPlot: Data visualization for statistics in social science. *R package version* 2.

Lumley, T., and Miller, A. (2009). Leaps: regression subset selection. R package version 2.9. *See http://CRAN. R-project. org/package= leaps*.

Masella, A.P., Bartram, A.K., Truszkowski, J.M., Brown, D.G., and Neufeld, J.D. (2012). PANDAseq: paired-end assembler for illumina sequences. *BMC bioinformatics* 13**,** 31.

Mcmurdie, P.J., and Holmes, S. (2013). phyloseq: an R package for reproducible interactive analysis and graphics of microbiome census data. *PLoS One* 8**,** e61217. doi: 10.1371/journal.pone.0061217.

Nikolenko, S.I., Korobeynikov, A.I., and Alekseyev, M.A. (2013). BayesHammer: Bayesian clustering for error correction in single-cell sequencing. *BMC Genomics* 14 Suppl 1**,** S7. doi: 10.1186/1471-2164-14-S1-S7.

Oksanen, J., Blanchet, F., Kindt, R., Legendre, P., Minchin, P., O'hara, R., Simpson, G., Solymos, P., Stevens, M., and Wagner, H. (2015). "Vegan: Community Ecology Package, R Package version 2.2-1". version 2.2-1 ed.).

Price, M.N., Dehal, P.S., and Arkin, A.P. (2010). FastTree 2--approximately maximum-likelihood trees for large alignments. *PLoS One* 5**,** e9490. doi: 10.1371/journal.pone.0009490.

Rohart, F., Eslami, A., Matigian, N., Bougeard, S., and Le Cao, K.-A. (2017). MINT: a multivariate integrative method to identify reproducible molecular signatures across independent experiments and platforms. *BMC bioinformatics* 18**,** 128.

Schirmer, M., Ijaz, U.Z., D'amore, R., Hall, N., Sloan, W.T., and Quince, C. (2015). Insight into biases and sequencing errors for amplicon sequencing with the Illumina MiSeq platform. *Nucleic Acids Res* 43**,** e37. doi: 10.1093/nar/gku1341.

Shetty, S.A., Hugenholtz, F., Lahti, L., Smidt, H., and De Vos, W.M. (2017). Intestinal microbiome landscaping: insight in community assemblage and implications for microbial modulation strategies. *FEMS microbiology reviews* 41**,** 182-199.

Taylor, M. (2014). *sinkr: A collection of functions featured on the blog 'me nugget'* [Online]. Available: https://github.com/menugget/sinkr [Accessed].
