## Supplementary S13-Parameters Measured for "Impact of industrial production system parameters on chicken microbiomes: mechanisms to improve performance and reduce *Campylobacter*"

**Explanatory variables considered in the model are given below (categorical highlighted as yellow):**

| **Parameter** | **Description** |
| --- | --- |
| *Production_System_N* | Chickens were reared from production system N - Standard |
| *Production_System_HW* | Chickens were reared from production system HW – High Welfare |
| *Production_System_O* | Chickens were reared from production system O – Omega & High Welfare |
| Day_7 | Samples were obtained from Day 7 |
| Day_30 | Samples were obtained from Day 30 |
| No_of_Parent_Flocks_Used_1 | These represent samples from flocks that had one parent flock |
| No_of_Parent_Flocks_Used_2 | These represent samples from flocks that had two parent flock |
| No_of_Parent_Flocks_Used_3 | These represent samples from flocks that had three parent flock |
| Birds_Placed | Number of birds placed in a poultry house |
| Birds_Killed | Number of birds killed at the factory |
| Placement_Birds_per_m2 | Denotes stocking density |
| Feed_Change_Age_Grower_to_Finisher_20 | These represent samples from chickens where the feed was changed from grower to finisher on day 20 |
| Feed_Change_Age_Grower_to_Finisher_22 | These represent samples from chickens where the feed was changed from grower to finisher on day 22 |
| Feed_Change_Age_Grower_to_Finisher_23 | These represent samples from chickens where the feed was changed from grower to finisher on day 23 |
| Feed_Change_Age_Finisher_to_Withdrawal_30 | These represent samples from chickens where the feed was changed from finisher to withdrawal on day 30 |
| Feed_Change_Age_Finisher_to_Withdrawal_31 | These represent samples from chickens where the feed was changed from finisher to withdrawal on day 31 |
| Feed_Change_Age_Finisher_to_Withdrawal_34 | These represent samples from chickens where the feed was changed from finisher to withdrawal on day 34 |
| Weight_Gain_per_Day | This represents the weight gained per day |
| Feed_Conversion_Ratio | This represents the feed conversion ratio |
| EPEF | European Poultry Efficiency Factor |
| Total_Mortality_percentage | Percentage of birds which died during the growing period |
| Total_Leg_Culls_percentage | Percentage of birds which were culled with leg issues during the growth period |
| Pododermatitis_percentage | Percentage of birds with pododermatitis |
| Hockmark_percentage | Percentage of birds with hockmarks |
| PMI_Rejects_percentage | Percentage of birds rejected at post mortem inspection |
| Avg_Weight_at_Slaughter | Average weight at slaughter |
| Age_At_Thin | Age at thinning |
| Age_at_Total_Depopulation | Age at point of total depopulation for the poultry house |
| Total_Water_Consumption | Total water consumed during growing period (flock level) |
| Water_Consumption_per_Bird | Total water consumed during growing period (per bird) |
| log_CFU_per_g_Campylobacter | Log CFU per gram of Campylobacter |
| Protein_perc_ration | Protein percentage within ration |
| Energy_of_Ration | Energy content |
